# CHA1: A New Combinatorial Therapy That Reciprocally Regulates Wnt and JAK/STAT/Interferon Signaling to Re-program Breast Tumors and the Tumor-Resident Landscape

**DOI:** 10.1101/2022.03.25.485706

**Authors:** Mariam K. Alamoudi, Mollie Chipman, Francesca Deieso-Frechette, Ahlam Mukhtar Bogis, Roaya S. Alqurashi, Kaiqi Li, Rui Zhang, Maricel Castañer, George Triafallou, Christopher G. Herbosa, Corinne Carland, AJ. Jaehoon Lee, Kyle Gillani, K. Eric Paulson, Amy S. Yee

## Abstract

Triple negative breast cancers (TNBC) pose exceptional challenges with fatal brain metastases as a clear and unmet need. Immune checkpoint inhibitors (ICIs) are promising therapeutic strategies, but most TNBC are resistant, or “cold” tumors, due to lack of tumor-resident immune engagement. No FDA-approved therapies exist which promote a “cold-to-hot” transition or induce the important biomarker PD-L1, often used for ICI clinical decision-making. Maximal ICI susceptibility, or a full “cold-to-hot” transition, requires reciprocal Wnt signaling inhibition and Jak/STAT/interferon signaling activation. We report a new compound combination (CHA1) that fits the above criteria. CHA1 is comprised of EGCG (epigallocatechin-3-gallate; green-tea compound) and decitabine (DNA-methyltransferase (DNMT1) inhibitor; 5’deaza-cytidine; FDA-approved for hematologic malignancies). We used immune-compromised and syngeneic TNBC pre-clinical models to investigate tumor-intrinsic and tumor-resident T-cell effects, respectively. All results required CHA1 (but not EGCG or decitabine alone) and utilized attainable human dose equivalences with manageable safety profiles. CHA1 triggered efficient Wnt signaling inhibition by elevating Wnt pathway inhibitors (HBP1 and SFRP1) and traversed the blood-brain barrier to reduce both tumor and brain metastatic growth. Transcriptomic and expression analyses revealed that CHA1 treatment effectuated a robust tumor-intrinsic JAK/STAT/IFN response 1) to induce PDL1 and 2) to induce antigen presentation and processing genes, including MHC-1, MHC-2 and numerous genes attributed to professional antigen-presenting cells; 3) to induce CD8+-T-cell infiltration and activation. Additionally, CHA1 pre-treatment improved anti-PDL1 efficacy in a syngeneic setting. Lastly, we derived a composite gene signature emblematic of CHA1 treatment and of a favorable clinical prognosis in-silico. Together, our work supports a model in which CHA1 influences epigenetics, Wnt and Jak/STAT/IFN signaling mechanisms—all to reprogram an epithelial-mesenchymal TNBC tumor to express antigen-presenting properties and to recruit and activate tumor-resident CD8^+^-T-cells. We discuss our findings in the context of cancer biology and immunity with implications for improving ICI susceptibility for TNBC.

## Introduction

About 1 in 8 American women will be diagnosed with breast cancer in her lifetime with higher global rates. While the prognosis is more favorable for localized disease, about 5% of patients present with distant disease and 30% of patients diagnosed with breast cancer proceed to metastatic breast cancer (Stage 4 disease). While earlier staged cancer has a better outcome, metastatic disease accounts for most deaths with an overall survival rate of just 27% (www.breastcancer.org). Triple-negative breast cancer (TNBC) accounts for about 10-15% of all breast cancers and is defined by the absence of therapeutically actionable markers, such as a lack of estrogen or progesterone receptors and of HER2 overexpression. TNBC has a worse prognosis with limited treatment options and has exceptional metastatic potential (reviewed (1, 2)). Breast cancer typically metastasizes to bone, lung, liver, and brain. Advances in the management of non-cranial breast cancer have improved survival, but the ironic consequence of improved management is the increasing prevalence of fatal brain metastases, which typically develop later in the cancer course. Previously, patients were succumbing to non-cranial metastases, but improved treatments have led to the prevalence of fatal brain metastases. The challenge for brain metastases is that most conventional therapies do not cross the blood-brain barrier (reviewed (3–5)). Thus, developing better strategies for metastatic breast cancer and for otherwise fatal brain metastases may greatly improve patient outcome and quality of life. More effective treatments for brain metastases for breast and other cancers represent a critical step improving patient outcomes.

Immune checkpoint inhibitors (ICIs) have revolutionized certain cancer treatments by engaging and activating the local immune environment to destroy tumors (reviewed (6, 7)). Notably, cancers propagate in an immune-suppressed local environment, mediated largely by PD-L1-PD-1 cell-cell interactions between tumor and T-cells in the local environment. The prime cells are the CD8^+^ cytotoxic T-cells that mediate local tumor destruction. The common ICIs (anti-PD-L1 (atezolizumab, avelumab) and anti-PD-1 (e.g., pembrolizumab (Keytruda)) disrupt inhibitory PD-L1-PD-1 interactions between tumor and T-cells, respectively, thus enhancing immune-mediated tumor destruction by promoting local T-cell activation. A third class, anti-CTLA-4, acts through dendritic cells in an inhibitory fashion, also promoting activation of cytotoxic CD8^+^ T-cells in the local environment. The first cancers to be treated were melanomas and lung cancer and with initial dramatic successes. Lung cancer represents an example with excellent criteria for the application of anti-PD-1. Generally, PD-L1 positivity in over 50% of the tumor is a good clinical biomarker for anti-PD1 efficacy ((8–13) reviewed in (14)).

ICI treatment successes are tempered by the sobering fact that only about 25% of tumor types are truly susceptible to ICIs, but most tumors have varying efficacy (e.g.,(8, 15)). Thus, an important goal is expanding the range of susceptible tumors that give efficacious responses to ICIs, while minimizing side-effects, often due to auto-immune reactions. The endowment of tumor susceptibility is commonly referred to as a cold-to-hot transition in which “cold” and “hot” generally refer to low and high inflammation or immune cell infiltration, respectively. The cold-to-hot susceptibility has been extensively discussed, but is a collection of biologically diverse and seemingly unrelated set of properties such as epigenetic statis, β-catenin status, epithelial character, etc., e.g (15)). To date, no FDA-approved therapies exist to induce PD-L1 and engage local immune cell infiltration. Any therapy that changes “cold” to “hot” tumors (15) should theoretically expand the tumor repertoire that can be efficaciously treated by ICIs and thus improve patient outcomes.

TNBC were amongst the first breast cancer subtypes to be treated by ICIs, but there is room for improvement (16, 17). Breast cancers were generally considered cold tumors, as they often lack immune cell infiltration and other molecular signaling characteristics of ICI-susceptible tumors. Because breast cancers have high prevalence, the efficacious use of ICIs can make an impact. In March 2019, anti-PD-L1 received accelerated approval to be used in recurrent or unresectable PD-L1 positive metastatic TNBC, based upon a landmark clinical trial (18). However, a closer inspection of the tumor data showed that those TNBC tumors with elevated PD-L1 expression appeared to better responses, so PD-L1 tests were approved alongside anti-PDL1. Despite verifying PD-L1 positivity, the sobering reality is that Roche voluntarily withdrew anti-PDL1 (under the name Atezolizumab) for TNBC in August 2021 (press release https://www.roche.com/media/releases/med-cor-2021-08-27.htm) due to little improvement to overall survival (19). Recently, anti-PD-1 has also been approved for use in TNBC, with improved outcomes in a neoadjuvant setting (17, 20), but initially as frontline treatment for metastatic TNBC (21, 22). Post-hoc analysis showed that TNBC patients with elevated PD-L1 expression also showed a better outcome. Subsequently, FDA granted full approval to anti-PD-1 in metastatic TNBC whose tumors expressed PD-L1 (20–22). These studies with TNBC highlight the need for developing criteria to improve ICI application and for collaborative therapies that elevate PD-L1 to improve TNBC and breast cancer treatment (22). The expression of PD-L1 in the target tumor may be necessary, but not sufficient to predict future efficacy of anti-PD-1 or anti-PD-L1. More knowledge will be required.

Notable impediments to improved ICI efficacy in tumors are excessive Wnt signaling and dysfunctional JAK/STAT/IFN signaling. A cellular feature of ICI-resistant tumors is a dearth of activated cytotoxic CD8^+^ T-cells in local tumor environment (e.g., rev. in (6, 7, 15)). Many ICI-resistant tumors have elevated Wnt signaling and dysfunctional JAK/STAT/IFN signaling. Thus, a better understanding and new tools aimed at these pathways are necessary to increase tumor susceptibility, but TNBC and resistant tumors have challenges. Most TNBC tumors have hyperactive Wnt signaling (23) that is often associated with excessive breast cancer growth, recurrence, and metastases, including fatal brain tumors (3, 5, 24, 25). There is significant interest for ICIs in treating brain metastases (26, 27). The signature increase in β-catenin levels as part of high Wnt signaling phenotypes contributes to the poor prognosis of patients with TNBC (28). Wnt signaling is a fundamental pathway associated with stem cell renewal and likely contributes to the proliferation of the cancer stem cell population, long linked to metastatic spread and recurrence. Recently, high Wnt signaling in the tumor is correlated to the immune microenvironment in breast and other cancers, including immune evasion and promoting an immunosuppressive environment. Spranger, Gajewski and colleagues have established that high Wnt signaling in numerous cancers is part of immune surveillance mechanisms that prevent decrease T-cell infiltration, a process that is critical for ICIs efficacy and immune-mediated destruction (6, 29–32). Thus, better ways to limit Wnt signaling are necessary. Existing clinical trials with Wnt pathway inhibitors (Wnt 974, PORCN) are ongoing but have revealed some adverse events, including bone fractures (33). Likely, Some Wnt signaling is needed for bone regeneration, but is also important for normal tissue maintenance and regeneration.

A dysfunctional JAK/STAT interferon (IFN) pathway is a major factor in ICIs resistance (6, 7, 33). IFN signaling generally promotes innate and adaptive immune response (34) in T-cells and professional antigen presenting cells (APCs). In addition, active IFN signaling is required for proper anti-tumor immune response, immune checkpoints inhibitors efficacy and for activation of tumor antigen presentation (6). Mutations JAK genes were linked to resistant to anti-CTLA-4 or anti-PD-1 in metastatic melanoma patients, underscoring the requirement for a functional IFN signaling pathway (35). In addition, JAK/STAT/IFN signaling is the major factor in regulating the expression of the important marker and checkpoint gene PD-L1. Using an unbiased shRNA screen with a PD-L1 promoter-reporter, Ribas and colleagues identified 33 genes in the IFN signaling pathway that regulated PD-L1 gene expression, underscoring the importance (36). These two beautiful studies alone highlight the essential role of JAK/STAT/IFN signaling to ICI susceptibility. One of the primary functional consequences of JAK/STAT/IFN signaling is triggering the expression of MHCs (major histocompatibility complex-I and -II, and antigen presentation proteins and processing machinery. In the context of tumor immunity, defects in antigen presentation and processing result in low tumor immunity (reviewed (7)). Both appear to be prognostic factors for favorable outcomes. Numerous studies have correlated defects to in MHC processing and cell surface appearance to defects in tumor immunity and that the proper expression of MHC and antigen processing presentation genes may be a principal output of tumor JAK/STAT/IFN signaling (reviewed in (7)). In the context of a usually epithelial-like tumor, the expression of MHC and antigen-presentation machinery is antithetical on a first glance but may speak to an unusual feature of tumor biology. At the minimum, a robust tumor intrinsic MHC response should recruit tumor-resident cytotoxic CD8^+^ T-cells, NK cells and other cells (reviewed (7)).

In fact, the unusual tumor intrinsic MHC and antigen presentation properties can be considered a “readout” of tumor JAK/STAT/IFN signaling. Still, a necessary requirement in the activation of the tumor-resident CD8^+^ T-cells to achieve maximal tumor killing and maximal ICIs responses. A recent chemical screen has used the appearance of tumor APC properties in the tumor and activated CD8^+^ T-cells as novel criteria in a chemical screen to identify new compounds that endow both necessary properties. The lead compounds would be predicted to globally disrupt epigenetic status through histone modifications (37). Several studies have identified biologically diverse signals that nonetheless and surprisingly intersect epigenetic mechanisms through a viral mimicry response to generate interferon (38–46). The work described here in the context of these studies support and elaborate fundamental mechanisms and the growing role of epigenetic regulation and machinery in defining tumor immunity and improving ICIs efficacy (47–49).

In this paper, we have discovered a compound combination (denoted CHA1) that inhibits Wnt signaling and promotes the cold-to-hot transition in TNBC tumors. CHA1 consists of EGCG (epigallocatechin-3-gallate; principal compound in green tea) and decitabine (an FDA-approved DNA methyltransferase (DNMT1) inhibitor for hematologic malignancies). 1) <u>EGCG.</u> Green tea consumption has been associated with numerous health benefits (50). The green tea compound epigallocatechin gallate (EGCG) is the main catechin component in dry green tea (about 30%). Green tea at therapeutic doses (4∼8 U.S. cups/day) has been tested in humans with no appreciable side effects (51–53) and is in numerous clinical trials for breast cancer and other diseases (clinicaltrials.gov; e.g.(54–58)). We showed that EGCG blocked Wnt signaling through induction of the HBP1 transcriptional repressor to decrease cellular processes relevant to invasive breast cancer (59). Our studies show that EGCG elevation of HBP1 also results in elevation of sFRP1, a Wnt signaling inhibitor. Thus, our work and other animal studies(60–63) show that EGCG blocks constitutive Wnt signaling, perhaps by multiple paths including elevation of Wnt pathway inhibitors HBP1 and sFRP1. 2) <u>Decitabine (</u>2’-deoxy-5-azacytidine; DAC) is an established epigenetics-based agent whose principal action is to inhibit DNA methyltransferases (DNMTs) and induce gene hypomethylation (64). As noted, DAC is FDA-approved for the treatment of myelodysplastic disorders and is used to treat various leukemias and has modest side effects including nausea, neutropenia, and myelosuppression. Both EGCG and Decitabine also has bioavailability to the brain in preclinical studies (65–68). Neither compound alone has had dramatic efficacy in solid tumors (69, 70).

An important challenge for TNBC is discovering strategies to convert tumors from “cold” to “hot”’ or to engage the local immune environment and increase tumor responsiveness to ICIs. As described above, Triple negative breast cancer (TNBC) has limited treatment options with uneven ICIs efficacy. As stated above, maximal ICIs efficacy will require an inhibition of Wnt signaling with an activation of JAK/STAT/IFN signaling. In this paper, we provide evidence that CHA1 treated tumors exhibit properties of a cold-to-hot transition and provides robust induction of PD-L1, a critical biomarker for clinical ICI decision-making. We additionally used CHA1 at equivalent doses attainable in humans to maximize future clinical trial applications. A remarkable feature is that CHA1 orchestrates a re-programming of the epithelial tumor cell to confer antigen-presentation properties that then initiate the local immune interactions. We began with a goal of attenuating TNBC and fatal brain metastases (e.g. (71, 72)), reasoning that Wnt signaling blockade would be efficient with CHA1 because of its induction of Wnt pathway inhibitors through epigenetic and other means and the brain bioavailability of both components. We used human MDA-MB-231 xenografts in immune-compromised mice and found that CHA1 was quite effective on the tumors and the brain metastases—underscoring that CHA 1 therapeutically penetrates the blood brain barrier. Biochemical analyses underscored the efficient inhibition of Wnt signaling through the induction of Wnt pathway inhibitors HBP1 and sFRP1. A notable feature is that we initially used the human TNBC xenografts in mice, but a human RNA-seq with human gene tools. This allowed a snapshot into the intrinsic human tumor changes, initially without a complicating immune system. In a surprise, an unbiased transcriptomic analysis and extensive subsequent bioinformatic and experimental verification revealed that CHA1 treatment effectuated a cold-to-hot transition and additionally a robust JAK/STAT/IFN response. From the biological point of view, CHA1 treatment, but not each component alone, resulted in an inhibition of Wnt signaling and an activation of JAK/STAT/IFN signaling. The intrinsic tumor changes featured a robust induction of MHCs and numerous genes involved in antigen presentation and processing. Both MHC-I and MHC-II could be detected by staining after treatment and with many genes usually attributes to professional antigen-presenting cells. The RNA-seq and much subsequent analysis revealed a robust tumor reprogramming with the acquisition of antigen-presentation (APC) properties and machinery normally found in APCs such as macrophages and dendritic cells. This antigen presentation was accompanied by robust immune cell infiltration in corroborating syngeneic settings, likely through epigenetic disruption and a viral mimicry response. Furthermore, CHA1 not only induces properties to promote increases in tumor-resident CD8^+^ T-cells, but these cells were activated and proliferating. Finally, CHA1 pre-treatment increases the efficacy of anti-PD-L1 in a syngeneic TNBC models. Centered around PD-L1, we also identify a useful hybrid gene signature of future predictions of prognosis and for assessing future compounds with a unique properties of inducing antigen presentation and T-cell activation. These extensive studies constitute a story of CHA1 as an epigenetic disruptor which reprograms an otherwise epithelial-mesenchymal TNBC tumor cells to have antigen presenting properties that allow recruitment and activation of tumor-resident CD8^+^ T-cells and other cells. As discussed, the studies with CHA1 reveal new implications for tumor biology and the tumor-resident environment, all with potential applications in expanding the susceptibility to the new immunotherapies.

## RESULTS

### The design rationale for the CHA1 compound combination

As discussed, Wnt signaling is either a fundamental driver, or involved in progression of many cancers (73, 74). However, clinical trial results from targeted inhibition of Wnt signaling have been mixed due to patient side-effects such as bone fractures, but may have some promise (33). We therefore considered alternatives to targeted inhibition with a hypothesis that embraced our previous findings and known information on epigenetic regulation of Wnt pathway inhibitors such as sFRP1 and HBP1. Further, the role of Wnt-signaling in tumor-intrinsic immune resistance (reviewed in (7)) suggested that an epigenetic strategy to inhibit Wnt signaling may have more far-ranging and significant outcomes (discovered in hindsight).

We hypothesized and extensively tested a compound combination of EGCG and decitabine (2’-deoxy-5-azacytidine (DAC))—to be denoted as CHA1. Our own work in cultured cells highlighted that the green tea compound EGCG inhibited Wnt signaling in TNBC cells (MDA-MB-231) by inducing and stabilizing the HBP1 transcriptional repressor, which we had discovered as an inhibitor of Wnt signaling (59, 75). HBP1 inhibits Wnt signaling by inhibiting the LEF/TCF transcription factors (75, 76). Additionally, the genes for one or more of the Wnt pathway inhibitors such as sFRP1 and HBP1 are themselves subject to suppression by DNA methylation (77, 78). Thus, we sought to effectively increase the levels of inhibitors such sFRP1 and HBP1 (perhaps others) to block Wnt signaling. Numerous studies have shown that decitabine (DAC) is an inhibitor of DNMTs and known to induce sFRP1 by gene DNA demethylation (77, 78). sFRP1 is part of a family of Wnt pathway inhibitors that sequester Wnt ligands away from their cognate FZD receptors, thus blocking pathway activation (79). Our later work also showed that HBP1 was a high affinity repressor for the gene encoding DNMT1, a critical enzyme for enacting maintenance DNA methylation (80).

As noted, neither EGCG nor DAC are targeted molecules and neither compound alone has had dramatic efficacy in solid tumors (69, 81), but if the combination CHA1 had efficacy in TNBC models, the known safety would be a plus for future human applications in clinical trials. Based on a hypothesis centered on the key regulatory points for Wnt signaling, we began a systematic analysis of the CHA1 compounds first in cell culture and then in established pre-clinical models of TNBC. Remarkably, this admittedly simple hypothesis, outlined in Fig 1A, nonetheless inspired systematic experimentation that resulted in the unexpected discovery of a useful biological mechanism/network for tumor cell reprogramming and immunology. Importantly, our unbiased studies also revealed potential to improve and enhance the range of applications for immune checkpoint inhibitors therapies with implications to improve outcomes for breast and other cancers. Finally, the results suggest CHA1 taps a wider molecular network that highlights a complex biological response with clinical implications.

**Figure 1:**
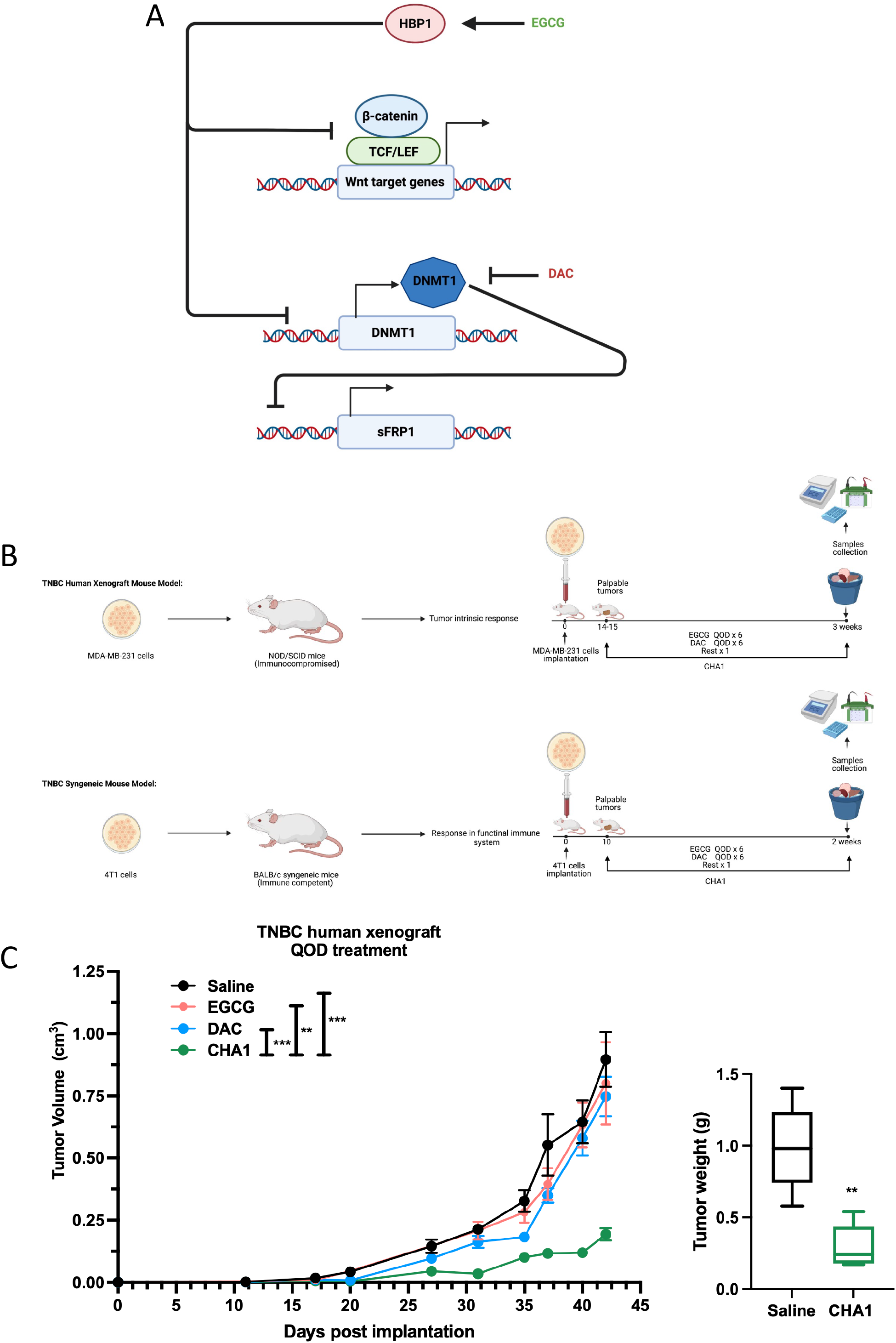
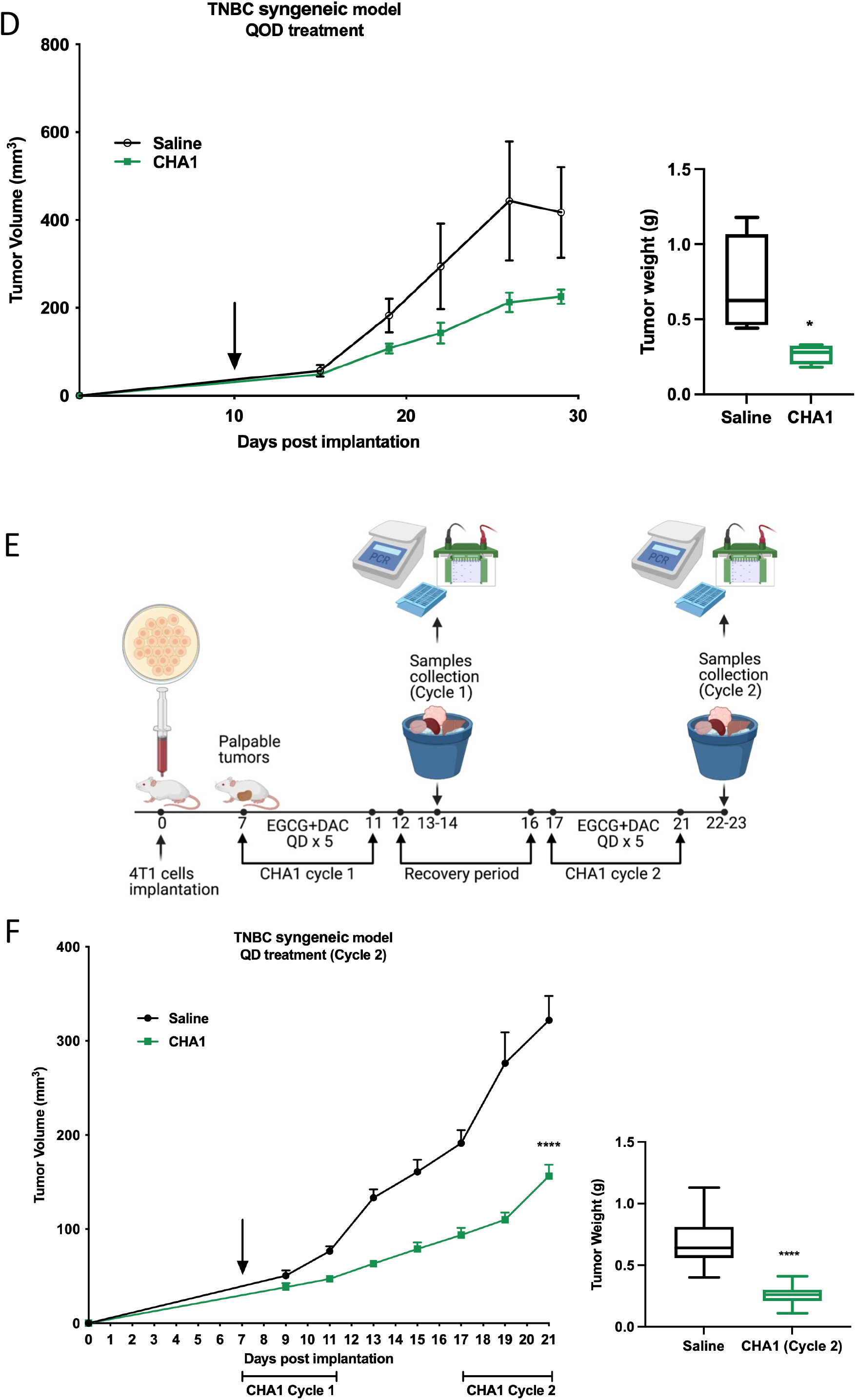
CHA1 suppressed tumor growth in TNBC human xenograft and TNBC syngeneic mouse model. **A)** The proposed mechanism of action of CHA1. CHA1 which is a combinatorial drug approach using EGCG and DAC. EGCG suppress Wnt by post-transcriptional stabilization of mRNA of Wnt inhibitor HBP1. DNMT1 serves as a transcriptional target of HBP1 repression. Targeting epigenetic modification by DAC can reactivate methylated silence gene including the Wnt inhibitor sFRP1. **B)** TNBC human xenograft model using MDA-MB-231 human cells implanted into NOD/SCID mice to study tumor intrinsic response. EGCG and DAC treatment modality (QOD) following detection of palpable tumors is shown. TNBC syngeneic mouse model using 4T1 cells implanted into Balb/c mice to study the CHA1 response in immunocompetent mice. The CHA1 QOD treatment regimen is as follows: EGCG (16.5 mg/kg) or DAC (0.5 mg/kg) was administrated 3 times per week on alternating days for up to 3 weeks in the TNBC human xenograft or up to 2 weeks in the TNBC syngeneic mouse model. **C)** CHA1 QOD treatment significantly suppressed tumor growth while EGCG or DAC monotherapy had no growth inhibitory effect in TNBC human xenograft (n=3-10). The results were combination of two different studies. **D)** CHA1 QOD treatment reduced tumor volume and tumor weight in TNBC syngeneic mouse model (n=4). **E)** CHA1 QD treatment timeline as the following: EGCG (16.5 mg/kg) and DAC (0.5 mg/kg) was administrated concomitantly for two cycles. Each cycle was 5 days with 5 days recovery period. **F)** There was a significant decrease in tumor growth after CHA1 QD cycle 2 treatment in TNBC syngeneic mouse model (n=15-23). The results were combinations of 3 different studies. **C, D, F)** Tumor volume was measured with calipers. Tumors were weighed at the end of the experiment. Mann-Whitney U-test was used for tumor volume and tumor weight comparisons. Fig 1A, 1B, and 1 E were created with BioRender.com. Data were represented as mean ± SEM. * P < 0.05, ** P < 0.01, *** P < 0.005, **** P < 0.001.

### CHA1 reduces tumor growth and metastases in two TNBC mouse models

The proposed mechanism for how EGCG and DAC could together inhibit Wnt signaling (Fig 1A) was bolstered by finding HBP1 expression was inversely correlated with DNMT1 expression in human breast cancer databases (S1A Fig). The initial approach was to test CHA1 and its constituent components in cell culture and then in human xenograft and syngeneic mouse models. As a first step, we treated cultured human TNBC cells (MDA-MB-231) with CHA1 or EGCG monotherapy or DAC monotherapy and by using concentrations that were used previously (up to 1µM DAC, up to 10 µM EGCG). As shown in S1B Fig and S1C Fig, EGCG or DAC monotherapy treatment increased sFRP1 and HBP1 mRNA levels as hypothesized, while the combination was best at increasing both inhibitors. The effect of EGCG on HBP1 gene expression was consistent with what we had previously shown (59). Next, we reasoned that if the induction of Wnt pathway inhibitors was necessary for the CHA1 response, then preventing their increase should diminish the response to CHA1. We used shRNA to knockdown the sFRP1 or HBP1 genes and then assessed the consequence of CHA1 treatment (S1D Fig-S1G Fig). Knockdown of sFRP1 and HBP1 lowered both basal and induced sFRP1 and HBP1 mRNA levels compared to the combination, demonstrating sFRP1 and HBP1 dependence in the CHA1 treatment. These first experiments hinted at the likely importance of HBP1 and SFRP1 in the inhibition mechanism of CHA1 from the Wnt signaling perspective.

We first assessed the efficacy of EGCG plus DAC in an immune compromised human TNBC human xenograft model, which lacks an adaptive immune system. Our initial considerations included safe dosing and eventual translatability to human use. For convenience, the route of administration was (i.p.) for CHA1, ECGC monotherapy, and DAC monotherapy. A detailed description of the dosing equivalencies can be found in (S10 Fig, references within), but in short, the EGCG dose of 16.5 mg/kg i.p. was somewhat less than a dose of 10 mg/kg i.v. which reaches a peak plasma blood level (Cmax) of 0.28 ±0.08 umol/L (86). A similar plasma level can be achieved in humans with oral consumption of between 200 and 400 mg EGCG, which is well below the reported toxic oral dose (51, 87, 88). A DAC dose of 5 mg/kg i.p. in mice is equivalent to a dose of 15 mg/m^2^ in human, which is the lowest of two doses FDA has approved for use in human hematopoietic malignancies (89). Both routes of administration for mice and human results in nanomolar plasma concentrations. Because we were combining the compounds and administering over a longer period of time, we weighted our initial studies towards safety and chose a lower dose of DAC at 0.5 mg/kg i.p. This safety with efficacy rationale proved to be true for our initial experiments with dosing of EGCG or DAC on alternating days (QOD) and for a period of more than 30 days (Fig 1B) (57–69). As shown in Fig 1C <u>and</u> S2A <u>Fig</u>, CHA1 potently inhibited tumor growth in the human TNBC xenograft model, while the individual compounds had little to no effect.

Knockdown of sFRP1 and HBP1 in human TNBC xenograft model promoted aggressiveness and enhanced tumor growth (S2B Fig). Similar to our data in vitro, the expression of sFRP1 or HBP1 was required for CHA1 efficacy in human TNBC xenograft model (S2C Fig-S2H Fig). CHA1’s inhibitory effect on tumor growth was reduced in sFRP1 KD and HBP1 KD human TNBC xenograft model (S2C Fig and, S2D Fig). In addition, CHA1 lost its induction effect on Wnt inhibtors sFRP1 and HBP1 in sFRP1 KD and HBP1 KD human TNBC xenograft model (S2E Fig-S2H Fig). Thus, we conclude that the rise in HBP1 and sFRP1 inhibitors is necessary to the Wnt signaling inhibition upon CHA1 treatment.

Because both components of the CHA1 combination cross the blood brain barrier, we took advantage of the expression of luciferase and GFP in the TNBC xenograft model to examine metastases to brain and other organs. Tumors from treated and untreated mice were surgically resected at 0.35-0.45 mm^3^ and tumor reappearance was monitored by luciferase imaging (S2I and S2J Fig). The appearance of smaller tumors that could not be detected by luciferase were surveyed by GFP imaging (S2J Fig) and multiple small tumors were observed in lung, liver and brain. One mouse from the saline group had seizures and a larger brain metastasis (S2J Fig). CHA1 treatment resulted in reduction of both overall metastases (lymph node, lung, liver and brain) and the subset of brain metastases (S2I Fig and S2J Fig).

We also tested CHA1 in a syngeneic mouse TNBC model (4T1 TNBC cells in Balb/c mice), to confirm anti-tumor effects were not exclusive to the human xenograft model, achieving a similar result of reducing tumor growth using the QOD mode of CHA1 administration (Fig 1D). Consistent with reduced tumor growth and weight, we observed a decreased in cellularity (S2K Fig) and reduction in cell proliferation as measured by Ki67 staining after CHA1 QOD treatment in syngeneic model (S2L Fig). Finally, we altered the delivery modality to better reflect clinical use of DAC, treating tumors in the immunocompetent 4T1 model with both DAC and EGCG together (rather than alternating days) for five-days straight (a cycle, QD), followed by a five-day rest before another CHA1 cycle (Fig 1E). This multi-cycle treatment modality was also effective at slowing or arresting tumor growth (Fig 1F), even after a single QD cycle 1 (S2M Fig).

### CHA1 inhibits Wnt signaling

CHA1 effectively inhibited Wnt signaling in tumors as measured by expression of β-catenin. Each TNBC mouse model treated with CHA1 showed a decrease in β-catenin protein level compared to control group (Fig 2A). The western blot result was confirmed by immunohistochemical (IHC) staining of tumors, which showed a significant decrease in cells expressing β-catenin in CHA1-treated mouse and human TNBC tumors (Fig 2B and S3A Fig). CHA1 treated TNBC tumors in both the immune compromised and immune competent xenograft models showed a significant decrease in AXIN2 mRNA compared to saline (Fig 2C). We also confirmed the requirement for both components of CHA1 to maximally inhibit Wnt signaling by analyzing expression of Wnt target gene AXIN2. In the human xenograft model, EGCG monotherapy had no effect on AXIN2 expression, while DAC monotherapy resulted in a surprising increase in AXIN2 expression (S3B Fig). Supporting the mechanism outlined in Fig 1A, we analyzed the expression of both sFRP1 and HBP1 mRNA in each tumor model. Regardless of treatment modality, both sFRP1 and HBP1 were significantly induced (Fig 2D). Finally, the induction effect of CHA1 was greater than the effect of individual compounds in the TNBC human xenograft model (S3C Fig). These results demonstrated that the CHA1 combination treatment in vivo could inhibit both growth and Wnt signaling in TNBC tumors.

**Figure 2:**
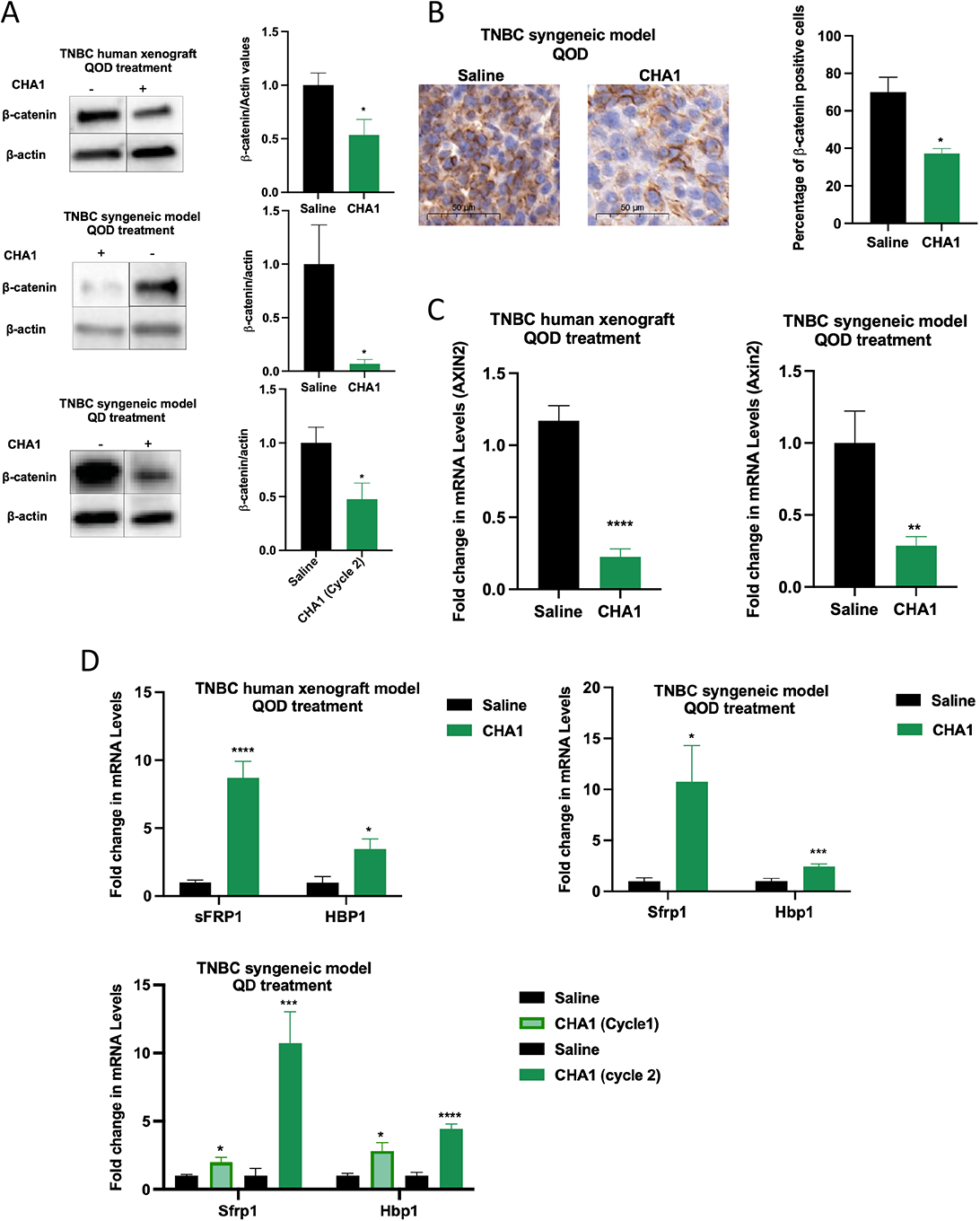
CHA1 treatment suppressed Wnt signaling in TNBC human xenograft and TNBC syngeneic mouse model. **A)** CHA1 significantly decreased the protein levels of β-catenin. Representative western blot (left) with quantification (right) of saline and CHA1 treatment tumors (n=4 mice/ group). **B)** Immunohistochemistry of β-catenin in saline and CHA1 QOD treatment in TNBC syngeneic model. Saline treated tumors demonstrated a strong membrane staining of β-catenin and high number of β-catenin positive cells (average Allred score = 8) compared to CHA1 treated tumors (average Allred score = 5). Different Sections of tumors were stained with anti-β-catenin antibody. Representative immunostaining section of with quantification of the percentage of β-catenin positive cells in saline tumors and CHA1 treated tumors (n= 3 mice/ group). **C)** CHA1 QOD significantly downregulated the gene expression of Wnt target genes AXIN2. AXIN2 gene expression was tested by qRT-PCR after saline and CHA1 treatment (human xenograft QOD, n=6 mice, syngeneic QOD, n=5). **D)** CHA1 significantly induced the gene expression of Wnt inhibitors sFRP1 and HBP1. The gene expression of sFRP1 and HBP1 was measured by qRT-PCR in control and CHA1 treated tumors (human xenograft QOD, n=5-6 mice, syngeneic QOD, n=5-6, syngeneic QD cycle 1, n= 4, syngeneic QD cycle 2, n=6-7). For figures A-D, unpaired two-tailed t-test was used for the comparison between two groups. qRT-PCR experiments were done in triplicates in three independent experiments. Western blot quantification was the combination of two different experiments (n=4/experiment). Data were represented as mean ± SEM. *\* P < 0.05, ** P < 0.01, *** P < 0.005, **** P < 0.001*.

### Unbiased bioinformatic analysis reveals CHA1 treatment re-programs multiple cellular functions: Tumor-immune functions

We sought additional mechanistic insight into the tumor-intrinsic consequences of CHA1-inhibition of Wnt signaling. We used an unbiased human-specific RNA-seq and bioinformatics approach to analyze human tumor xenografts with and without CHA1 treatment. This proved to be a serendipitous choice, since the analysis of human genes provided insight into molecular re-arrangements intrinsic to the tumor and were uncomplicated by a full (mouse) immune system, leading to the discovery of an unusual tumor reprogramming that resulted in enhanced tumor-immune functions. Fig 3A displays the top 20 canonical pathways significantly affected by CHA1 treatment using Ingenuity Pathway Analysis (IPA). As predicted from the results of S1 Fig, S2 Fig, Fig 2, and S3 Fig, Wnt signaling was downregulated. Multiple Wnt targets identified in the RNA-seq were significantly inhibited (Fig 3B and S1 Table), supporting our prior analysis of Wnt inhibition.

**Figure 3:**
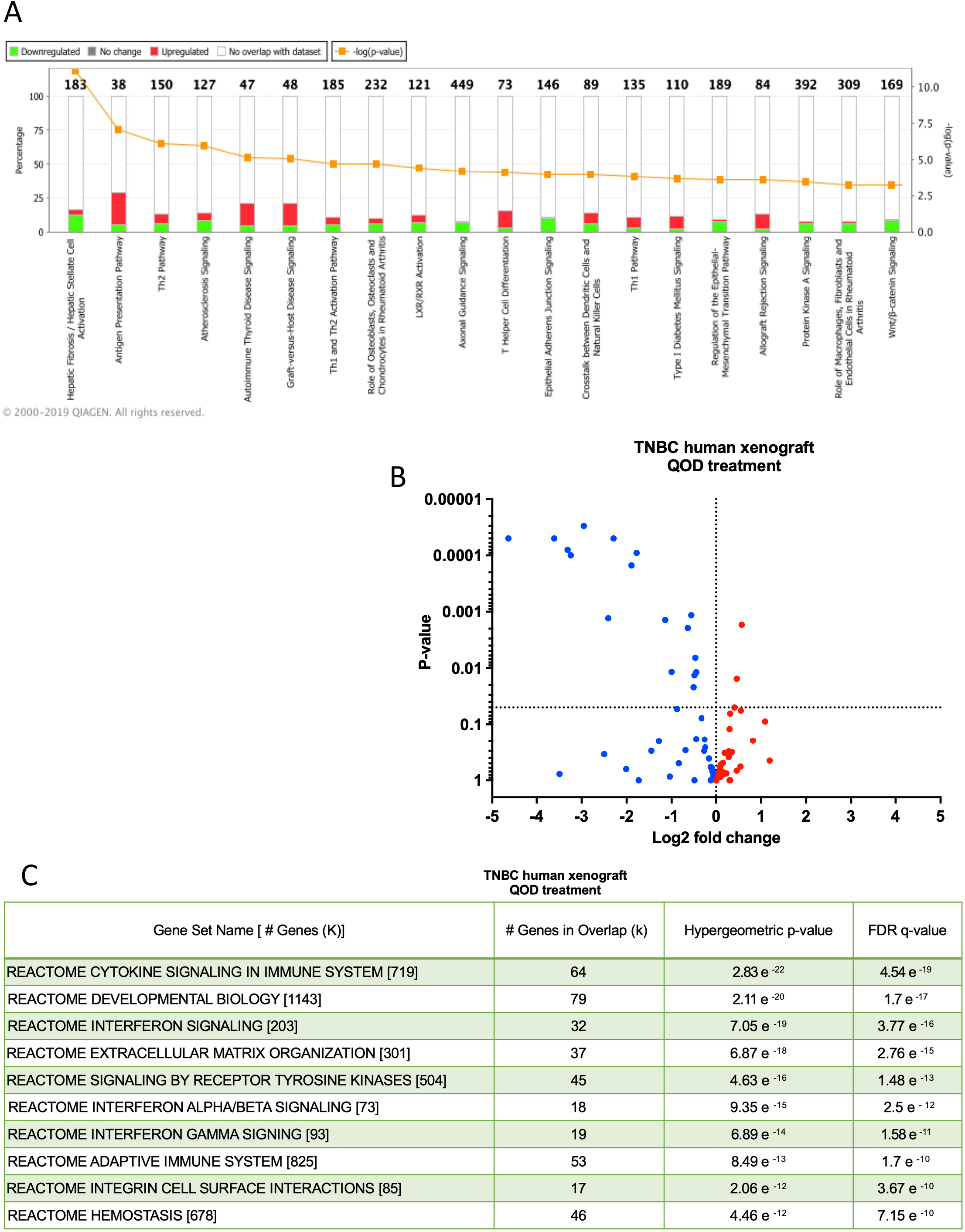
Bioinformatic analysis of CHA1 treated tumor indicated a broad activation of immune-related pathways and downregulation of Wnt pathway in TNBC human xenograft model. Saline and CHA1 treated tumors were subjected to RNA-seq. **A)** The data was analyzed by IPA. The top 20 canonical pathways significantly altered after CHA1 treatment were depicted. There was a broad activation of immune system related signaling pathway e.g., antigen presentation, autoimmune thyroid disease signaling, graft-vs-host disease signaling, Th1 and Th2 activation and allograft rejection signaling. Antigen presentation pathway was activated, indicating upregulation of the genes involved in anti-tumor immune response. Also, Wnt signaling was downregulated. **B)** Volcano plot demonstrated suppression of Wnt signaling pathway after CHA1 QOD treatment. The gene set was extracted from IPA after analysis of CHA1 RNA-seq. 45 out of 83 (54.2%) Wnt target genes were downregulated after CHA1. 17 out of 45 (37.7%) Wnt target genes were downregulated with *P < 0.05* after CHA1. **C)** GSEA MSigDB analysis identifying the REACTOME gene sets of CHA1 treated tumors indicated activation of IFNα/β and IFNγ.

While the suppression of Wnt signaling was gratifyingly confirmed, we found a surprising and broad cellular re-programming, in particular finding multiple pathways involved in immune surveillance activated. These overlapping, tumor-intrinsic gene sets centered around antigen presentation, an unusual property for an epithelial cell. Such properties would not have been easily observed in a similar analysis of a syngeneic tumor model, that is with both a mouse tumor and mouse immune system. As shown in S2 Table-S8 Table, the core set consisted of MHC class I and MHC class II antigen presentation genes. MHC-I and MHC-II are also IFN-stimulated genes. While CHA1 appears to have particularly broad function, others have also tapped into some of these mechanisms through diverse paths such as CDK4/6 inhibition and H3K4 demethylation by an LSD1 inhibitor (40, 90) (see Discussion). As described later, this broad re-programming has important implications for tumor immunology and the efficacy and susceptibility to immune checkpoint inhibitors.

### CHA1 treatment enhances antigen presentation

The bioinformatic analyses strongly suggested CHA1 induces antigen presentation in tumors. As shown in Fig 3A and S2 Table -S8 Table, the core set consisted of MHC-I and MHC-II antigen presentation genes. The bioinformatic analysis highlighted the CHA1 induction of antigen presentation in tumors, in particular the HLA genes comprising the major histocompatibility complexes (MHC) (IPA data, Fig 3A and S2 Table -S8 Table) and the machinery leading to antigen processing and display. MHC-I and MHC-II components are required for proper antigen presentation, including tumor neo-antigens and tumor-associated antigens such as cancer-testis antigens (CTAs)(see below). We used both the human xenograft and the syngeneic mouse model tumors to verify and discriminate the tumor intrinsic responses and as a tool to distinguish CHA1 action in the immune microenvironment.

This RNA-seq data set reliably identified many genes that we later confirmed by qRT-PCR, but we selected some informative genes for a detailed verification. First, confirming the bioinformatics results, MHC-I and MHC-II gene expression in both immune compromised and immune competent TNBC models were induced by CHA1 treatment (Fig 4A). Second, IHC staining demonstrated CHA1 treatment induced MHC-I and MHC-II proteins in human xenograft (Fig 4B). Third, the expression pattern of MHC-II was dependent on CHA1, but not the individual constituents. CHA1 treatment induces expression of MHC-II genes in human xenograft (S4A Fig) but neither EGCG monotherapy nor DAC monotherapy had any remarkable effect alone. Fourth, the induction of MHC-II by CHA1 in human xenograft suggested the induction of tumor-specific MHC-II (tsMHC-II) that appears to promote tumor recognition by the immune system (91). It has been reported that tsMHC-II correlated with favorable response to ICIs in humans, a better prognosis and with tumor rejection in murine models (92–96). Fifth, we also observed increased expression of CD74, MHC class II invariant chain (li), in human xenograft model, that has important role in proper folding and trafficking of MHC class II complex, leading to CD4^+^ helper T-cells activation (97). Sixth, our data showed that the expression of PSMB9 and Psmb10 was significantly elevated after CHA1 in human xenograft and syngeneic models, respectively (Fig 4A). Both PSMB9 and Psmb10 are subunits of immune immunoproteasome that play important role in processing internal antigen and loading on to MHC-I complex (a necessary factor for later activation of CD8^+^ cytotoxic T-cells (98). In summary, the finding of CHA1-induced MHC-I and MHC-II expression in the human xenograft model as well as the number of cells with MHC expression suggests that CHA1 treatment activates a robust antigen presentation in a TNBC tumor with epithelial-mesenchymal characteristics.

**Figure 4:**
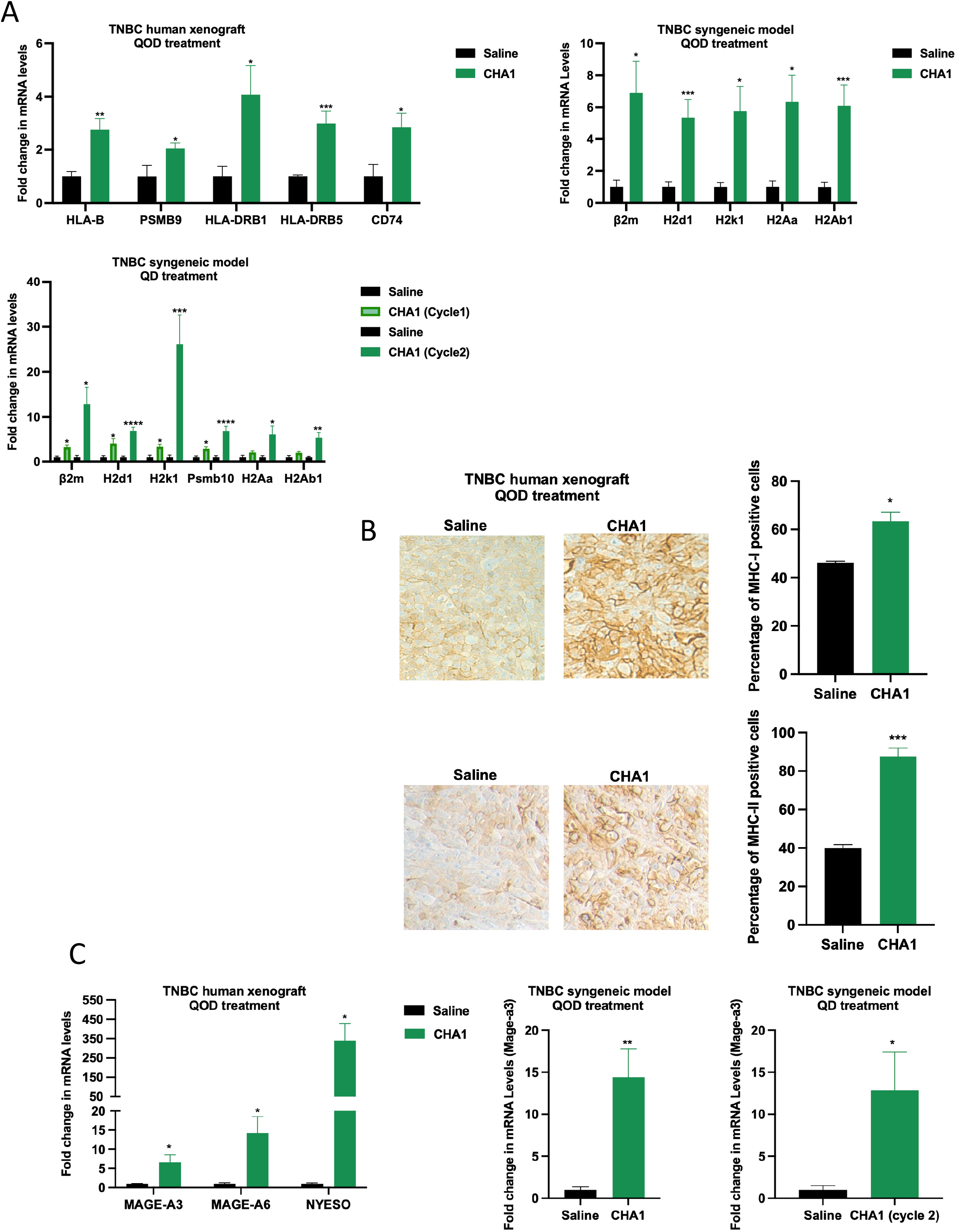
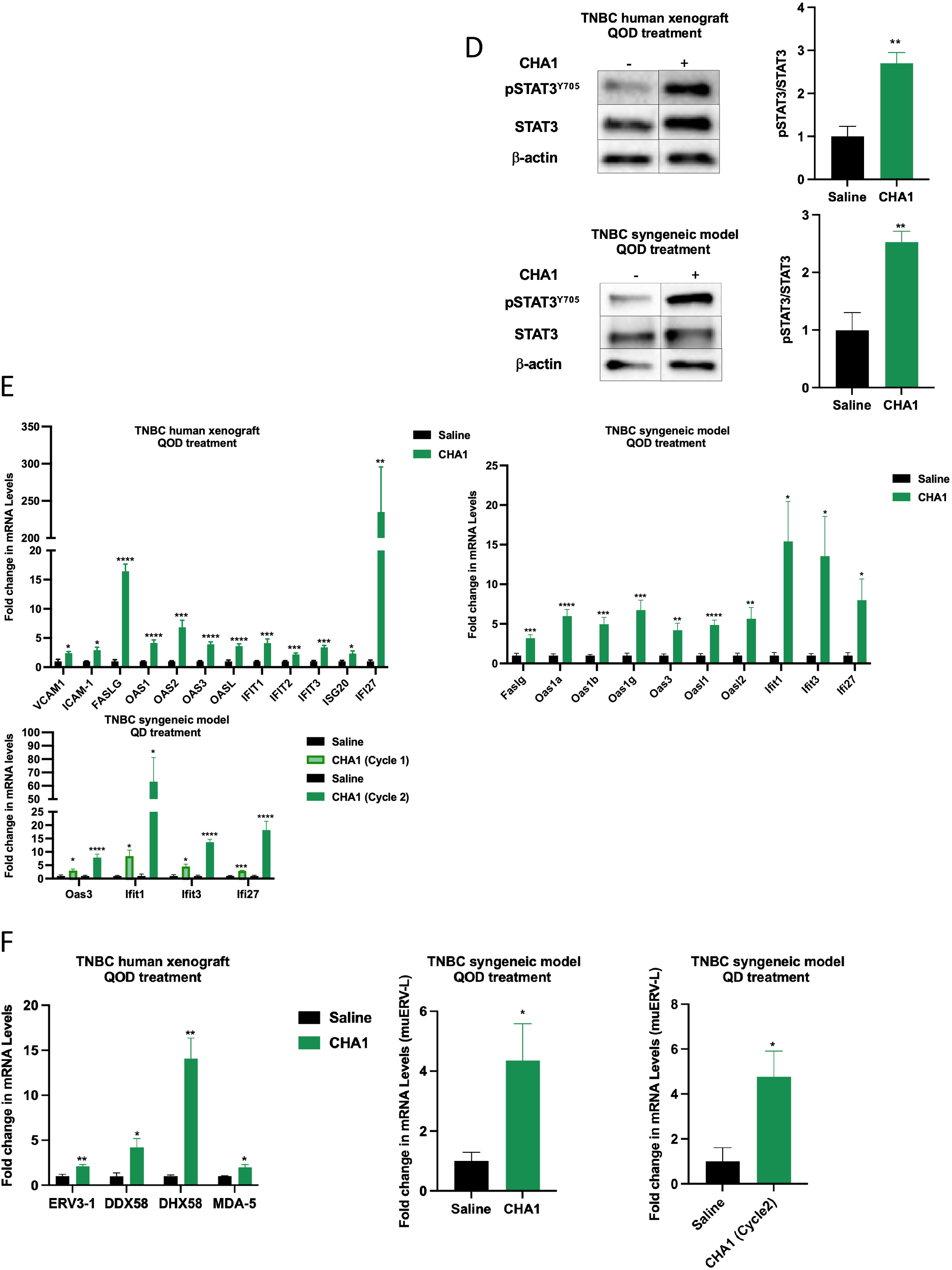
CHA1 treatment activated antigen presentations and stimulated IFNα/β and IFNγ pathways in TNBC human xenograft and TNBC syngeneic mouse model. **A-C)** CHA1 QOD treatment in TNBC human xenograft and TNBC syngeneic mouse model and CHA1 QD treatment in TNBC syngeneic mouse model enhanced antigen presentation. **A)** CHA1 significantly induced the gene expression of MHC-I (HLA-B in TNBC human xenograft, β2m, H2d1 and H2k1 in TNBC syngeneic model), MHC-II (HLA-DRB1 and HLA-DRB5 in TNBC human xenograft, H2Aa and H2Ab in TNBC syngeneic model), immunoproteasome (PSMB9 in TNBC human xenograft and Psmb10 in TNBC syngeneic model) and the invariant chain (CD74 (Ii) in TNBC human xenograft). The gene expression was measured by qRT-PCR in saline and CHA1 treated tumors (human xenograft QOD, n=4-6 mice, syngeneic QOD, n=6, syngeneic QD cycle 1, n=3-4, syngeneic QD cycle 2, n=5-7). **B)** Immunohistochemistry of MHC-I and MHC-II in TNBC human xenograft demonstrated a weak staining of MHC-I and MHC-II in saline group (Average Allred score =4-5) and a strong MHC-I and MHC-II staining in CHA1 QOD treated group (Average Allred score = 8). Different sections of different untreated and treated tumors were stained with pan anti-MHC-I antibody and anti-MHC-II antibody. Representative immunostaining section of MHC-I and MHC-II with quantification of the percentage of MHC-I and MHC-II positive cells in saline tumors and CHA1 QOD treated tumors (n= 3 mice/ group). **C)** CHA1 treatment significantly upregulated the gene expression of CTAs. The gene expression was measured by qRT-PCR in saline and CHA1 treated tumors (human xenograft QOD, n=4-6 mice, syngeneic QOD, n=6, syngeneic QD cycle 2, n=5). **D-F)** CHA1 QOD treatment in TNBC human xenograft and TNBC syngeneic mouse model and CHA1 QD treatment in TNBC syngeneic mouse model stimulated IFN pathways. **D)** CHA1 significantly increased the protein expression of pSTAT3Y705. Representative western blot (left) with quantification (right) of saline and CHA1 QOD treatment tumors in TNBC human xenograft and TNBC syngeneic model (n=4 mice/ group). **E)** CHA1 significantly induced the gene expression of ISGs. The gene expression was measured by qRT-PCR in saline and CHA1 treated tumors (human xenograft QOD, n=4-6 mice, syngeneic QOD, n=5-6, syngeneic QD cycle 1, n=3-4, syngeneic QD cycle 2, n=4-6). **F)** CHA1 activated viral mimicry status. The gene expression of ERV3-1 and dsRNA PRRs was tested in saline and CHA1 QOD treatment in TNBC human xenograft (n=4-6). The gene expression of muERVL-1 was tested in saline and CHA1 QOD treatment (n=5) and CHA1 QD treatment (n=4-6) in TNBC syngeneic model. **A-F)** Unpaired twotailed t-test was used for the comparison. qRT-PCR experiments were done in triplicates in three independent experiments. Western blot quantification was the combination of two different experiments (n=4/experiment). Data were represented as mean ± SEM. *\* P < 0.05, ** P < 0.01, *** P < 0.005, **** P < 0.001*.

The tumor reprogramming to apparent immune cell function was not limited to antigen presentation, usually the purview of T-cells. We also observed and verified the acquisition of natural killer cells and cytotoxic CD8^+^ T-cells functions (99). In the human xenograft model, we verified the mRNAs for human perforin, normally associated with cytotoxic CD8^+^ T-cells and NK cells (S5A Fig). In this same xenograft models, we also noted the appearance of mouse perforin and granzyme, normally attributed to the mouse NK cells, which are presented in the NOD/SCID mice used in the xenograft (100) (S5B Fig). These results suggest that CHA1 engages a mechanism that can also be engaged by many diverse signals and may fundamentally dictate how a tumor engages its local immune environment.

### A broader cellular context to CHA1 reprogramming

In addition to renewed expression of antigen presentation machinery, we observe expression of cancer testis antigens (CTAs). CTAs such as NY-ESO-1 and MAGE-A family are tumor-associated antigens that represent a class of antigens that may increase immune recognition of cancers as a result of their restricted expression in normal adult tissue (101, 102). CHA1 re-activates the expression of NY-ESO-1, MAGE-A3 and MAGE-A6 CTAs in human xenograft and MAGE-A3 in syngeneic models (Fig 4C). A more detailed analysis of CTAs MAGE-A3, MAGE-A6 and NY-ESO-1 with each component (EGCG monotherapy or DAC monotherapy) compared to CHA1 in human xenograft (S4B Fig) shows that EGCG monotherapy has a negligible effect on CTAs expression, while DAC treatment results in increased the expression, consistent with previous report (101). However, the CHA1 combination produces what appeared to be a synergistic response, even compared to the substantial DAC response.

Another change was the reversal of epithelial-to-mesenchymal transitions resulting from CHA1 treatment (S5C Fig and S5D Fig). MDA-MB-231 and most TNBC tumors have more mesenchymal characteristics. Supporting the bioinformatic result, we found that protein for E-cadherin was robustly induced with CHA1 treatment in both mouse models (S5C Fig and S5D Fig). While the E-cadherin gene is known to be methylated in tumors (103) and thus a candidate for epigenetic modification by CHA1, this and other tumor-altering mechanisms potentially regulated by CHA1 treatment may merit further analysis, including decreased in cell survival and cell viability, inhibition of cellular movement, activation of apoptosis and increased necrosis and other tumor-inhibitory pathways (S5E Fig and S9 Table). In summary, CHA1 treatment endows the breast tumor with broad antigen presentation and immune cell properties, but also effects a broad re-programming of an otherwise mesenchymal tumor cell. Upon further investigation, our work appears to intersect with newer studies that indicate that tsMHC-II presence has positive functional properties for tumors.

### CHA1 activates the IFN signaling pathway

We next asked how CHA1 could trigger a robust and strong antigen presentation response. Some evidence supports crosstalk between Wnt and IFN signaling (104–106), therefore we used a second unbiased analysis of the RNA-seq dataset (Gene Set Enrichment Analysis Molecular Signatures REACTOME Database, GSEA MSigDB) to refine the identification of the molecular pathways that are altered upon CHA1 treatment. As shown in Fig 3C, the top-ranked pathways include, immune cytokine signaling, general interferon signaling, interferon α/β signaling, and interferon γ signaling, and adaptive immune system. Notably interferon γ is associated with induction of genes for MHC and antigen presentation (107).

The signature events for interferon α/β signaling and interferon γ signaling are JAK-mediated tyrosine phosphorylation of a STAT transcription factor and the expression of numerous interferons stimulated genes (ISGs). First, we assessed for a possible CHA1 activation of IFN signaling by measuring pSTAT3^Y705^ western blot in CHA1-treated immune compromised and immune competent TNBC tumors (Fig 4D). The use of the human xenograft especially underscored a tumor-intrinsic response to IFN signaling. The requirement for both components of CHA1 for maximal STAT3^Y705^ phosphorylation was again shown in tumors treated with CHA1, or EGCG monotherapy or DAC monotherapy (S4C Fig). Second, we examined the expression of selected genes derived from the overlap between our RNA-seq of CHA1-treated human xenograft mice and the and the IFNα/β and IFNγ GSEA MSigDB REACTOME dataset (S4D Fig). To investigate activation, a hypergeometric probability analysis was conducted, indicating that we required to confirm the induction of at least eight pathway genes for statistical significance (S4D Fig). Fig 4E shows a significant increase in the transcription of ISGs in CHA1-treated immune compromised and immune competent TNBC tumors. According to the analysis of the immune compromised RNA-seq by GSEA MSigDB REACTOME dataset, the HLA-B, OAS1, OAS2, OAS3, and OASL genes overlap between the IFNα/β and IFNγ pathways. IFIT1, IFIT2, IFIT3, ISG20, and IFI27 are IFNα/β-specific target genes while HLA-DRB1, HLA-DRB5, ICAM-1 are VCAM1 are IFNγ-specific target genes. Analysis in the immune compromised TNBC model indicated that CHA1 treatment is required for induction of IFNα/β -and IFNγ stimulated genes. DAC monotherapy treatment resulted in increased the expression of IFNα/β and IFNγ stimulated genes except IFIT1 and IFIT3, however, the gene expression stimulated by CHA1 treatment was greater. EGCG monotherapy treatment had no remarkable induction effect on ISGs (S4E Fig). Lastly, the strength of interferon signaling was highly correlated with ISG expression as shown by plotting pSTAT3^Y703^ levels against OAS1, OAS2, OAS3, OASL, IFIT1, IFIT2 and IFIT3 mRNA level, with r^2^ values between 0.6 and 0.8 (S4F Fig).

We next asked how IFN was being produced. A major clue came from the human xenograft model, which is grown in an immune compromised mouse where immune cells cannot be a source of IFN of any type. We noticed a gene expression pattern consistent with ancient mechanisms known as viral mimicry, which is cellular response to a perceived insult. Previous work has shown that DNMT inhibition can result in re-activation of endogenous retroviruses (ERVs), elevating the level of dsRNA and activating dsRNA pattern recognition receptors such as RIG-1 (DDX58), LGP2 (DHX58), and MDA5 (IFIH1) (108). Indeed, CHA1 induced endogenous retrovirus (ERV3-1) mRNA in the human TNBC xenograft and induced murine endogenous retrovirus like-1 (muERVL-1) in mouse TNBC models (Fig 4F). In addition, CHA1 treatment activated gene expression of dsRNA pattern recognition receptors in the immune compromised model (Fig 4F). As will be expanded in the discussion, the path from inhibition of DNA methylation to viral mimicry, interferon induction and acquisition of antigen presentation in an epithelial cell maybe part of fundamental cellular functions that can be tapped to respond to diverse signals via epigenetic rearrangements (CDK4 inhibition, LSD1, CDK9; see discussion).

### CHA1 treatment, cold-to-hot transition and immune checkpoints: conferring susceptibility to immune checkpoint inhibitors?

We next tried to discern a workable hypothesis for the panoply of changes that include antigen presentation and EMT—all within a breast epithelial tumor/cell. The most plausible framework is the cold-to-hot transition that has been often used to describe the transition from a tumor with an immunosuppressive environment (cold) to a tumor with an immunocompetent environment (hot) (15). The disparate properties of hot tumors include acquisition of antigen-presentation properties, the otherwise unrelated EMT. High Wnt signaling is a characteristic of the cold tumor; has emerged as a major pathway for immune evasion or suppression; and is also associated with EMT (109, 110). High IFN signaling is a necessary part of hot tumor for the induction of APC and other pathways. Numerous studies have highlighted decreased Wnt signaling and increase IFN signaling as a feature of hot tumors with maximal potential for immune destruction and importantly, maximal susceptibility to the class of immune checkpoint inhibitors that enable local immune destruction.

The connection of disparate results from an unbiased analysis to the “cold to hot” transition, or the regulatory mechanisms that specify tumor-immune interactions provided a path and focus to the subsequent work. We hypothesized that CHA1 may be a useful tool to improve efficacy and susceptibility to immune destruction by immune checkpoint inhibitors. The final sections of the Results seek to test this hypothesis and to add greater molecular detail for future clinical applications. While these investigations gave rise to many intriguing findings, a major conclusion of that CHA1 treatment heightens expression of a key clinical biomarker PD-L1 and of signatures of a T-cell inflamed phenotype, both of which often predict efficacy to ICIs treatment.

For the most successful immune checkpoint inhibitor treatments, the tumor often has a combination of conditions that promote treatment success and include the changes that were observed in Fig 1-Fig 4 (15). As shown in Fig 5A, tumors infiltrated with CD8^+^ T-cells, NK cells and macrophages, termed “hot” tumors, are the most responsive to checkpoint inhibitors. In contrast, tumors infiltrated with T_reg_ cells (denoted “cold” tumors) are often less responsive. As demonstrated in Fig 1-Fig 4, in mice with and without an adaptive immune system, we observed that CHA1 treatment results in significant changes that may promote the “hot” tumor phenotype, including reduced Wnt signaling, activated IFN signaling, induction of CTAs and antigen presentation machinery. These features correlated with a dramatic decrease in tumor growth upon CHA1 treatment. Therefore, we sought to test if CHA1-treated tumors in mice with a full immune system acquired some or all the “hot” state characteristics.

**Figure 5:**
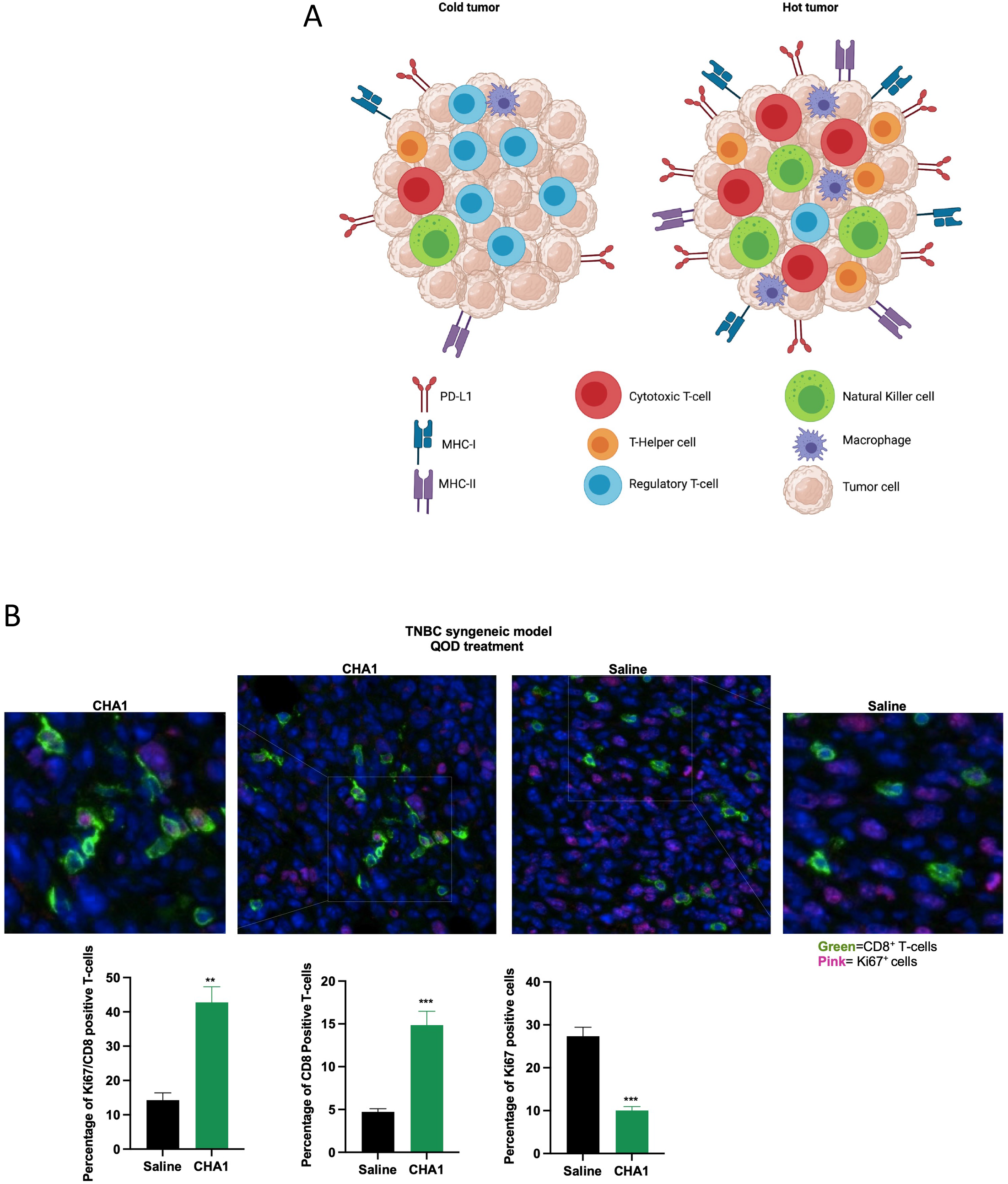
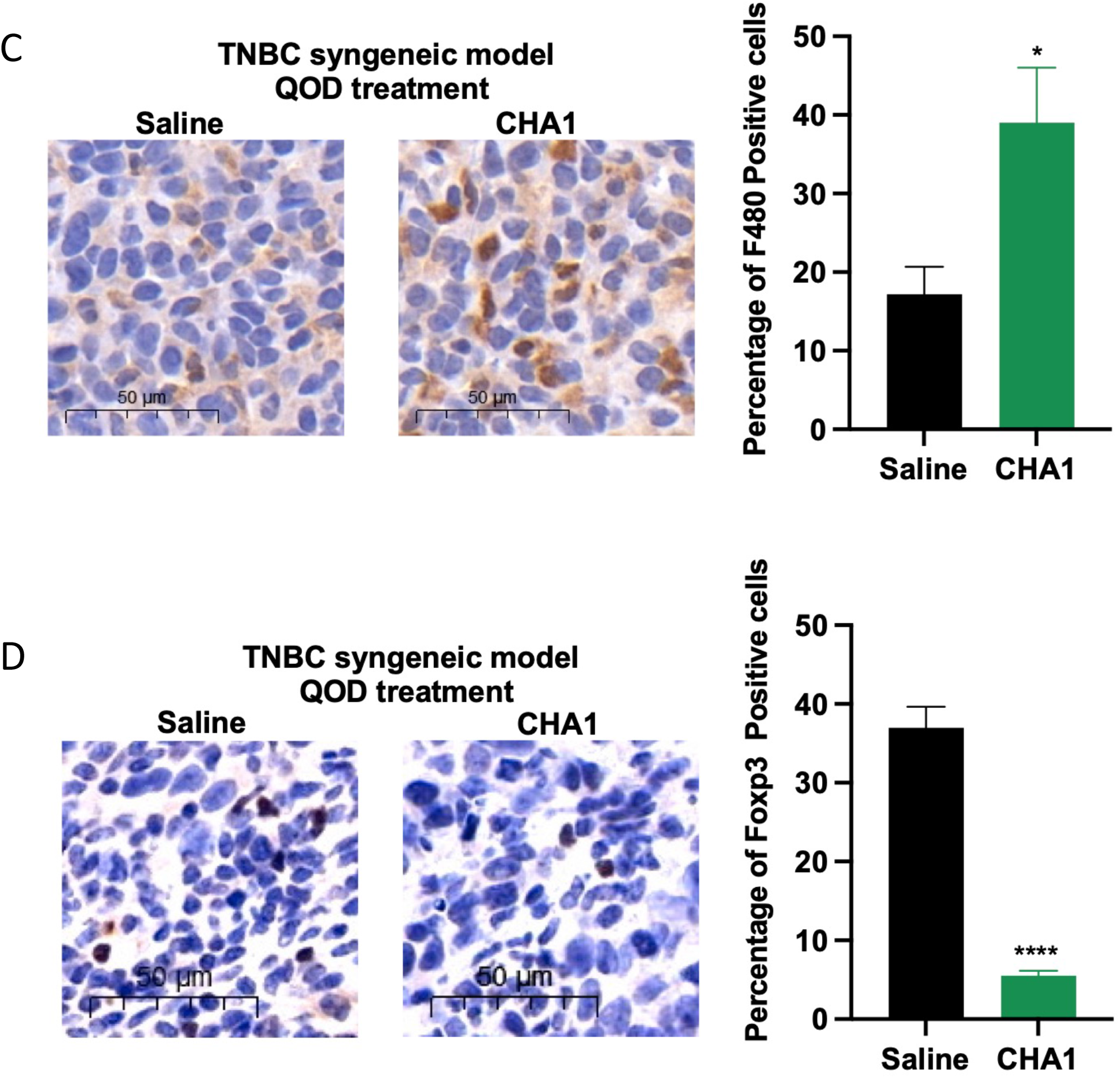
CHA1 treatment increased infiltration of immune-effector cells and decreased infiltration of immune-suppressor cells in TNBC syngeneic mouse model. **A)** The characteristic of hot and cold tumors. Hot tumor is characterized by increased infiltration of immune effector cells such as cytotoxic T-cells, T-helper cells, and natural killer cells and decreased infiltration of immune suppressor cells such as regulatory T-cells. Hot tumors also have high expression of PD-L1 and antigen presentation machinery including MHC-I and MHC-II. Fig 5A was created with BioRender.com. **B, C)** CHA1 QOD treatment increased infiltration of immune effector cells into TME in TNBC syngeneic model. **B)** Immunofluorescent co-staining of Ki67^+^CD8^+^T-cells revealed that CHA1 QOD in TNBC syngeneic model increased infiltration of active proliferating IFNγ-secreting Ki67^+^CD8^+^ cytotoxic T-cells. Different sections of different untreated and treated tumors were stained with anti-CD8 and anti-Ki67 antibody. Representative immunofluorescence staining section of Ki67^+^CD8^+^ T-cells with quantification of the percentage of Ki67^+^CD8^+^ T-cells, CD8^+^ T-cells and Ki67^+^ tumor cells in control tumors and CHA1 QOD treated tumors (n= 4 mice/group). **C)** Immunohistochemistry of F480^+^ cells indicated increased infiltration of macrophages after CHA1 QOD in TNBC syngeneic model. Different sections of different untreated and treated tumors were stained with anti-F480 antibody. Representative immunostaining section with quantification of the percentage of F480^+^ cells in control tumors and CHA1 QOD treated tumors (n= 5 mice/group). **D)** CHA1 treatment decreased infiltration of immune-suppressor cells into TME in TNBC syngeneic mouse model. Immunohistochemistry of FOXP3^+^ T-cells indicated decreased infiltration of Treg after CHA1 QOD in TNBC syngeneic model. Different sections of different untreated and treated tumors were stained with anti-FOXP3 antibody. Representative immunostaining section with quantification of the percentage of FOXP3^+^ T-cells in control tumors and CHA1 QOD treated tumors (n= 4 mice/group). **A-C)** Unpaired two-tailed t-test was used for the comparison between two groups. Data were represented as mean ± SEM. * *P* < 0.05, ** *P* < 0.01, *** *P* < 0.005, **** *P* < 0.001.

Extending the mechanism of CHA1 function to the tumor immune environment, we first, examined infiltration of T-cells by staining for CD8^+^ T-cells in CHA1-treated and untreated tumors. As shown in Fig 5A and S6D Fig, the number of CD8^+^ T-cells was significantly higher in CHA1-treated tumors. Moreover, CHA1 treatment increased the percentage of Ki67^+^CD8^+^ T-cells, indicating that CHA1 treatment recruited active proliferating cytotoxic T-cells that secrete IFNγ (Fig 5A) (111). In contrast, Ki67 staining in the tumor was reduced with CHA1 treatment. Thus, CHA1 or the changes induced in the tumor may promote T-cell activation. To further assess the immune cells in the tumor microenvironment, macrophages were stained using F480 as a marker. As shown in Fig 5B, elevated F480^+^ cells were observed after CHA1 treatment. In addition, we examined whether there was a decrease in FOXP3-positive T_reg_ cells. As shown in Fig 5C, FOXP3 staining declined dramatically in CHA1-treated tumors. Lastly, returning to the human xenograft model where we initially defined tumor intrinsic changes, we noticed the induction of mRNA for mouse Prf1 and Gzmb (S5B Fig). NOD/SCID mice lack B-and T-cells but are known to have residual NK cell function (100) suggesting that the human tumor may recruit the mouse NK cells. Regardless of model, the tumor changes upon CHA1 treatment promotes recruitment of local immune cells—whether the necessary CD8^+^ T-cell, macrophage infiltration, while excluding FOXP3^+^ T_reg_ cells. Even the human xenograft appears to recruit the vestigial immune system in the immune-compromised setting. These results are consistent with the changes in antigen presentation etc. on the tumor side and show that the local immune environment is altered, following CHA1 treatment.

We further defined how CHA1 might alter the tumor T-cell population by examining its effect on known immune checkpoints -PD-L1, PD-1 and CTLA-4. CHA1 treatment results in increased PD-L1 but decreased PD-1 and CTLA-4 in the syngeneic model, but only changes in PD-L1 in the human xenograft, immune compromised model. The significance of this pattern requires a review of pertinent findings. A central feature is the PD-L1-PD-1 interactions between tumor and T-cells (112), respectively, which inhibit proinflammatory CD8^+^ T-cells to promote an immunosuppressive environment in which the tumor cells escape destruction. The expression of PD-1 defines an exhausted T-cell and promotes the known immunosuppressive environment (117). Finally, high CTLA-4 prevents T-cell activation (118). The ICI therapeutics (anti-PD-1, anti-PD-L1, and anti-CTLA-4) were developed based on the many cellular and molecular studies defining immune checkpoints. The use of the ICIs to disrupt the cell-cell interactions then promote T-cell activation and immune destruction of tumor cells (112). Furthermore, elevation of PD-L1 is one characteristic of hot tumors and is often used as a clinical biomarker in decisions to determine which patients may benefit from ICI treatment. We also reasoned that PD-L1 status may provide insights into how CHA1 may influence the tumor-immune cell infiltration and action.

### Immune Checkpoint Patterns: PD-L1 is elevated, but PD-1 and CTLA-4 are reduced

We found that PD-L1 mRNA and protein expression are elevated in tumors upon CHA1 treatment in the human xenograft model (tumor-intrinsic) and in the mouse syngeneic models (tumor+immune). PD-L1 is expressed on tumor cells, antigen-presenting cells, and non-lymphoid tissues (113). Here, we focus our analysis on the tumor cells and immediate local environment, The gene expression of PD-L1, but not the related PD-L2 mRNA is increased in CHA1-treated tumors from both the human and syngeneic models (Fig 6A). Using anti-PD-L1 IHC to localize the expression to the tumors or immune microenvironment (Fig 6B), we found a robust induction of PD-L1 expression after CHA1 treatment in either model at levels greater than 50% of the tumor cells (the data from human xenograft tumor). This high level of PD-L1 expression is used in lung and other cancers to apply ICIs treatment in patients and with good outcomes ((8–13) reviewed in (14)).

**Figure 6:**
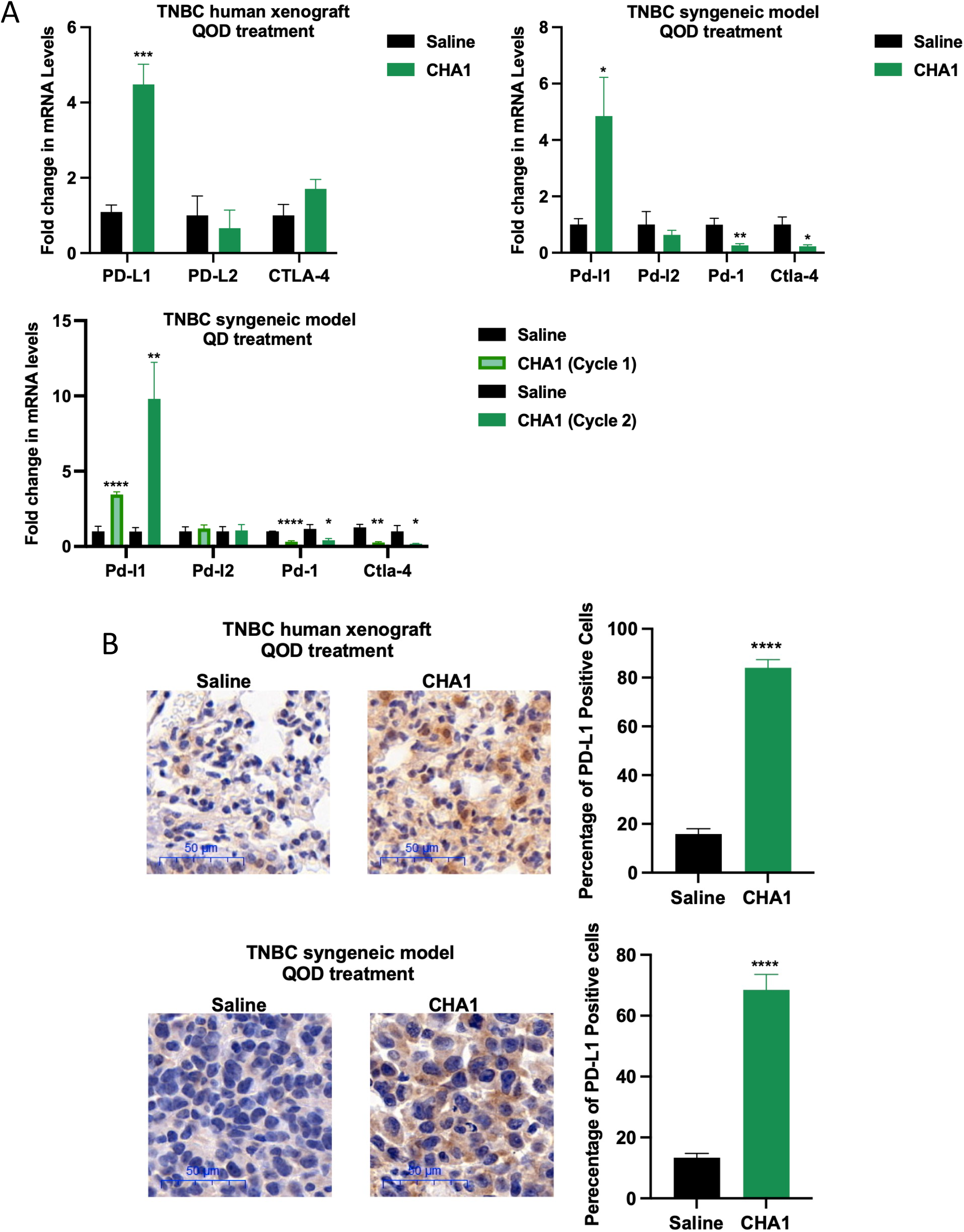
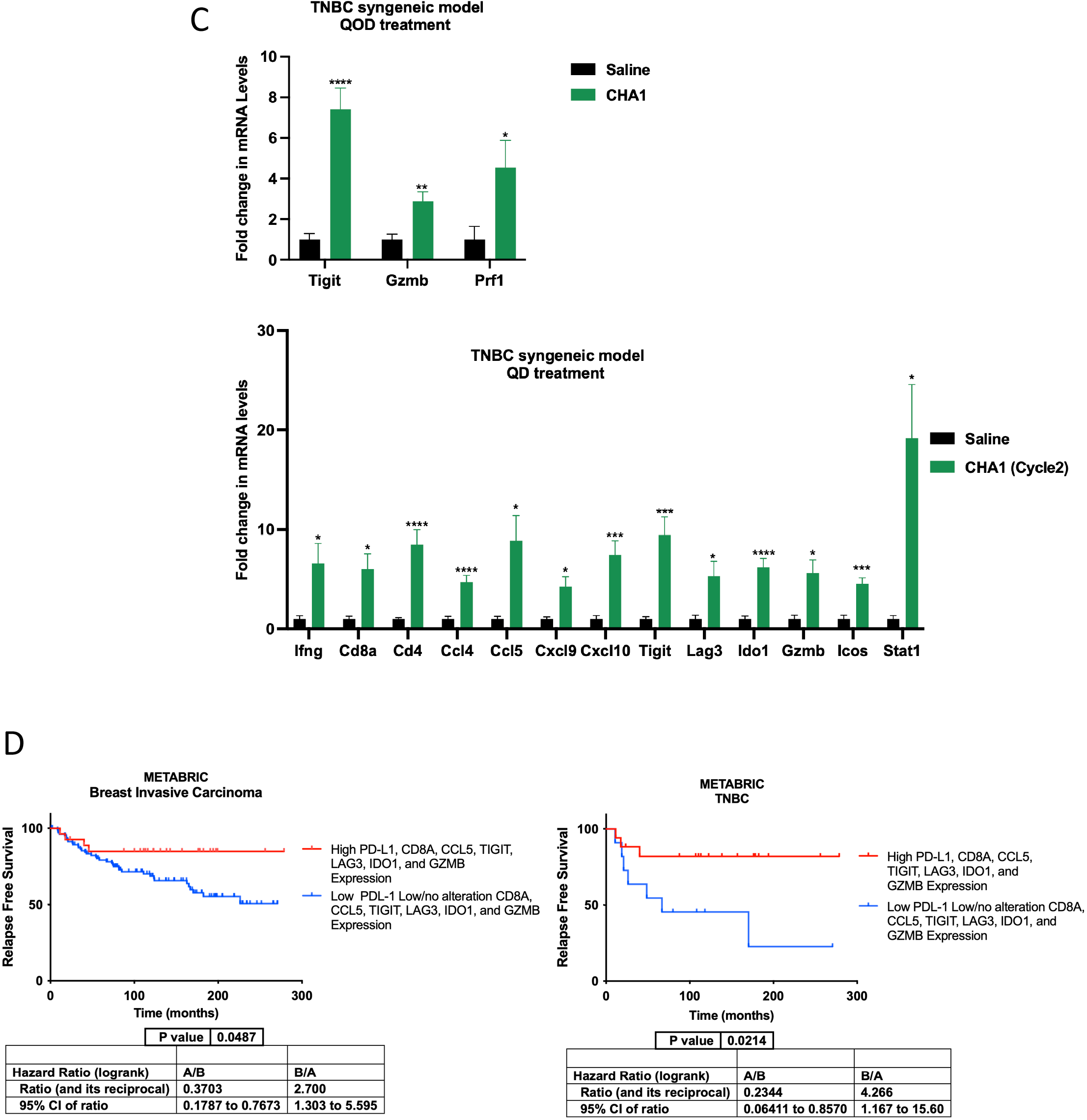
CHA1 upregulated PD-L1 expression and induced T-cell inflamed signature that associated with favorable patient prognosis. **A)** CHA1 altered the expression profile of immune checkpoints. CHA1 QOD treatment significantly upregulated the gene expression of PD-L1 and did not change the gene expression of PD-L2 and CTLA-4 in TNBC human xenograft model. CHA1 QOD and CHA1 QD treatment significantly upregulated the gene expression of Pd-l1, did not change the gene expression of pd-l2 and significantly downregulated the gene expression of Pd-1 and Ctla-4 in TNBC syngeneic mouse model. The gene expression was measured by qRT-PCR in saline and CHA1 treated tumors (human xenograft QOD, n=4-6 mice, syngeneic QOD, n=5-6, syngeneic QD cycle 1, n=3-4, syngeneic QD cycle 2, n=5-7). **B)** Immunohistochemistry of PD-L1^+^ cells indicated increased the protein expression of PD-L1 after CHA1 QOD treatment in TNBC human xenograft and TNBC syngeneic mouse models. Different sections of different untreated and treated tumors were stained with anti-PD-L1 antibody. Representative immunostaining section with quantification of the percentage of PD-L1^+^ cells in control tumors and treated tumors (n= 4 mice/group). **C)** CHA1 QOD and CHA1 QD treatment significantly induced T-cell inflamed signature by upregulation of the dendritic and CD8^+^ T-cells-associated genes in TNBC syngeneic mouse model. The gene expression was tested after CHA1 QOD (n=5) and CHA1 QD (n=5-7) treatment in TNBC syngeneic model. **D)** CHA1 T-cell inflamed gene signature was associated with improved prognosis in patients with invasive breast carcinoma and TNBC patients. Patients with high expression of PD-L1, CD81, CCL5, TIGIT, LAG3, IDO1 and GZMB had a prolonged relapse free survival. cBioPortal for Cancer Genomics was used to extract the data. “METABRIC” (83–85) was used to obtain the survival data. mRNA expression relative to all samples with z-score threshold of 1.3 was used to extract the data (refer to materials and methods). Kaplan-Meier survival curve with the hazard ratio (HR), 95% confidence intervals (CIs) and log-rank *P*-value was used for survival analysis (refer to materials and methods). **A-C, D)** Unpaired two-tailed t-test was used for the comparison between two groups. qRT-PCR experiments were done in triplicates in three independent experiments. Data were represented as mean ± SEM. *\* P < 0.05, ** P < 0.01, *** P < 0.005, **** P < 0.001*.

In the human xenograft, the increased PD-L1 mRNA and protein expression with CHA1 treatment reflected tumor expression and highlighted that induction of PD-L1 is intrinsic to the tumor. In addition, the induction effect of CHA1 is also more significant than upon use of either individual component (EGCG or DAC) (S7A Fig and S7B Fig). Lastly, PD-L1 is a known ISG as previously reported (6, 114). Accordingly, the level of STAT3 phosphorylation is positively correlated with PD-L1 mRNA levels in the human xenograft (S7C Fig). Together, CHA1 robustly induces PD-L1 in the tumor cells grown in syngeneic or immune compromised models and consistent with interferon signaling.

Another consequence of CHA1 treatment is a reduction of both PD-1 and CTLA-4 mRNA in the local T-cells in the syngeneic models (Fig 6A). Elevated PD-1 expression in T-cells is a marker of exhaustion or of a quiescent T-cell population (117). Activation of T-cells requires proliferation, often measured by Ki67 staining. One function of the ICI anti-PD-1 is to disrupt the PD-1-PD-L1 binding and promote T-cell activation with ensuing tumor destruction (112). Exhausted T-cells become hyporesponsive and they lack their effector functions (loss of IFNγ production) (118). Thus, the reduction of PD-1 expression upon CHA1 treatment (Fig 6A) is consistent with a relief of T-cell exhaustion. Additionally, while CHA1 treatment decreased proliferation and Ki67 staining in the tumors, CHA1 treatment had the opposite effect on CD8^+^ T-cells in the tumor, increasing in KI67^+^CD8^+^ T-cells. The PD-1 decrease and KI67^+^CD8^+^ increase in the local CD8^+^ T-cells are consistent with CHA1 treatment promoting activation of the recruited and local T-cells.

The final immune checkpoint that is targeted to enhance antitumor immunity is cytotoxic T-lymphocyte-associated antigen 4 (CTLA-4), which can be expressed by CD8^+^ T-cells, CD4^+^ T-cells, and T_reg_ (113). The ICI anti-CTLA-4 promotes tumor destruction by inactivating CTLA-4 and promoting T-cell activation. CTLA-4 expression is measured in saline and in CHA1 treated tumors of the syngeneic models. The expression profile is not altered after treatment in human xenograft model (Fig 6A) because T-cells are not present (100). However, the syngeneic mouse model shows that CHA1 treatment decreases the CTLA-4 mRNA, which is another marker of T-cell exhaustion (Fig 6A) (118). Together, the data from Fig 5 and Fig 6 suggest that CHA1 promotes cytotoxic T-cells activation and enhances their function. We cannot yet distinguish whether CHA1 acts directly on the local immune environment, or the activation of T-cells is an indirect consequence of tumor-intrinsic rearrangements. But CHA1 treatment appears to provide evidence of T-cell activation by decreases in immune checkpoints PD-1 and CTLA-4 and increases in proliferation (Ki67).

Finally, Fig 6 highlights the induction of PD-L1, but the reduction of PD-1 and CTLA-4 upon CHA1 treatment. First, PD-L1 is a critical clinical decision marker for determining whether patients receive immune checkpoint inhibitors, first in lung cancer, but also in other cancers such as TNBC. The fact that the percentage of cells expressing PD-L1 is greater than 50% is a critical factor in dictating ICIs treatment and is a known marker of the cold-to-hot transition. The decreases in PD-1 and in CTLA-4 suggest a relief of T-cell exhaustion and/or T-cell activation—as evidenced by an increase in Ki67^+^CD8^+^ T-cells. Together, these imply that CHA1 treatment promotes reduced tumor proliferation while reprogramming the tumor into a “hot” tumor with APC-like, epithelial-like properties and accompanied by activation of CD8^+^ T-cells 109.

### CHA1 promotes a T-cell inflamed signature and correlation with an improved patient prognosis

Recent evidence suggests that elevated PD-L1 expression is not the only biomarker dictating the response into immune checkpoint inhibitors. Several reports indicate that the response to immune checkpoint inhibitor arises in tumors with high expression of dendritic cells and CD8^+^ T cell–associated genes, which is called T cell” inflamed” phenotype (S7D Fig) (115, 116). In addition to PD-L1 induction, CHA1 treatment of the syngeneic model also induces many genes from the published T-cell inflamed gene signature from human tumors (Fig 6C and S7D Fig). This signature is a composite of the tumor and the local environment. By contrast, our RNA-seq data is derived from the human xenograft and reflected tumor-intrinsic changes. Thus, to better focus on the tumor changes, we used a Venn diagram with hypergeometric analysis to derive an overlap signature of T-cell inflamed genes and of CHA1 treatment. In other words, we sought to identify gene expression changes in the tumor that might promote T-cell inflammation or infiltration. The overlap genes and their functions are shown in S7D Fig. All the genes described in the above sections are members of the published T-cell inflamed set and some of these genes additionally are identified as part of the reprogramming to antigen presentation. In addition, CHA1 upregulates Psmb10, which is a T-cell inflamed gene, in the syngeneic mouse model (Fig 4A). Furthermore, our data demonstrates that the CHA1 upregulates MHC-II gene HLA-DRB1 (Fig 4A and S4A Fig), which is linked to the T-cell inflamed signature in human xenograft, indicates CHA1 reprograms the tumor cells themselves to express genes involved in T-cell activation. Thus, CHA1 increases the chance of successful response of immune checkpoint inhibitors.

What might be the potential molecular or clinical relevance? First, there was a positive correlation between expression of PD-L1 and of members of the T-cell inflamed gene overlap set in our syngeneic and human xenograft model (S7E Fig). We also showed that PD-L1 expression is positively correlated with T-cell inflamed genes in invasive breast cancer patient dataset retrieved from TCGA (S7F Fig). Thus, CHA1 is effective in induction of gene signature with PD-L1, and other T-cell inflamed gene signatures, thus increasing the likelihood of a favorable T-cell environment for maximizing success to ICIs.

Might there be prognostic relevance to the overall signature? Might CHA1 treatment induce a favorable prognostic signature? We use the literature and an in-silico test of the TCGA and METABRIC databases (82–85). First, the reprogramming to express MHCs in the tumor cells is a recent and known favorable marker of a better prognosis (91). We further investigate CHA1 overlap gene signature in METABRIC invasive breast cancer patient’s dataset. To achieve this, the patient dataset is classified into two groups based on mRNA expression of CHA1 overlap T-cell inflamed gene signature: patients with high expression of PD-L1, TIGIT, LAG3, IDO1, CD8A, GZMB, and CCL5 profile and patients with low/no alteration of PD-L1, TIGIT, LAG3, IDO1, CD8A, GZMB, and CCL5 expression profile. We found that invasive breast cancer and TNBC patients with CHA1/T-cell inflamed overlap gene signature-enriched had improved prognosis and prolonged survival (Fig 6D). Thus, the overlap gene set derived from our homegrown CHA1 signature, and the published T-cell inflamed signature appears to have some prognostic value and may predict a favorable outcome in invasive breast cancer patients in silico, likely through tumor changes that promote T-cell inflammation/infiltration. Furthermore, analysis of TCGA and METBRIC dataset demonstrates that CHA1 gene signature including PD-L1, TIGIT, HLA-B (MHC-I), HLA-DRB1 (MHC-II), OAS2 (ISG), IFI27 (ISG) and HBP1 (Wnt inhibitor) could predict favorable outcome in invasive breast cancer patients (S8 Fig).

### CHA1 Collaborates with Immune Checkpoint Inhibitor Anti-PD-L1

The above data predicts a model by which CHA1 uniquely enacts a cold-to hot transition and reprograms the tumor towards enhances antigen presentation and promotes T-cell inflammation/infiltration. There may be additional effects to promote activation of the recruit T-cells. The signature induction is the important protein and clinical biomarker PD-L1, with concomitant reductions in the inhibitory immune checkpoints PD-1 and CTLA-4. The data demonstrating CHA1 reprograms both the tumor and the tumor microenvironment further suggests that may endow susceptibility to immune checkpoint inhibitors. Thus, we deliver CHA1 on a 5-day cycle, rested, and then treat with either saline or ICI (Fig 7A). Only anti-PD-L1 gave an observable result. As shown in Fig 7B, pre-treatment with CHA1 resulted in reducing tumor size and weight, but the dose of anti-PD-L1 alone had no effect. The combination of CHA1 pre-treatment together with anti-PD-L1 gave an additional and significant diminution in tumor size and weight. As shown in S9B Fig, CHA1 plus anti-PD-L1 treated tumors have a more necrotic morphology. And anti-CTLA-4 or anti-PD-1 have not yet shown any single or collaborative effect at the doses used (data not shown).

**Figure 7:**
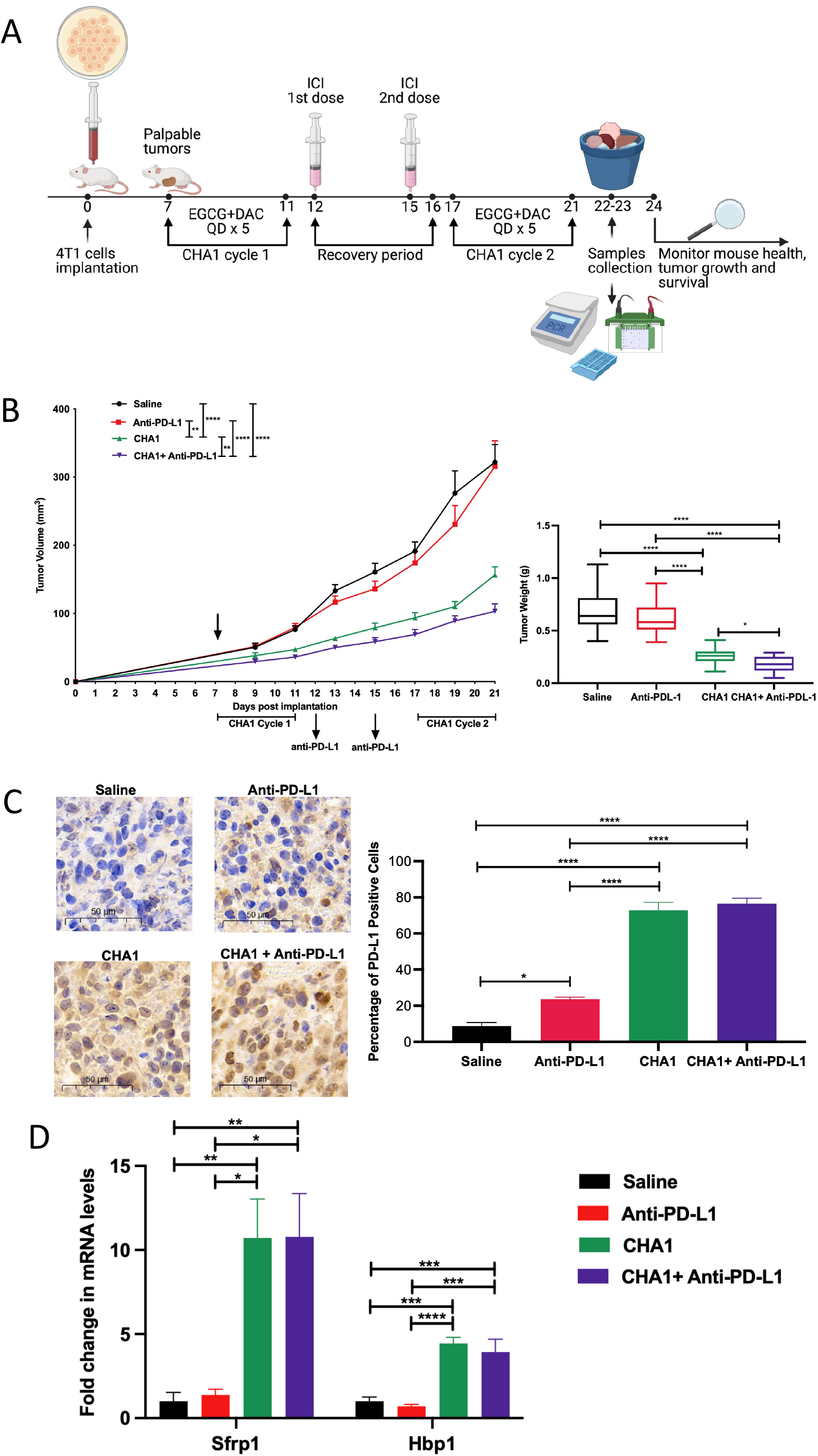
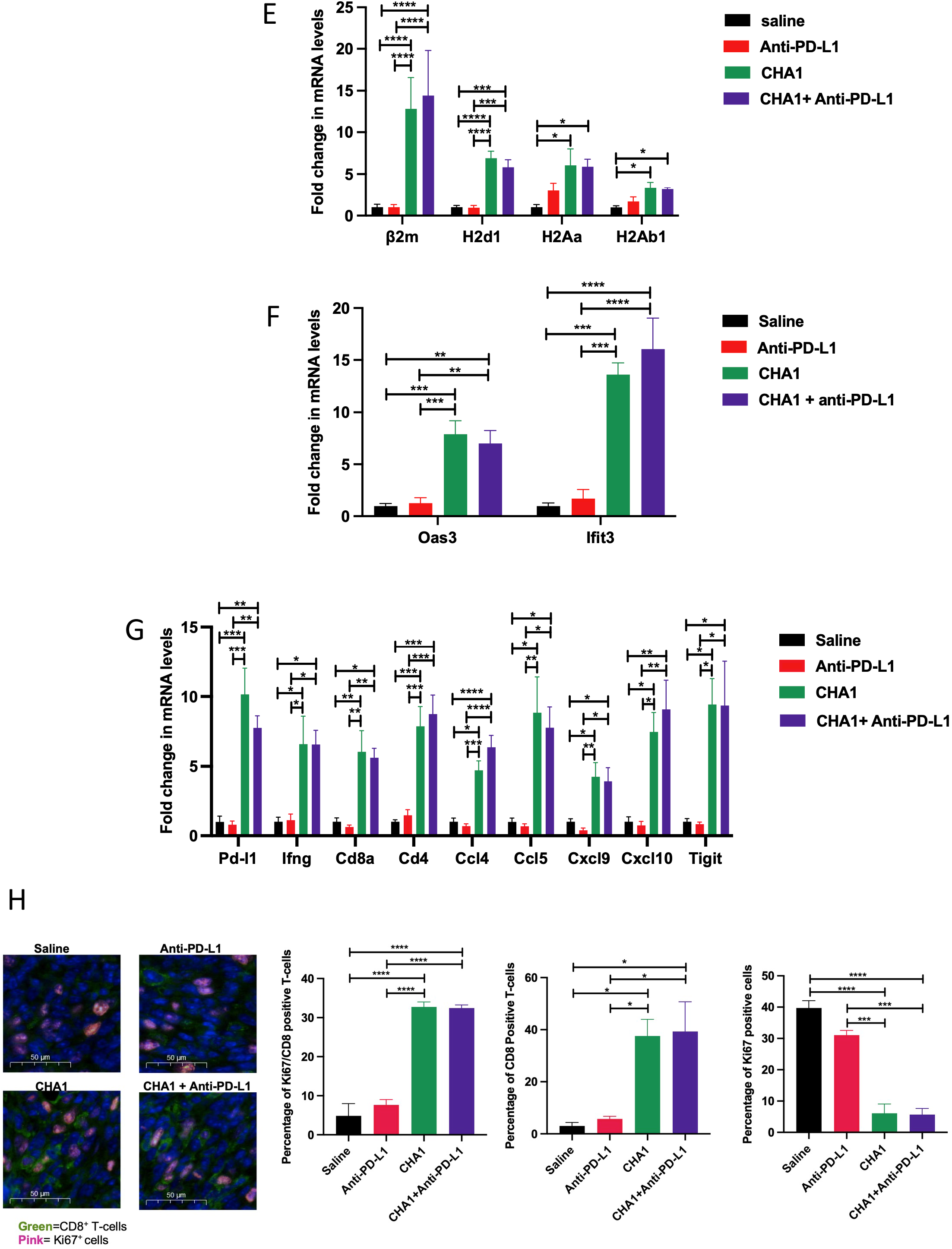
CHA1 enhanced anti-tumor effect of anti-PD-L1 in TNBC syngeneic mouse model. **A)** Study design of CHA1 treatment in combination with anti-PD-L1. The CHA QD treatment timeline was followed as in Fig 1E. Anti-PD-L1 (100 µg/mouse) was administrated on day 12 and day 15 during recovery period. Fig 7A was created with BioRender.com. **B)** CHA1 as collaborative agent with anti-PD-L1. The combination treatment of CHA1 with anti-PD-L1 significantly slowed the tumor growth and reduced the tumor weight (n=15-23). The results were combination of three different studies The 4T1 cells were implanted orthotopically into Balb/c mice. Mice were treated with saline, anti-PD-L1, CHA1, CHA1+anti-PD-L1 for the indicated time. Tumor volume was measured with calipers. Tumors were weighted at the end of the experiment. **C)** Immunohistochemistry of PD-L1^+^ cells indicated increased the protein expression of PD-L1 after anti-PD-L1, CHA1 and CHA1+anti-PD-L1 treatment. Different sections of different untreated and treated tumors were stained with anti-PD-L1 antibody. Representative immunostaining section with quantification of the percentage of PD-L1^+^ cells in control tumors and treated tumors (n= 3 mice/group). **D-G)** The addition of CHA1 into anti-PD-L1 improved the response to anti-PD-L1 through **D)** suppression of Wnt signaling by upregulation of the gene expression of Wnt inhibitors Sfrp1 and HBP1, **E)** activation of antigen presentation by upregulation of the gene expression of MHC-I (β2m and H2d1) and MHC-II (H2Aa and H2Ab), **F)** stimulation of IFN response via upregulation of ISGs (Oas3 and Ifit3), and **G)** activation of CHA1 gene signature associated with T-cell inflamed tumor microenvironment. **D-G)** The gene expression was tested by qRT-PCR after saline, anti-PD-L1, CHA1, CHA1+ anti-PD-L1 treatment (n= 5-8 mice/group). **H)** Immunofluorescent co-staining of Ki67^+^CD8^+^T-cells revealed that the addition of CHA1 treatment into anti-PD-L1 increased the infiltration of actively proliferating IFNγ-secreting cytotoxic Ki67^+^CD8^+^ T-cells. Different sections of different untreated and treated tumors were stained with anti-Ki67 and anti-CD8 antibody. Representative Immunofluorescence staining section of Ki67^+^CD8^+^ T-cells with quantification of the percentage of Ki67^+^CD8^+^ T-cells, CD8^+^ T-cells and Ki67^+^ tumor cells in control tumors and treated tumors (n= 3 mice/group). **B)** Mann-Whitney U-test was used for tumor volume and tumor weight comparisons. **C-H)** Adjusted one-way ANOVA was used for multiple comparison. qRT-PCR experiments were done in triplicates in three independent experiments. Data were represented as mean ± SEM. * *P* < 0.05, ** *P* < 0.01, *** *P* < 0.005, **** *P* < 0.001.

Next, we assessed PD-L1 status, Wnt signaling, antigen presentation, IFN signaling, PD-L1 and T-cell inflamed signature and tumor CD8^+^T-cell activation status in each of the treatment groups. In Fig 7C, IHC of PD-L1 shows a robust induction in the CHA1 and CHA1+anti-PD-L1 groups. Fig 7D shows a likely inhibition of Wnt signaling, as evidence by robust induction of the Sfrp1 and Hbp1 Wnt inhibitors. Fig 7E shows that several MHCs are elevated with CHA1. Lastly, two representative IFN genes (Oas3 and Ifit3) are induced (Fig 7F). Several of a T-cell inflamed signature (above) are induced (see Fig 7G). Lastly, the Ki67 profiles show that tumor cells are diminished in the CHA1 and CHA1+anti-PD-L1 group, but there are increased in active proliferating IFNγ-secreting cytotoxic Ki67^+^CD8^+^ T-cells group (Fig 7H), again suggesting reduced tumor proliferation but heightened T-cell proliferation/activation with treatment. The gene expression effects are dominated by CHA1, when present in the treatment group and show a sustained response with no additional effect with anti-PD-L1. This suggests that CHA1 actions are independent of anti-PD-L1 effect since they act upstream of PD-L1 and may have similar effects. Alternatively, CHA1 pre-treatment may endow a susceptibility to anti-PD-L1 action. A limitation of the 4T1 immune competent TNBC model is the amount of ICI drug that can be used. Anti-PD-L1 has known effects in the bone marrow (119). In addition, the 4T1 tumor itself has a deleterious effect on the hematopoietic system resulting in splenomegaly and granulocytosis (120). In addition, it has been reported that more than two doses of anti-PD-L1/anti-PD-1 in 4T1 tumor-bearing-Balb/c mice caused a hypersensitivity reaction resulting in death (120, 121). These mitigating factors all limit an expanded dosing of anti-PD-L1 in our models. However, the data all are consistent with a mode of action that addition of CHA1 enhanced the immune response to anti-PD-L1. These require a dual signaling action in the inhibition of Wnt signaling with concomitant activation of IFN signaling. The dual pathway action reprograms the tumor cells to endow APC and facilitate the recruitment and activation of the resident CD8^+^ T-cells. A critical aspect is the robust induction of the PD-L1 and of an inflamed T-cell signature, both of which represent an established and emerging clinical criterion, respectively, for determining efficacy of ICIs treatment (115, 116).

## Discussion

### Summary

In this study, we elaborate a new therapeutic strategy CHA1 for inhibiting Wnt signaling but discovered that CHA1 did far more in altering the tumor microenvironment and improving immunotherapy for TNBC, the initial design of CHA1 treatment (Fig 1A) was to inhibit Wnt signaling as a means of inhibiting tumor growth, which was successful. Intriguingly, the unbiased analysis of our RNA-seq from the human xenograft model, while further confirming the inhibition of Wnt signaling, also revealed greater functionality for CHA1 action. Our goal was to thoughtfully consider strategies and minimize the known side-effects for the patient. The compound combination elaborated here (EGCG and DAC) have well known safety profiles and modest effects on tumors when used independently, but when used together (CHA1) have far greater effects. Together, the summary depicted in Fig 8 point to widespread consequences of CHA1 treatment affecting the basic biology of the TNBC tumor cells and with similarly widespread biological and clinical implications, which will be discussed below.

**Figure 8:**
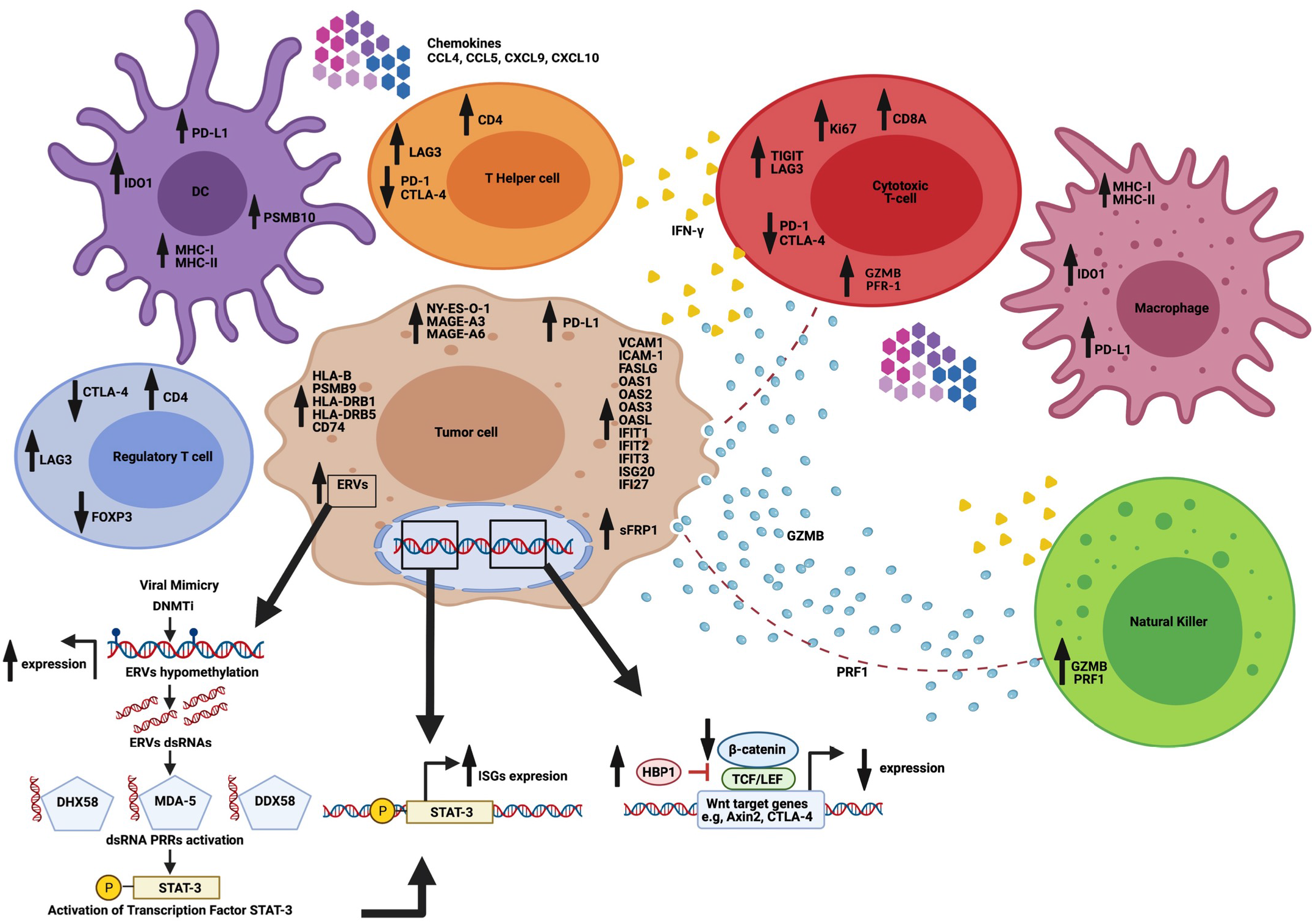
CHA1 reprograms tumors and alters tumor-immune cell interaction. CHA1 can convert cold tumors into hot tumors by altering key signaling pathways and cellular processes and reprogramed a tumor cell to have antigen presentation properties, upregulate PD-L1, increase effector immune cells infiltration, decrease immune suppressor cells infiltration, and induce T-cell inflamed signature. Thus, CHA1 could be an ideal collaborative agent to enhance immune checkpoint inhibitors (ICIs) response. CHA1 suppressed Wnt/β-catenin pathway, which is a tumor intrinsic factor that play important role to determine the response into ICIs, by induction of Wnt inhibitor including sFRP1 and HBP1. CHA1 also activated IFNα/β and IFNγ pathways, that correlated with successful ICIs treatment, and upregulated interferon stimulated genes (ISGs). One possible mechanism was through activation of viral mimicry. Also, CHA1 induced CTAs on tumor cells that can result in increased immune recognition of tumor cells and therefore, activation of T-cells to secrete IFNγ. In addition, CHA1 increased the infiltration of activate cytotoxic IFNγ-secreting Ki67^+^ CD8^+^ T-cells. The production of IFNγ by activated cytotoxic T-cells, and other cells such as T-helper cells and natural killer (NK) cells resulted in stimulation of key downstream signaling molecules. IFNγ can upregulate the expression of MHC-I, MHC-II and other components of the immunoproteasome and antigen-presenting machinery. Activation of antigen presentation is important for T-cells activation, resulting in tumor cells elimination. Thus, antigen presenting cells (APCs) such as dendritic cells (DC), macrophages and tumor cells can present the antigen to cytotoxic CD8^+^ T-cells via MHC-I or to T-helper cells via MHC-II. One important effect of CHA1 was induction of antigen machinery in APCs including tumor cells. Furthermore, CHA1 induced the expression of chemokines such as CCL4, CCL4, CXCL9, and CXCL10 that function to recruit additional cytotoxic CD8^+^ T cells. Moreover, CHA1 upregulated perforin (PRF1), and granzymes (GZMB). Active CD8^+^ T cells and NK cells can secrete PRF1 and GZMB which play important role in apoptosis. Furthermore, CHA1 upregulated the expression of T-cell inflamed signature. One of the functions of IFNγ is induction of a host of checkpoints, such as PD-L1, on the surface of macrophages, DCs and tumor cells. In addition to upregulation of PD-L1, other checkpoint molecules including TIGIT, LAG3, and IDO1 can also be homeostatically upregulated by T cell activation and IFNγ signaling in order to restrain the antitumor immune response. In addition, CHA1 downregulated T-cell exhaustion markers including PD-1 and CTLA-4 which is Wnt target gene and decreased the infiltration of immune suppressor FOXP3^+^ Treg. Overall, CHA1 function through an array of effector molecules to induce a T-cell inflamed tumor microenvironment (TME) that crucial for ICIs effectiveness. Fig 8 was created with BioRender.com

While CHA1 is not a single target therapeutic, the compilation of its diverse mechanisms (Fig. 8) reveals unexpected clinical and biological implications for tapping an unappreciated biological network, triggering re-programming of tumor cells and revealing applications as a collaborative agent for ICI compounds. Under the umbrella of the biologically diverse cold-to-hot transition, the CHA1 combination inhibits both Wnt signaling and activates interferon signaling, which are the two major signaling pathways that dictate the change from an immune suppressive (cold) tumor to an immune-active (hot) tumor. Neither component of CHA1 was effective and it is the combined actions that have maximal efficacy. The full CHA1 acts to alter both Wnt signaling and IFN, which appears necessary together (S1 Fig, S2 Fig, Fig 2, Fig 4D, 4E, and S4 Fig). The combined and reciprocal Wnt and interferon signaling enacts a fundamental biological reprogramming of the otherwise epithelial tumor into a tumor with antigen presentation functions, but also with cancer testes antigen, and mesenchymal-epithelial properties initiated through an epigenetic deregulation, at least in part through a viral mimicry mechanism—which in turn activates IFN signaling, which is the key to APC expression and important immune checkpoint proteins such as PD-L1. The detailed analyses to characterize tumor immune changes and depicts the criteria that are emblematic of involved cell types. Careful analyses of CHA1 for tumor-intrinsic and overall function demonstrated a far broader alteration of the tumor and its microenvironment. CHA1 treatment also resulted in activation of immune-related pathways, including the IFN pathway, induction of antigen presentation and upregulation of a T-cell inflamed gene signature, including increased PD-L1, infiltration of active proliferating IFNγ-secreting cytotoxic Ki67^+^CD8^+^ T-cells, F480^+^ macrophages and decreased infiltration of FOXP3^+^ T_reg_. These are all consistent with a shift to an immune active tumor resident environment, and away from an immune-suppressive phenotype. Finally, the combination of CHA1 pre-treatment and anti-PD-L1 was significantly more effective than either compound alone, although CHA1 had a larger effect (Fig 7B).

Many TNBC tumors have a common phenotype of hyperactive Wnt signaling (23). Activation of Wnt signaling has been linked to breast cancer cell growth and to recurrence of the disease (25, 122). β-catenin levels are elevated by Wnt signaling, which is thought to contribute to the poor prognosis of patients with TNBC (28). The data from both human xenograft and syngeneic model revealed that CHA1 induces Wnt inhibitor sFRP1 and HBP1 as well as reduces β-catenin protein levels and AXIN2, a Wnt target gene. The data from human xenograft suggested that both drugs were required to inhibit Wnt target gene AXIN2. Several studies reported EGCG suppressed Wnt signaling in colon cancer cells, lung cancer stem cells and breast cancer cells (59, 123, 124). However, previous work showed that DAC monotherapy was not able to downregulate Wnt target genes in AML cell lines and mouse model (125).

The second reason for failure of ICI therapy (in melanoma) was dysfunction of JAK/STAT/IFN pathway (6). Further, high Wnt signaling results in decreased T-cell infiltration into tumor sites which is critical for ICI efficacy (6, 32, 126–128). Activation of IFN signaling promotes innate and adaptive immune response (107). In addition, active IFN signaling is required for proper anti-tumor immune response, ICI efficacy and for activation of tumor antigen presentation (6). In addition, IFN activation induced PDL-1 expression (6). It has been reported that PD-L1 is regulated by JAK/STAT/IFN pathway (114) and that mutations in the interferon receptor pathway were linked to resistance to anti-CTLA-4 or anti-PD-1 in metastatic melanoma patients (129). GSEA analysis of the RNA-seq indicates activation of IFNα/β and IFNγ pathways after CHA1 treatment. Our data shows that CHA1 increases phosphorylation status of STAT3 in both human and syngeneic models, indicating involvement of IFNα/β and IFNγ pathways (107). Looking at downstream targets, CHA1 treatment markedly increased the expression of IFN stimulated genes (ISGs). In addition, our data from TCGA indicated CHA1 ISGs such as IFIT2 and OAS2 are associated with increased survival. We investigated the possible source of IFN stimulation, and a major possibility is induction of viral mimicry via alteration of the epigenome. Several reports from studies of decitabine suggested that part of its efficacy lies in the hypomethylation of ERVs (39). Previous work reported that DNMT1 inhibitors enhanced the efficacy of immune checkpoint inhibitor anti-CTLA-4 in melanoma by trigger IFN response via viral mimicry (39). Therefore, we examine whether CHA1 induces human endogenous retroviruses (ERVs) and muERVL-1. Indeed, human ERVs and muERVL-1 are induced by CHA1 (Fig 4F).

A key aspect to our investigations was the use of both the human xenograft (immune compromised) and syngeneic (immune-competent) models. The human xenograft combined with the RNA-seq analysis focused on human gene allowed a unique snapshot of the gene expression changes intrinsic to the tumor and without a complicating full mouse immune system. Some analyses of mouse genes in this model provided limited insights into the defective immune system, as we saw induction of mouse perforin, consistent with the remnant NK cell function. Thus, our studies will emphasize the tumor changes over the immune environment. The syngeneic model combined both tumor and tumor-resident immune environment, although either can be discerned by single cell imaging or methods. It is fortuitous that the human xenograft was used first, so that we could craft hypotheses for a sensible investigation of the environment in the syngeneic specimens and confirm the tumor intrinsic changes. In this way, we had the opportunity to discover new insights from the tumor changes and which would have been missed, had we used only the syngeneic models in unbiased analyses.

### Biological Implications

An exceptionally striking observation upon CHA1 treatment is the re-programming and transformation of a TNBC breast tumor cell into one that now robustly expresses antigen presentation proteins and processing machinery and then further engages a vastly altered tumor-resident immune environment. These observations highlight that CHA1 treatment triggers a certain plasticity and perhaps a universal path in an otherwise epithelial tumor to express genes attributed to professional T-cells and APC cells This unusual property additionally promotes a functional tumor-resident immune environment with CD8^+^ T-cells that collectively overcomes the usual immunosuppressive environment. In recent years, the expression of a tumor-derived MHC has correlated with a more favorable prognosis, so understanding this expression pattern and developing therapeutics like CHA1 has biological and clinical importance. Thus, an epithelial re-programming and a functional switch from immunosuppressive to immune-active in the tumor-resident environment. We believe CHA1 represents a potential therapeutic that fortuitously engages interferon and Wnt signaling to create a unique reprogramming. Notably, the results section and above summary figures highlight that the CHA1 has dual and opposite action of Interferon and Wnt signaling to effectuate an important reprogramming in the tumor and tumor-resident immune environment.

We believe that the biological re-programming tapped by CHA1 treatment provides further insight into an unappreciated alternative cellular phenotype when the cellular epigenetics balance is disrupted by chemical or genetic means. Our work here and several other published studies are consistent with the identification of an unappreciated epithelial plasticity that may also alter the tumor-resident immune environment away from immunosuppression. Collectively, our work in the context of the field may provide an important molecular insight into what constitutes a “hot” tumor and the importance of disrupting or of regulating the cancer epigenome. In putting a context to the completed CHA1 studies, we noticed several diverse published studies that nonetheless reported as (unexpected phenotype) and unbiased RNA-seq analysis that shows antigen presentation or other immune processes as major with hundreds of genes, many of which are prevalent in T-cells or interferon signaling, but certainly not to an otherwise epithelial tumor. Similar phenotypes to our CHA1 studies here were triggered by very diverse signals-including DNMT1 inhibitors, CDK4/6 inhibitors (42, 43, 130), CDK9 inhibitors, LSD1 inhibitors or LINE regulation. The initiating signals represent the diversity of cellular function with no obvious biological pattern. Yet, each report highlighted that the immune phenotypes in the epithelial tumor cell were initially surprising, but each also landed upon a similar mechanism, as we have elaborated for CHA1. Collectively, we argue for vastly different agents, including CHA1 that engages an intricate and fundamental epigenetics-based network that may determine an alternative cell fate as a tumor-derived antigen presenting cells, highlighting a usual epithelial plasticity and implications for tumor-resident immune interactions and function.

In the present paper, the CHA1 combination gives the maximal expression of MHC and other APC genes, while neither component at the concentration used has a demonstrable effect on tumor phenotype or gene expression. Part of the CHA1 mechanism point to an IFN induction and STAT3 activation for the expression of ISGs, many of which are part of MHC and APC gene regulation and have important roles in other immune functions. The likely source of IFN signaling results from a viral mimicry response that triggers IFN production and the above signaling to express genes for MHC and its machinery (Fig 4F). Our studies confirmed the bona fide expression of MHC-I and MHC-II proteins in the TNBC tumor sections. We also noticed that a consequence of disrupting the cancer epigenome is an acquisition of unusual and unexpected antigen presentation properties in the tumor cell, which we will denote as an alternative epithelial fate. The cancer epigenome is a catch-all term for epigenetic modification in the cancer genome: DNA methylation, chromatin remodeling, histone methylations, histone acetylation. Furthermore, the common theme to achieve this alternative epithelial fate with tumor-localized antigen presentation and processing is an epigenetic disruption that was nonetheless achieved by a plethora of unrelated triggers. Our work and each of the examples highlighted below shows that different epigenetic disruptions all lead to the re-expression silenced heterochromatic regions such as endogenous retrovirus (ERVs) or repeat elements (e.g., LINES) and a viral mimicry response to produce Interferons that then increase MHC genes and other antigen presentation and processing genes. A remarkable correlation is that each of the mechanisms outlined below have global epigenetic effects, yet the predominant functional manifestation is the de-regulation of heterochromatin and re-expression of previously-thought “junk” DNA. These published studies also attest to the unexpected importance of heterochromatin and in maintaining the boundaries with euchromatin, which is the site of more actively transcribed genes. For example, while ERVs are present in heterochromatin, epigenetically silenced genes such as sFRP1, would be likely found in euchromatic regions.

The pioneering work of Chiappinelli, Baylin and colleagues first addressed that inhibition of DNA methylation by the DNMT inhibitor DAC in cells triggered a viral mimicry response, interferon signaling, and induction of a genes associated with antigen presentation and processing (38, 131). These studies attribute global disruption of DNA methylation. Additionally, another study using a CDK4/6 inhibitor designed to work at G1 in the cell cycle nonetheless also resulted in APC re-programming in breast cells with a likely mechanism as a decrease in DNMT1 gene expression and triggering a global decrease in DNA methylation and re-expression of ERVs, ds RNAs, and IFN signaling activation (42). These studies also attribute global disruption of DNA methylation. However, other reports indicate that other epigenetic players can be engaged to achieve the same effect. One report highlighted that inhibitor of CDK9, a cell cycle kinase and transcription factor regulator, nonetheless triggered reprogramming to express APC genes and machinery to enhance infiltration of tumor resident T-cells. A viral mimicry mechanism was also reported, but the epigenetic disruption worked through a chromatin remodeling protein (41). Two reports featured an inhibitor of LSD1, which is an H3K4 demethylase (40, 132, 133). LSD1 (KDM1) was the first histone demethylase identified in landmark studies by Shi and colleagues. H3K4me3 is mark of active gene transcription, so LSD1 inhibition would likely promote H3K4 methylation (by preventing demethylation) or regions of active transcription to disrupt silenced and/or heterochromatic regions, where ERVs may reside. A nuance of these beautiful studies is that LSD1 did not bind to any promoters for IFN genes, but instead bound to ERV promoter, implying the H3K4 demethylation is part of the mechanism for transcriptional silencing of ERVs. LSD inhibitors would then disrupt this regulation and promoted H3K4 methylation and transcriptional activation. Finally, some reports have examined re-expression of LINE and other repeat elements. Classon and colleagues implicated a specific H3K9 methylase, known to be critical for heterochromatin formation and maintenance (134). Chiappinelli et al reported increase in repeat element expression, another component in heterochromatic silenced regions (135). Finally, other studies implicate PRC2 and EZH2 in a similar mechanism (48, 136). H3K27me3 is widely associated with silenced genes and with DNA methylation. Together, these studies and our own with CHA1 cover a panoply of epigenetic disruptions and widely different biological triggers. The take-home message is that regulation of the cancer epigenome can specify cellular fate and reprogramming with a phenotype for the acquisition of APC properties. The epithelial-like tumor cells have an unexpected plasticity and intrinsic ability to exhibit antigen-presentation and other properties, which then dictate interactions with the tumor-resident immune cells. The widespread engagement suggests an intriguing and fundamental epigenetically regulated network—to achieve some features of a” hot” tumor. Importantly, the re-expression of antigen presentation and processing properties may be a clinical and biochemical readout of this epithelial phenotype upon epigenetic disruption.

Our work showed that CHA1 induced the expression of CTAs, notably NYESO1 (Fig 4C). Previous studies have found that DAC treatment results in the upregulation of CTAs on tumors and in hematologic tumor cells (101, 137, 138). Moreover, it has been reported that there was complete remission in a patient with a relapsed solid tumor who received treatment with DAC and a CTA-based dendritic cell (DC) vaccine. After vaccination, the patient had an increased number of MAGE-A3-specific T cells, along with recognition of tumor cells by the immune system (139). Numerous types of auto-immunogenic tumor antigens have been identified including cancer testis antigens (CTAs) (140). The expression of CTAs (e.g., MAGE, SSX gene families, PRAME, NY-ESO-1, and SP17) are restricted to solid tumors and to immune-privileged sites such as the testis, the placenta, and during fetal development (140). Malignant cells evade immune recognition by epigenetic modification of some key immune stimulatory molecules. The expression of CTAs, major histocompatibility complex class-I (MHC-I) and major histocompatibility complex class-II (MHC-II) are regulated by DNA methylation (101, 141–143). In cancer, the transcription of these genes is often silenced by hypermethylation. Demethylating agent such as DAC can reactivate these methylation-silenced genes (101, 141–143). Our data show that CHA1 induces MHC-I and MHC-II in human xenograft and syngeneic models. The upregulation of MHC-I and MHC-II in human xenograft reveals that CHA1 manipulated the tumor cells to become APCs. Also, our data from TCGA showed that high expression of MHC-I HLA-B and MHC-II HLA-DRB1 is associated with increased survival.

### Functional Epithelial Plasticity and CD8^+^T-cell activation?

The CHA1 tumor re-programming phenotype (Fig 4, S4 Fig, S5 Fig) suggests that CHA1 and the above inhibitors/mechanisms work through epigenetic disruption, resulting in IFN-stimulated gene and protein expression of antigen processing and presentation machinery. An important question is whether expression of MHCs and CTAs are functional. Does the tumor re-programming promote immune cell infiltration to the tumor-resident environment? Are CD8^+^ T-cells increased in the local environment? Are the CD8^+^ T-cells active or exhausted?

Our data suggests that the Wnt signaling regulatory function of CHA1 is likely to be important for CD8^+^ T-cell recruitment and that it is both the actions on Wnt signaling and IFN signaling that provide the observed effects. For our work in this paper, the DAC or EGCG concentrations, when used singly, have little effect on tumor growth and at best modest effects on measurable parameters of signaling in the tumors. Our data suggest that the dual action of suppressing Wnt signaling and activating interferon signaling of CHA1 is necessary for inducing epithelial tumor changes with infiltration of active CD8^+^ T-cells. One of the biochemical mechanisms associated with cold tumors and resistance to ICIs is elevated Wnt signaling, a frequent manifestation of TNBC and cancer with a poor prognosis. Recent discoveries have underscored that Wnt signaling in numerous cancers encourages immune evasion and is counterproductive to the necessary immune cell infiltration that is necessary for ICIs efficacy and immune-mediated destruction (29). Wnt signaling has been linked to resistance to ICIs in melanoma (6). High Wnt signaling results in decreased T-cell infiltration in numerous tumors, a process that is critical for ICIs efficacy (6, 32, 126). it has been difficult to inhibit Wnt signaling therapeutically, with few effective in clinical trials with numerous adverse consequences, although numerous inhibitors exist for laboratory studies. A likely explanation is that some level of Wnt signaling is required for normal tissue maintenance and for stem cells, so a “just right” dosing is necessary. Thus far, the CHA1 components have not shown side effects that appear with targeted Wnt signaling. As noted, Porcupine (Wnt ligand maturation enzyme) inhibitor adverse effects include loss of taste and a prevalence of bone breaks (144). Nonetheless, the Wnt signaling inhibition of CHA1 is necessary to realize the functionality of the tumor cell reprogramming towards APC.

A common feature of CHA1 and of the LSD1 inhibitor (133) is that both lead to increased tumor infiltration of CD8^+^ T-cells. While CD8^+^ T-cells might be recruited to the tumor-resident environment, exhausted T-cells would still define an immunosuppressive environment. Of course, it is the ICIs that disrupt the interaction between PD-1, PD-L1 and/or CTLA-4 to promote activation, or cell cycle entry, but prior activation may improve efficacy (see below). For CHA1 treatment, two lines of evidence indicate that the tumor-resident CD8^+^ T-cells are active and not exhausted. First, the expression of PD-1 and CTLA-4, which are markers of T-cell exhaustion are reduced with Cha1 treatment, supporting an activation. Second, the number of CD8^+^ T-cells that are also Ki67 positive are increased. Ki67 is a marker of proliferation and of T-cell activation. Yet, the number of Ki67^+^ cells decreased in the tumor. Thus, CHA1 had an opposite proliferative effect on the (re-programmed) tumor and on the resident CD8^+^ T-cells, indicating opposite functions depending on cell type. The exact reciprocal scenario was reported by Shi and colleagues for LSD1 inhibitors (40, 133). The important, but differential and opposite effect of CHA1 and LSD1 inhibitors with respect to proliferation needs further investigation.

In considering our findings with CHA1 and the literature with fundamentally similar findings for other epigenetic disruptors, a plausible hypothesis is that the appearance of antigen presentation and processing genes and protein in an epithelial tumor is a practical and functional readout of an epigenetic re-arrangement engaged by numerous and diverse signals. CHA1 and the collective studies also re-enforce the parameters necessary to attain a “hot” tumor, or one with maximal susceptibility to immune destruction by engaging the recruitment and activation of CD8^+^ T-cells. This notion was applied in a recent paper that outlined a sophisticated chemical screen to identify compounds that induce APC properties and activated the local T-cell environment (37) and focused on breast cancers. The resulting screen identified LSD1 and HDAC inhibitors. Similarly, the results of our results of our systematic mechanistic dissection also create a platform for discovering therapeutics with the property of conferring susceptibility to ICIs and enacting the cold-to-hot tumor transition.

### Perspectives for CHA1 in improving ICI efficacy

The very intriguing biological and functional insights have important implications on how ICIs can be better applied for treating TNBC and possibly other cancer. While ICIs are game-changing treatments that activate the local immune environment to destroy tumors, they work on tumor types that express machinery to engage the resident immune system, termed “hot” tumors. While there are many successes, the majority (>75%) tumors respond poorly to ICIs and are termed “cold” for immune system engagement (8, 15). The important challenges are identifying responsive tumors and defining therapeutic strategies to convert tumors from “cold” to “hot”’ i.e., to engage the local immune environment, increasing tumor responsiveness to ICIs. There is noted tumor intrinsic resistance to ICIs (7).

Triple negative breast cancer (TNBC) is one of the cancers that has limited treatment options and uneven ICI efficacy (8, 15, 16, 145–147). Treatment of TNBC with immune checkpoint inhibitors has recently gained significant positive and negative attention and still represents a challenge with room for improvement. A recent study classified TNBC based on spatial immune cell contexts in relation to patient prognosis and response to anti-PD-1 as well as pathways of T-cell evasion into excluded, ignored and inflamed (148). The inflamed phenotype showed a favorable response to ICIs. The other two types (ignored and excluded) was resistance to ICIs (148). The ignored phenotype is characterized by low densities of CD8^+^ T-cells and activation of Wnt signaling with up regulation of Wnt target genes (148). Intrinsic Wnt/β-catenin pathway has also been linked to disfavor T-cells inflamed microenvironment and exclusion across cancers, including TNBC (29). High Wnt signaling is also correlated with brain metastases (149). ICIs are also under consideration for fatal brain metastases (26, 27).

Multiple factors play important roles in determine the responsiveness to immunotherapy. Some factors derived from tumor cells or from within the tumor micro-or macro-environment, including neoantigen load, diversity of the immune infiltrate, expression of immune checkpoints, and the enrichment of prognostic immune gene signatures. Turning cold non-inflamed tumor into hot inflamed tumor is critical for successful immune checkpoint inhibitors. An effective response to immune checkpoints inhibitors has been correlated with inflamed tumor that is characterized by infiltration of T-cells, the presence of IFNγ in tumor microenvironment and the expression of PD-L1 (150). Conversely, there is a lack of therapeutic efficacy response to immune checkpoints inhibitors in non-inflamed tumor (29). One approach to achieve a durable clinical response in patient with cold tumor is the use of collaborative agents to create immunogenic tumor microenvironment (151). Much has been reported on the use of anti-PD-L1 and anti-PD-1 in TNBC (reviewed in ((16)). In March 2019, anti-PD-L1 received accelerated approval to be used in recurrent or unresectable PD-L1 positive metastatic TNBC, based upon a landmark clinical trial (152). In 2021, the FDA granted full approval to anti-PD-1 in metastatic TNBC whose tumors expressed PD-L1(153–155). PD-L1 is a surrogate biomarker used in clinical trials to predict the effectiveness of immune checkpoints inhibitors (115), indicating that intrinsic PD-L1 expression can contribute to treatment efficacy (reviewed in (14)). While there are numerous applications for anti-PD-1 and anti-PD-L1, a sobering reality is that Roche (anti-PD-L1 Atezolizumab) voluntarily withdrew the indication for TNBC in August 2021 (press release https://www.roche.com/media/releases/med-cor-2021-08-27.htm) with little improvement to overall survival (19). Anti-PD-1 remains in use for local and metastatic TNBC (153, 155, 156), though ICIs side effects were noted. While ICIs remain an option for TNBC patients, the uneven clinical results indicate that some improvement is necessary to maximize efficacy and minimize side effects in current regimens.

The mechanisms and frameworks unveiled in the current paper may provide useful insights for improving ICIs for TNBC. A minimal requirement for a collaborative agent is a robust induction of PD-L1 and possibly other markers, but compounds that re-program and activate the tumor-resident environment might be effective in setting an environment for ICI’s to act at lesser levels. The surprising finding from our work and the literature is the prevalence of a tumor re-programming towards MHCs expression and antigen presentation through a viral mimicry to IFN mechanism (see above). Epigenetic inhibitors have emerged as candidates to improve cancer immunotherapy because of the wide impact of global hypomethylation and on PD-L1 status (47). The reprogramming mechanisms described above is also critical. A recent screen focused on epigenetic inhibitors with the express function o inducing antigen presentation properties in TNBC organoids to express antigen presentation machinery and to create a functional compartment with a with infiltration and active (not exhausted) CD8^+^ T-cells (37).

The properties of CHA1 described in the present work suggest an excellent candidate pre-treatment for TNBC to improve the efficacy of anti-PD-L1 treatment for 4 important reasons. **First**, CHA1 pretreatment robustly induces PD-L1 mRNA and protein to levels that would predict an efficacious ICIs response (S7 Fig, Fig 6A, Fig 6B, Fig 7C, Fig 7G). This robust induction is consistent with an epigenetic and IFN mechanism, both of which are orchestrated by CHA1 (Fig 4D and Fig 4E, S4 Fig). However, the initially approved and then withdrawn studies with anti-PD-L1 included a test for PD-L1 expression. Inspection of the original clinical trial data revealed that those tumors that already had PD-L1 expression were more responsive to anti-PD-L1 (147, 152). An assessment of PD-L1 status was part of the FDA approval for anti-PD-L1 and anti-PD-1. Despite the focus on PD-L1, the use of anti-PD-L1 was still withdrawn by Roche in 2021, suggesting that PD-L1 status might be necessary but not sufficient for efficacy in TNBC, in contrast to other cancers such as melanoma and lung where PD-L1 status is widely used as a clinical decision marker. Other criteria may be required for TNBC and would be useful in assessing pre-treatment regiments. In other words, what else in addition to PD-L1 induction is required for efficacy in TNBC? Second, our studies provide criteria besides PD-L1 for possibly improving efficacy. Recent evidence suggested that upregulation of PD-L1 expression was not the only biomarker to the response into immune checkpoint inhibitors. The result of Phase 1b clinical trials of anti-PD-1of 9 different types of cancer found that T-cells inflamed gene signature consists of 18 genes that associated with increased the response into anti-PD-1 (115). Our data showed that in addition to PD-L1, CHA1 induced the expression of T-cell inflamed signature. It has been reported that this gene signature was associated with prolonged the survival (115). This was consistent with our data in invasive and TNBC patients. Figs. 6C, 7G and S7 Fig. reveals additional CHA1 gene-based criteria to augment PD-L1 for predicting ICI efficacy, namely parts of an inflamed immune t-cell and IFN signature (157). We created a derivative signature to assess T-cell inflammation in the syngeneic pre-clinical models. Intriguingly this CHA1-induced composite signature predicted an improved prognosis, when assayed in the TCGA and METABRIC database for breast cancers. Thus, CHA1 treatment leads to robust expression of PD-L1 and other criteria that predict a more favorable outcome in silico. Third, CHA1 pre-treatment leads to the formation of an active tumor-resident CD8^+^ T-cell population, setting the stage for improved anti-PD-1 and anti-PD-L1 efficacy. Fig 6A shows that CHA1 pre-treatment diminishes immune checkpoints PD-1 and CTA-4, considered indicators of T-cell exhaustion, (while increasing PD-L1 in tumors and other immune cells such as macrophages and dendritic cells). The number of Ki67^+^CD8^+^ T-cells increased with CHA1 treatment, while the number of tumor Ki67^+^ cells decreased. Thus CHA1 has reciprocal and beneficial effects on the tumor and tumor-resident T-cells. The activated CD8^+^ T-cells are major factor for ICI efficacy and this activation is one function of ICIs. Again, it is the dilution of Wnt signaling and activation of IFN signaling by CHA1 pre-treatment that likely facilitates this tumor and tumor-resident T-cell phenotype. Fourth, CHA1 does improve anti-PD-L1 in the TNBC preclinical model (Fig 7). CHA1 treatment increased anti-PD-L1 effects in reducing tumor growth and weight, when compared to anti-PD-L1 alone or CHA1 alone. In our assays, we used minimal anti-PD-L1 because of a known hypersensitive reaction in the 4T1 Balb/c models (see results). Even with the model limitations (see results), we could still detect a collaborative effect. Fifth, CHA1 pre-treatment may also reduce ICI dosage needed because the tumor site may be primed. CHA1 is built from compounds with few side effects or with known side effects and that have been use or FDA-approved for other cancer. Similarly, reducing the efficacious dose of ICIs needed for efficacy may lessen the known ICI toxicities (158, 159). Together, we think the evidence is compelling to test CHA1 to improve anti-PD-L1 efficacy.

In surveying the recent literature, we note that the LSD1 inhibitors have a similar and useful property for inducing APC properties in tumor cells and reducing tumor growth, while oppositely recruiting and activating the local T-cell environment (37, 40, 133). Future phase 1 clinical trials in human patients will be required to build upon the preclinical data with CHA1 and other epigenetics disruptors like LSD1 inhibitors. The work in this paper provides insights with CHA1 for enacting a complex cold-to-hot transition which is built upon an exceptionally complex epigenetic network with both reciprocal Wnt and IFN signaling, causing a wholesale re-programming of the tumor and its tumor-resident environment. Our studies here set a solid foundation for thoughtful compound development to improve the current ICIs and look towards newer immune checkpoint drugs.

## Acknowledgements

Drs. Amy Yee and K. Eric Paulson are co-founders of Cha Therapeutics, Inc. and of U.S. National Patent Application No.: 17/293,648, both of which will develop this work for future commercial applications. The studies in this paper were supported through the years by grants to ASY from the National Cancer Institute (CA 104236, CA94187), Dept. of Defense (BC121274), American Institute for Cancer Research, and the Susan B. Komen Foundation (KG080816; KG100913), Hackensack Research Foundation, and Cha Therapeutics.

Ahlam M. Bogis, Roaya S. Alqurashi, and Mariam A. Alamoudi were supported by graduate fellowships from the Saudi Arabian Cultural mission. MA was additionally supported by her the Pharmacology and Toxicology Department, College of Pharmacy, Prince Sattam bin Abdulaziz University, AlKharj, Saudi Arabia Roaya Alquarashi and Ahlam Bogis acknowledge support from the Pharmacology and Toxicology Department, College of Pharmacy, Umm Al-Qura University, Makkah, Saudi Arabia.

ASY especially acknowledges and appreciates Nobel Laureate and Prof. Bill Kaelin for his insightful critique of this study at an early stage (after discovery of the Wnt signaling mechanisms and for encouraging further investigations that led to the discovery of the new findings on tumor immunity. Had Dr. Kaelin not provided his useful comments, this study would have stopped et Figure 2. We acknowledge Drs. Gail Sonenshein and Nora Minerva for providing the Luciferase/GFP-expressing MDA-MB-231 cells for the human xenograft experiments and for many helpful discussions on green tea. We additionally acknowledge previous students from the Pharmacology graduate programs (Mohammed Alshagawi, Salwa Hafez), Biomedical sciences program (Helen Uong, Zixu Wang, Ashley Waplinger), and talented Tufts university undergraduate (Anise Applebaum, Eileen Liu), and a Saudi Arabian medical student exchange program (Ismail Shakir, Lamma Hassan)—for all their numerous contributions to the work of this paper.

## Paper contributions

This project was conceived, developed, and supervised by ASY and KEP, providing all the intellectual framework. The other members of the Yee lab, past and present, contributed many suggestions and experimental refinement during our interactive lab meetings. ASY and KEP wrote and finalized the manuscript. MA wrote the first draft as her PhD. thesis, revised later drafts, and assembled their data into figures. MC and FDF are talented Tufts University undergraduates provided extensive contribution to the project design and execution in the early stages as part of honors thesis research. The remaining authors all performed critical experiments, provided critical data that underlie all the figures, and which became part of their master’s or undergraduate theses.

## Materials and Methods

### Cell Lines

#### Human cell lines

A derivative of MDA-MB-231 that express both luciferase and GFP, a gift from Dr. Gail Sonenshein, was used for the TNBC human xenograft experiment.

#### Mouse Cell Lines

4T1 cell lines, purchased from ATCC, were used for the TNBC syngeneic model.

### Mutant Cell Construction in Human Cell Lines

#### Lentivirus knockdowns

LM1 cells, a derivative of MDA-MB-231 cell lines (164), were transfected with lentiviral particles. Two shRNA constructs targeting sFRP1 and HBP1, as well as empty vector pLKO control were used. The lentiviral transduction protocol from Sigma-Aldrich was used for viral transfection. 8 μg/ml of hexadimethrine bromide was added for enhancing transduction. Cells were selected with 1 to 2 μg/ml of puromycin. Knockdown of HBP1 and sFRP1 were confirmed with qRT-PCR by comparing the gene knockdown constructs to control.

### Treatment Regimen for *in vitro* Study

DAC and EGCG were dissolved in PBS and ethanol with PBS, respectively. Cells were treated with 1μM of DAC + 10μM of EGCG (combination treatment), 1μM of DAC, 10μM of EGCG, and no treatment. Treatment was conducted for 96 hours. All cells were maintained in the normal media that contained Dulbecco’s Modified Eagle Medium (DMEM) supplemented with 10% fetal bovine serum (FBS), antibiotics, and anti-trypsin during treatment. Media and drugs were replaced every 24 hours (four times during the 96-hour treatment period).

### Animals

Six-to eight-week-old female NOD/SCID mice and Balb/c mice were purchased from Jackson Laboratory. Mice were housed in a sterile environment and maintained by DLAM (Division of Laboratory Animal Medicine) personnel in accordance with guidelines established by the Institutional Laboratory Animal Care and Use Committee (IACUC) at Tufts University-Tufts Medical Center.

### Cell Preparation for Animal Surgery

Cells were cultured in the normal growth media (DMEM supplemented with 10% FBS, penicillin (100 units/ml), streptomycin (0.1 mg/ml), and 10 mM L-glutamine) and allowed to grow to no more than 85% confluence. To harvest cells, cells were initially trypsinized with 0.05% trypsin and subsequently washed with PBS. The cells were then re-suspended in serum-free, antibiotic-free Dulbecco’s Modified Eagle Medium. To determine cell concentration for accurate inoculation into xenograft models, cells were stained with 0.4% trypan blue solution (Sigma) and counted using a hemocytometer. A 1:1 volume ratio of Matrigel (BD Biosciences) was used to set the cells for implantation/inoculation into the mammary fat pad of each individual mouse.

### Animal Surgery

All animal surgeries were carried out in a procedure room under a sterile environment following guidelines for animal surgery establishment by Tufts Medical Center’s Division of Laboratory Animal Medicine. Animals were initially anesthetized in an isoflurane chamber. During surgery, mice were administrated a steady flow of isoflurane through a nose conical. *Surgery:* area around the 4^th^ mammary fat pad was prepared by shaving hair and sterilizing with iodine and alcohol. Care was taken to locate the 4^th^ mammary fat pad and an incision was made according to expose the area for implantation of cancer cells. Connective tissues were released to fully access the area for the inoculation of cells. A total volume of 30 μl or 35 μl containing 3 x 10^5^ -5 x 10^5^ cells suspended in a 1:1 ratio of serum-free, antibiotic-free media and Matrigel was injected into the mammary fat pad using a sterile Hamilton Syringe. Following inoculation of cells, the incision was repaired using wound clips. The animals were allowed to recover from anesthesia under close monitoring. Buprenorphine was used as an analgesic (0.06 mg/kg). Animals were monitored for three days post-surgery for proper healing and any post-operative complication. Wound clips were removed between 10 and 14 days post surgery. For tumor resection, mice were anesthetized and opened as described here, and the tumors (0.4 – 0.5 cm diameter) carefully revealed. The three main blood vessels leading to the mammary gland/tumor were heat cauterized to prevent bleeding and the tumor carefully excised. The wounds were closed as described here with wound clips.

### Drug Treatment Plan for *in vivo* Study

When tumors reached approximately 0.125 cc, treatments were initiated. Both EGCG (Sigma-Aldrich) and DAC (Sigma-Aldrich) were dissolved in PBS and then filter sterilized. A 28-gauge insulin syringe was used for intraperitoneal injection (i.p). All treatments continued for the indicated time unless tumor size exceeded that allowed by IACUC guidelines or if the animal’s health appeared to be deteriorating. Treatment plan was carried out in human xenograft model with MDA-MB-231 tumors, sFRP1 knockdown MBA-MB-231 tumors, and HBP1 knockdown MBA-MB-231 tumors implanted into NOD/SCID mice, as well as in syngeneic model with 4T1 tumors implanted into Balb/c mice.

#### CHA1 QOD treatment protocol in TNBC human xenograft and TNBC syngeneic models

mice were randomly assigned to either control or experimental groups with at least five mice in each group. Mice in the CHA1 experimental group were injected i.p. on alternating days with either 16.5 mg/kg EGCG or 0.5 mg/kg DAC. Control mice were injected i.p. with 100 µL of saline each day.

#### EGCG QOD monotherapy in TNBC human xenograft

mice were injected i.p. on alternating days with 16.5 mg/kg of EGCG.

#### DAC QOD monotherapy in TNBC human xenograft

mice were injected i.p. on alternating days with 0.5 mg/kg of DAC.

#### CHA1 QD treatment in TNBC syngeneic models

mice were randomly assigned to either control or experimental groups with at least five mice in each group. 16.5 mg/kg of EGCG and 0.5 mg/kg of DAC were administrated concomitantly via i.p. injections during cycle 1 and cycle 2. The duration of each cycle was 5 days. There was 5 days recovery period between the cycles.

#### CHA1 + ICIs treatment in TNBC syngeneic models

mice were randomly assigned to either control or experimental groups with at least eight mice in each group. CHA1 QD treatment protocol was followed. Mice in ICIs treatment group and CHA1+ ICIs group received two doses (100 μg/mouse) of anti-PD-L1 antibody (clone 10F.9G2, Bio X cell). The doses were two days apart and administrated during the recovery period between the cycles of CHA1 treatment. The first dose of ICIs was administrated via i.p. injections on the first day of the recovery period and the second dose was administrated via i.p. injections on the fourth day of the recovery period.

### Monitoring of Tumor Growth: Caliper Measurements

After cells implantation, animals were palpated for evidence of tumor. Once a measurable tumor was noted, caliper measurements were taken every other day. The two longest dimensions of the tumor were used to calculate tumor volume (the formula width² x length was used to determine tumor volume).

### Bioluminescence Imaging for *in vivo* Luminescence

Bioluminescence imaging in vivo luminescence was accomplished using the Tufts Imaging Core IVIS Imaging System (Xenogen). MDA-MB-231 cells used contain a luciferase gene construct that enabled visualization of live cells upon administration of luciferin (0.1 mg/mouse). Animals were anesthetized in an isoflurane chamber after i.p. injection of luciferin and remained so through a nose conical during the entire imaging procedure. Acquisition time was 60 seconds when tumors were small and signal strengths were low. As tumor size increased, acquisition time was reduced to 30 seconds to avoid saturation. Analysis was performed using Living Image software (Xenogen).

### Termination of *in vivo* Study

All animals were carefully monitored to ensure that tumor size was within limits established by IACUC and the animals were in good health. After euthanasia of animal, an autopsy was performed to look for any evidence of metastases and abnormalities. Suspected tissues were harvested for analysis. The primary tumor was harvested from each mouse and fixed with formalin. Fixed tissues were preserved in 70% ethanol. A small, unfixed sample from each mouse was stored at −80°C for further analysis.

### Immunohistochemistry and Immunofluorescence

Immunostainings of β-catenin, MHC-I and MHC-II of human xenograft tumor samples were performed by Tufts University Core Facility (Boston, MA). Immunostainings of Ki67β-catenin, PD-L1, CD8, F480, and FOXP3 of syngeneic tumor samples were performed by iHisto (Salem, MA). Cell membrane immunostaining for control and treated tumors was assessed by three investigators in a blinded fashion. The Allred scoring method was used as previously described for assessing the membrane proportion and staining intensity in scale between 0 to 8 (160). The score was calculated from the sum of a proportion score (0-5 scale reflecting the percentage of cells with any stain) and a staining intensity score (0-3 scale reflecting the intensity of staining among the positive cells) (160). A score between 0 and 2 indicates negative staining and a score between 3 and 8 indicates positive result (160). Immunofluorescence staining of Ki67 and CD8 of syngeneic tumor samples was performed by iHisto (Salem, MA).

### Total RNA Extraction and cDNA Synthesis

For in vitro experiment: cells were harvested for total RNA extraction after completion of treatments. For in vivo experiment: a small sample of tumor tissue was used for total RNA isolation. RNA extraction was performed using RNeasy Plus Mini Kit (Qiagen) and was eluted with RNase-free water. Nanodrop spectrophotometer (Thermo Scientific) was used to measure RNA concentration by measuring absorbance at 260nm. After determining RNA concentration, 500 μg of RNA was used to synthesize cDNA using iScript cDNA Synthesis Kit (Bio-Rad).

### qRT-PCR

cDNA synthesized from cells and animal tumors tissues was used for real-time PCR performed on iCycler (Bio-Rad) using a 2x SYBR Green master mixture (Bio-Rad) in a 20 μl reaction. All transcript quantitation was normalized to endogenous 18S RNA control. The relative quantitative value for each target gene compared with the calibrator for the target was expressed as comparative Ct (2^-(ΔCt-Cc)^) method (Ct and Cc are the mean threshold cycle differences after normalizing to 18S). qRT-PCR experiments were performed in triplicates. Primer sequences were presented in S10 Table and S11 Table. Human primers for HLA-DRB1 and VCAM1 were purchased from Bio-Rad.

### Protein Extraction

Cell and tumor lysates were lysed for 30 minutes at 4°C with RIPA buffer (Thermo Scientific) that mixed with protease and phosphatase inhibitor cocktail (100X) (Thermo Scientific Halt^TM^ kit) and 0.5 M EDTA (Thermo Scientific) (1:100 dilution). The lysate was centrifuged for 10 minutes at 14,000 rpm at 4°C. The supernatant was kept and frozen at −80°C. Protein concentration measurement was done following Thermo Scientific Quant-iT^TM^ Assay. Protein concentration was measured with Qubit^TM^ fluorometer (Thermo Scientific).

### Western Blotting

30 μg protein extracts were separated on 7.5% gradient polyacrylamide gel (Mini-PROTEAN TGX gel, Bio-Rad) by electrophoresis (SDS-PAGE). Then, proteins were transferred to PVDF membrane (Bio-Rad), washed with PBST (1xPBS, 0.1% Tween) and blocked with 5% BSA solution for at least one hour at 4°C. Once the membrane is done blocking, primary antibodies were incubated overnight at 4°C. The list of primary antibodies used was presented in S12 Table. The western blot results were developed with SuperSignal West Femto Chemiluminescent substrate (Thermo Scientific) and quantitated using a Chemidoc MP Imaging System (Bio-Rad).

### RNA Sequencing

RNA from control and CHA1 treated tumors of human xenograft was sequenced at the Tufts Core Facility-TUCF Genomics (Boston, MA). Total RNA from three saline treated tumors and three CHA1 treated tumors were submitted and 25-28 million human mRNA reads were analyzed for each sample. The data were analyzed using IPA (QIAGEN Inc., https://www.qiagenbioinformatics.com/products/ingenuity-pathway-analysis). GSEA analysis was done using Molecular Signatures Database (MSigDB) (161). MSigDB REACTOME gene set was used for the analysis.

### Human Cancer Transcriptome Analysis

The data were generated using the cBioPortal for Cancer Genomics (http://www.cbioportal.org) (162, 163). Data were obtained from the following breast cancer dataset: “METABRIC, Nature 2012 & Nat Commun 2016 (n=2509)” (83–85), “TCGA, PanCancer Atlas (n=1084)“ and “TCGA, Cell 2015 (n=818)” (82). mRNA expression relative to all samples with z-score threshold of 1.3 was used to extract the data.

### Statistical analysis

All experiments were performed in triplicate. Student’s unpaired *t*-test was used for comparison between two groups. One-way ANOVA with Tukey’s post-hoc test was used for multiple comparison. Manne-Whitney U-test was used to compare between tumor volumes and tumor weights. Kaplan-Meier survival curve with the hazard ratio (HR), 95% confidence intervals (CIs) and log-rank *P-*value was used for survival analysis. A *P-*value less than 0.05 was considered statistically significant. All analyses were performed by GraphPad Prism 9.

---

**S1 Figure: The mechanistic rationale for combining EGCG and DAC to inhibit Wnt signaling. A)** There was a negative correlation between DNMT1 mRNA expression and HBP1 mRNA expression. The co-expression data was obtained from cBioPortal for Cancer Genomics. “TCGA, Cell 2015” (82) and “METABRIC, Nature 2012 & Nat Commun 2016” (83–85) were used to extract the data (refer to materials and methods). **B, C)** CHA1 inhibited Wnt signaling in vitro in MDA-MB-231 cells. CHA1 significantly induced the mRNA levels of Wnt inhibitors **B)** sFRP1 and **C)** HBP1 at higher levels than the levels induced by monotherapy. **D-G)** sFRP1 and HBP1 were necessary for CHA1 function in vitro. The effect of CHA1 on basal and induced sFRP1 and HBP1 mRNA levels were decreased in sFRP1 KD MDA-MB-231 cells and HBP1 KD MDA-MB-231 cells compared to control empty vector MDA-MB-231 cells (pLKO). **B-H)** The gene expression of Wnt inhibitors sFRP1 and HBP1 was measured by qRT-PCR in empty vector MDA-MB-231 cells (pLKO), sFRP1 KD, and HBP1 KD MDA-MB-231 cells treated with control, 10 µM EGCG or 1 µM DAC or CHA1. **B, C)** Unpaired two-tailed t-test was used for the comparison between two groups. **D-G)** Adjusted one-way ANOVA was used for multiple comparison. Data were represented as mean ± SEM. *\* P < 0.05, ** P < 0.01, *** P < 0.005, **** P < 0.001*.

**S2 Figure: CHA1 treatment efficacy in TNBC human xenograft and TNBC syngeneic mouse model. A)** CHA1 QOD treatment reduced the primary tumor size as seen by luciferase imaging. Orthotopically implanted MDA-MB-231 luciferase-tagged cell tumors grew for approximately 3 weeks with or without CHA1 treatment. Mice were anesthetized with isofluorane, injected i.p. with 90 μl of luciferin and luciferase expression activity imaged using an IVIS Spectrum CT Biophotonic Imager (Tufts Small Animal Imaging Preclinical Testing Facility). **B)** Knockdown of sFRP1 and HBP1 promoted tumor growth. Empty vector control (pLKO), sFRP1 KD, and HBP1 KD MDA-MB-231 cells (S1 Fig) were implanted orthotopically into NOD/SCID mice. Tumor growth was monitored by caliper measurement for the indicated time (n=6-7). **C-D)** sFRP1 and HBP1 were necessary for CHA1 efficacy in TNBC human xenograft model. The inhibitory effect of EGCG only, DAC only, or CHA1 treatment on tumor growth was tested on **C)** MDA-MB-231 sFRP1 KD cell (S1Fig) tumors (n=3) and **D)** MDA-MB-231HBP1 KD cell (S1 Fig) tumors (n=3). Tumor volume was measured with calipers at indicated times. **E-H**) CHA1 induction of sFRP1 and HBP1 was reduced in sFRP1 KD and HBP1 KD tumors. RNA was prepared from empty vector control (pLKO), sFRP1 KD, and HBP1 KD MDA-MB-231 tumors from Fig 1C, S2C Fig and S2D Fig receiving CHA1, EGCG or DAC QOD monotherapy. Wnt inhibitors sFRP1 and HBP1 mRNA levels were measured using qRT-PCR (n=3-6). **I, J)** CHA1 QOD treatment suppressed overall metastases **(I)** and brain metastases **(J)** in TNBC human xenograft model. The total and brain metastases were assessed after CHA1 treatment. Orthotopic implants of MDA-MB-231 cells, dually tagged with luciferase and GFP (gift of Drs. G Sonnenshein and N. Mineva), were treated with CHA1 or saline for 2 weeks, followed by resection of the tumor with continued saline or CHA1 treatments. Large metastases were identified by luciferase imaging **(I)**. Smaller metastatic foci were identified in liver, lung or brain by GFP imaging **(J)**. One mouse with a somewhat larger brain tumor presented with seizures. **K)** H&E staining of untreated and treated tumor demonstrated less cellularity after CHA1 QOD treatment in TNBC syngeneic mouse model. **L)** immunohistochemistry of Ki67 indicated decreased cell proliferation after CHA1 QOD treatment in TNBC syngeneic mouse model. Different sections of different untreated and treated tumors were stained with anti-Ki67 antibody. Representative immunostaining section with quantification of the percentage of Ki67 positive cells in control tumors and CHA1 QOD treated tumors (n= 3 mice/ group). **M)** CHA1 QD cycle 1 treatment significantly decreased tumor volume in TNBC syngeneic mouse model (n=15-23). In addition, tumor weight was reduced after CHA1 QD cycle 1 treatment in TNBC syngeneic mouse model (n=5). 4T1 cells were implanted orthotopically into Balb/c mice. Tumor volume was measured with calipers. Tumors were weighted after CHA1 cycle 1 treatment. The tumor volume data were combinations of 3 different studies. **E, F, J)** Unpaired two-tailed t-test was used for the comparison between two groups. **E-H)** Adjusted one-way ANOVA was used for multiple comparison. **B, C, D, H)** Mann-Whitney U-test was used for tumor volume and tumor weight comparisons. Data were represented as mean ± SEM. *\* P < 0.05, ** P < 0.01, *** P < 0.005, **** P < 0.001*.

**S3 Figure: The effect of CHA1 on Wnt signaling in TNBC human xenograft. A)** IHC of β-catenin after saline and CHA1 QOD treatment in TNBC human xenograft tumors demonstrated an intense membrane staining of β-catenin in saline group. Representative immunostaining section of β-catenin after saline and CHA1 QOD treatment in TNBC human xenograft model. **B)** CHA1 significantly downregulated the gene expression of Wnt target genes AXIN2 while EGCG QOD monotherapy and DAC QOD monotherapy showed no inhibitory effect. **C)** The gene expression of sFRP1 was significantly upregulated after CHA1 QOD, EGCG QOD monotherapy, and DAC QOD monotherapy treatment in TNBC human xenograft (n= 5-6 mice/group). The gene expression of HBP1 was significantly upregulated after CHA1 QOD and DAC QOD monotherapy treatment in TNBC human xenograft (n= 5-6 mice/group). For Figures **B** and **C**, the gene expression was tested by qRT-PCR after saline, EGCG, DAC and CHA1 treatment (n= 5-6 mice/group). Adjusted one-way ANOVA was used for multiple comparison. qRT-PCR experiments were done in triplicates in three independent experiments. Data were represented as mean ± SEM. *\* P < 0.05, ** P < 0.01, *** P < 0.005, **** P < 0.001*.

**S4 Figure: CHA1 efficacy on activation of antigen presentation and IFN pathways in TNBC human xenograft. A)** CHA1 QOD and DAC QOD monotherapy significantly induced the gene expression of MHC-I (HLA-B). The gene expression of MHC-II (HLA-DRB1 and HLA-DRB5) was significantly induced after CHA1 QOD treatment compared to EGCG QOD monotherapy or DAC QOD monotherapy. **B)** CHA1 QOD significantly upregulated the gene expression of CTAs compared to EGCG QOD monotherapy or DAC QOD monotherapy. **A, B)** The gene expression was measured by qRT-PCR in saline, EGCG, DAC and CHA1 treatment (n= 4-6 mice/group). **C)** pSTAT3^Y705^ protein levels were significantly elevated after CHA1. pSTAT3^Y705^ protein levels were unchanged after EGCG QOD monotherapy or DAC QOD monotherapy. Representative western blot (left) with quantification (right) of saline, EGCG, DAC and CHA1 treatment tumors (n=4 mice/ group). **D)** The overlapping genes between RNA-seq of CHA1 treated tumors and GSEA MSigDB REACTOME IFNα/β and IFNγ gene sets. A hypergeometric probability test was utilized. The induction of at least 8 genes in each interferon pathway was required to validate stimulation of such pathway. **E)** CHA1 QOD treatment significantly upregulated ISGs. DAC QOD monotherapy also significantly increased the expression of ISGs except IFIT1 and IFIT3, however, the fold change in the gene expression was higher after CHA1 QOD treatment. EGCG QOD monotherapy had no stimulatory effect on ISGs. The gene expression was tested by qRT-PCR after saline, EGCG, DAC and CHA1 treatment (n= 4-6 mice/group). **F)** STAT3 phosphorylation status (Fig 4D) was positively correlated with mRNA levels of OAS1, OAS2, OAS3, OASL, IFIT1, IFIT2, and IFIT3 (Fig 4E) after CHA1 QOD treatment in TNBC human xenograft model. r^2^ value was calculated using GraphPad Prism. **A-C, E)** Adjusted one-way ANOVA was used for multiple comparison. qRT-PCR experiments were done in triplicates in three independent experiments. Western blot quantification was the combination of two different experiments. Data were represented as mean ± SEM. *\* P < 0.05, ** P < 0.01, *** P < 0.005, **** P < 0.001*.

**S5 Figure: CHA1 reprogramming effects on tumor cells, immune cells, and biological functions in TNBC human xenograft and syngeneic models. A)** CHA1 QOD treatment increased expression of human PRF1 (n=5-6) in TNBC human xenograft tumors, indicating reprogramming of tumor cells to express gene indicative of NK cells and T-cells activation. **B)** CHA1 QOD treatment activated NK cells via upregulation of the gene expression of mouse Prf1 and mouse Gzmb (n=4-5) in TNBC human xenograft tumors. **A, B)** The gene expression was measured by qRT-PCR in untreated and treated tumors. **C, D)** CHA1 reversed epithelial to mesenchymal transition. **C)** CHA1 QOD treatment significantly increased the protein expression of E-cadherin in TNBC human xenograft and TNBC syngeneic mouse model. Representative western blot (left) with quantification (right) of saline and CHA1 treatment tumors (n=4 mice/ group). **D)** Immunohistochemistry of E-cadherin in saline and CHA1 QOD treatment in TNBC human xenograft tumors indicated increased E-cadherin protein expression after CHA1 QOD treatment. Representative immunostaining section of E-cadherin after saline and CHA1 QOD treatment in TNBC human xenograft model. **E)** Changes in biological functions as a result of CHA1 QOD treatment. For example: CHA1 suppressed cellular movement and cellular growth (blue), as well as stimulation of cell death (orange). **A)** Adjusted one-way ANOVA was used for multiple comparison. **B)** Unpaired two-tailed t-test was used for the comparison between two groups. qRT-PCR experiments were done in triplicates in three independent experiments. Western blot quantification was the combination of two different experiments. Data were represented as mean ± SEM. *\* P < 0.05, ** P < 0.01, *** P < 0.005, **** P < 0.001*.

**S6 Figure: CHA1 effect on immune cells. A)** H&E staining of untreated tumor and CHA1 QOD-treated tumor in TNBC syngeneic model revealed the development of lymphoid structure. **B, C)** Immunohistochemistry of **B)** CD8^+^ T-cells and **C)** F480^+^ cells after CHA1 QOD treatment in TNBC syngeneic model indicated the presence of lymphoid structure. **D)** Immunohistochemistry of CD8^+^ T-cells indicated increased infiltration of cytotoxic CD8^+^ T-cells after CHA1 QOD in TNBC syngeneic model. Different sections of different untreated and treated tumors were stained with anti-CD8 antibody. Representative immunostaining section with quantification of the percentage of CD8^+^ T-cells in control tumors and treated tumors (n= 5 mice/group). Unpaired two-tailed t-test was used for the comparison between two groups. qRT-PCR experiments were done in triplicates in three independent experiments. Data were represented as mean ± SEM. *\* P < 0.05, ** P < 0.01, *** P < 0.005, **** P < 0.001*.

**S7 Figure: CHA1 effect on PD-L1 expression and T-cell inflamed signature.**

**A)** CHA1 QOD treatment significantly upregulated the gene expression of PD-L1 compared to EGCG QOD monotherapy or DAC QOD monotherapy in TNBC human xenograft model. The gene expression was measured by qRT-PCR in saline, EGCG, DAC and CHA1 treated tumors (n=4-5). **B)** IHC of PD-L1^+^ cells demonstrated significant increase in the protein levels of PD-L1 after CHA1 QOD treatment compared to EGCG QOD monotherapy and DAC QOD monotherapy in TNBC human xenograft. Different sections of different control, EGCG, DAC, and CHA1 treated tumors were stained with anti-PD-L1 antibody. Representative immunostaining section with quantification of the percentage of PD-L1^+^ cells in control tumors and treated tumors (n= 4 mice/group). **C)** STAT3 phosphorylation status (Fig 4D) was positively correlated with mRNA levels of PD-L1 (Fig 6A) after CHA1 QOD treatment in TNBC human xenograft model. r^2^ value was calculated using GraphPad Prism. **D)** The overlapping genes between the published T-cell inflamed signature (115, 116) associated with improved the response to anti-PD-1/anti-PDL-1 and CHA1 gene signature. There were 17 overlapping genes between the two signatures (hypergeometric p-value = 7.7 e-46). **E)** Pdl-1 mRNA levels were positively correlated with mRNA levels of Ifng, Cd8a, Ccl5, Tigit, Lag3, Ido1, and Psmb10 after CHA1 QD treatment in TNBC syngeneic mouse model. Fig 4A, Fig 6A, Fig 6C**, and** Fig 7G were used to generate the data. **F)** PD-L1 mRNA levels (Fig 6A) were positively correlated with mRNA levels of HLA-DRB1 (Fig 4A) after CHA1 QOD treatment in TNBC human xenograft model. **E, F)** r2 value was calculated using GraphPad Prism. **G)** PD-L1 expression was positively correlated with CHA1 T-cell inflamed gene signature including IFNG, CD8A, CCL5, TIGIT, LAG3, IDO1, GZMB, and PRF1 in patients with invasive breast carcinoma. The co-expression data was obtained from cBioPortal for Cancer Genomics. “TCGA, Cell 2015” (82) was used to extract the data (refer to materials and methods). **A, B)** Adjusted one-way ANOVA was used for multiple comparison. qRT-PCR experiments were done in triplicates in three independent experiments. Data were represented as mean ± SEM. *\* P < 0.05, ** P < 0.01, *** P < 0.005, **** P < 0.001*.

**S8 Figure: CHA1 gene signature predicts better prognosis.**

Invasive breast carcinoma patients with high expression of CHA1 gene signature including **A)** PD-L1, **B)** TIGIT, **C)** HLA-B (MHC-I), **D)** HLA-DRB1 (MHC-II), **E)** OAS2 (ISG), **F)** IFI27 (ISG) and **G)** HBP1 (Wnt inhibitor) had a prolonged survival. cBioPortal for Cancer Genomics was used to extract the data. “TCGA, Cell 2015” (82) was used to obtain the survival data for **A, C, E & F**. “TCGA, PanCancer Atlas” was used to obtain the survival data for **B & D**. “METABRIC, Nature 2012 & Nat Commun 2016” (83–85) was used to obtain the survival data for **G**. mRNA expression relative to all samples with z-score threshold of 1.3 was used to extract the data. Kaplan-Meier survival curve with the hazard ratio (HR), 95% confidence intervals (CIs) and log-rank *P*-value was used for survival analysis (refer to materials and methods).

**S9 Figure: CHA1 in combination with ant-PD-L1 in TNBC syngeneic mouse model. A)** H&E staining of saline, anti-PD-L1, CHA1, and CHA1+ anti-PD-L1. **B)** Picture of saline, anti-PD-L1, CHA1, and CHA1+ anti-PD-L1 tumors dissected at the end of the experiment. CHA1+ anti-PD-L1 tumors demonstrated the presence of necrotic area.

**S1 Table -S8 Table: Wnt target genes and immune related pathways altered after CHA1 treatment in TNBC human xenograft model.** The gene set of each pathway was extracted from IPA after analysis of CHA1 RNA-seq in TNBC human xenograft model.

**S9 Table: Biological functions altered after CHA1 treatment in TNBC human xenograft model.** The table was extracted from IPA after analysis of CHA1 RNA-seq in TNBC human xenograft model.

