## Supplemental Figures for "CHA1: A New Combinatorial Therapy That Reciprocally Regulates Wnt and JAK/STAT/Interferon Signaling to Re-program Breast Tumors and the Tumor-Resident Landscape"

A

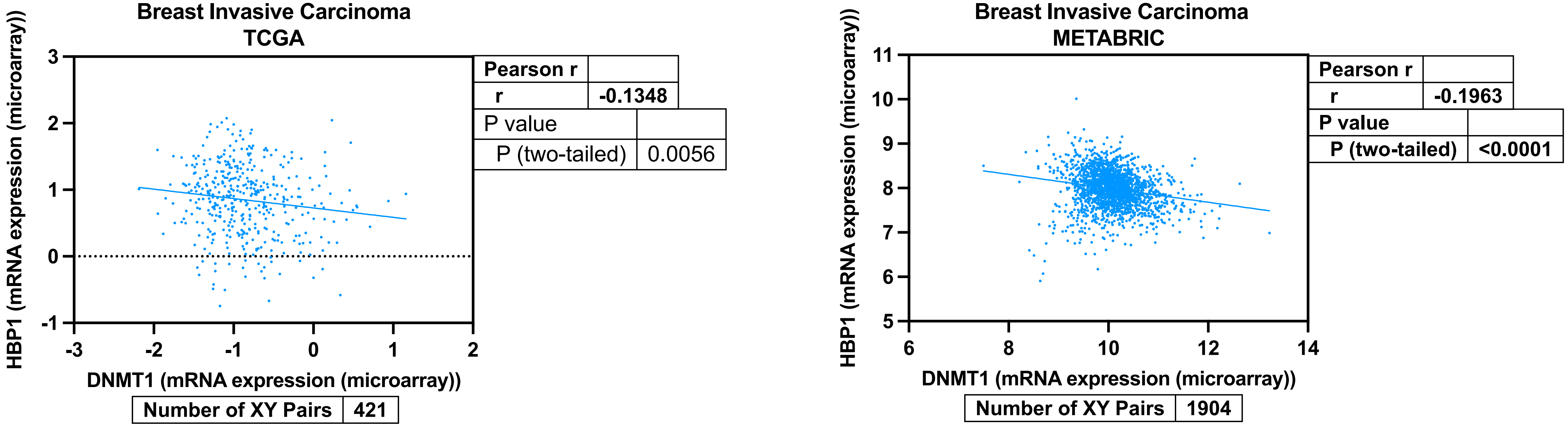
Breast Invasive Carcinoma
TCGA
HBP1 (mRNA expression (microarray))
DNMT1 (mRNA expression (microarray))
Pearson r
r
-0.1348
P value
P (two-tailed)
0.0056
Number of XY Pairs
421
Breast Invasive Carcinoma
METABRIC
HBP1 (mRNA expression (microarray))
DNMT1 (mRNA expression (microarray))
Pearson r
r
-0.1963
P value
P (two-tailed)
<0.0001
Number of XY Pairs
1904

B

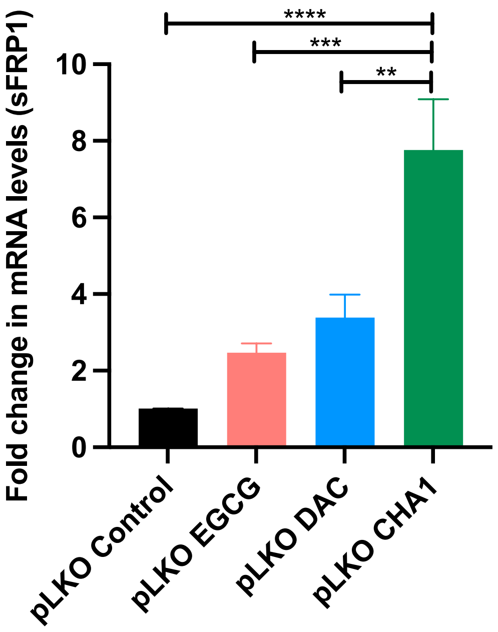

****
***
**
10
8
6
4
2
0
Fold change in mRNA levels (sFRP1)
pLKO Control
pLKO EGCG
pLKO DAC
pLKO CHA1


C

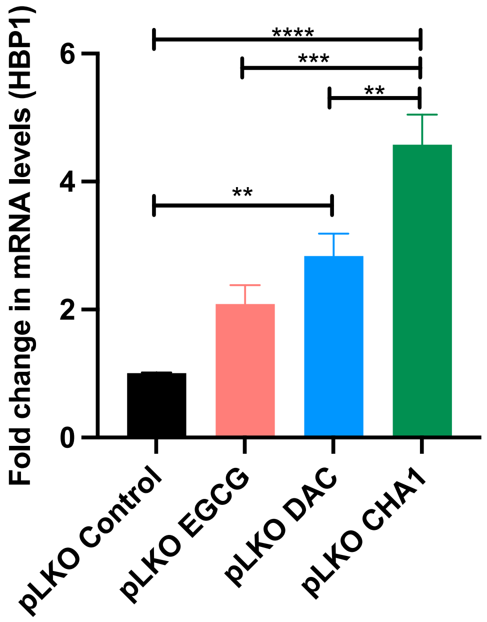

****
***
**
**
6
4
2
0
Fold change in mRNA levels (HBP1)
pLKO Control
pLKO EGCG
pLKO DAC
pLKO CHA1

D

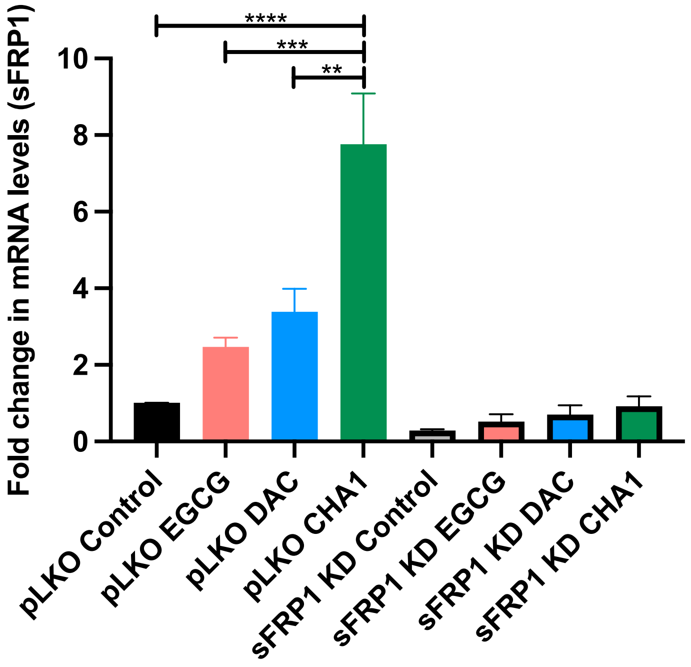

Fold change in mRNA levels (sFRP1)
10
8
6
4
2
0
****
***
**
pLKO Control
pLKO EGCG
pLKO DAC
pLKO CHA1
sFRP1 KD Control
sFRP1 KD EGCG
sFRP1 KD DAC
sFRP1 KD CHA1


E

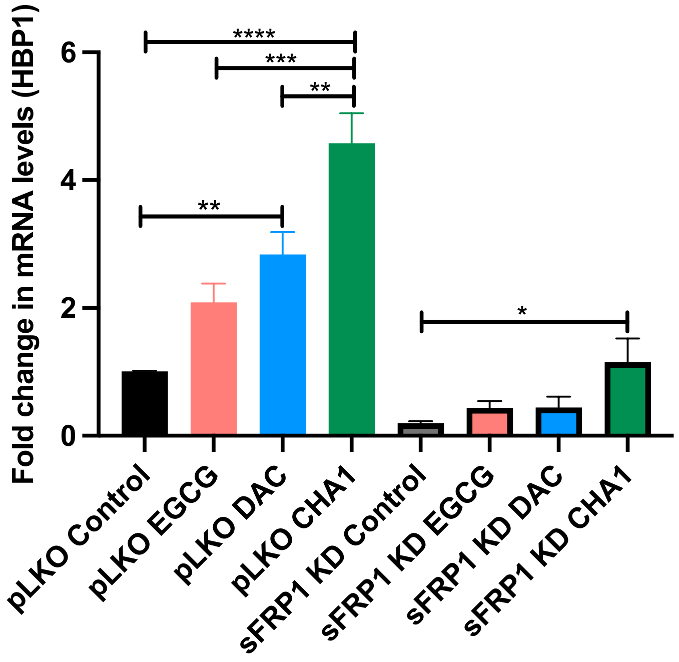

Fold change in mRNA levels (HBP1)
6
4
2
0
****
***
**
**
*
pLKO Control
pLKO EGCG
pLKO DAC
pLKO CHA1
sFRP1 KD Control
sFRP1 KD EGCG
sFRP1 KD DAC
sFRP1 KD CHA1

F

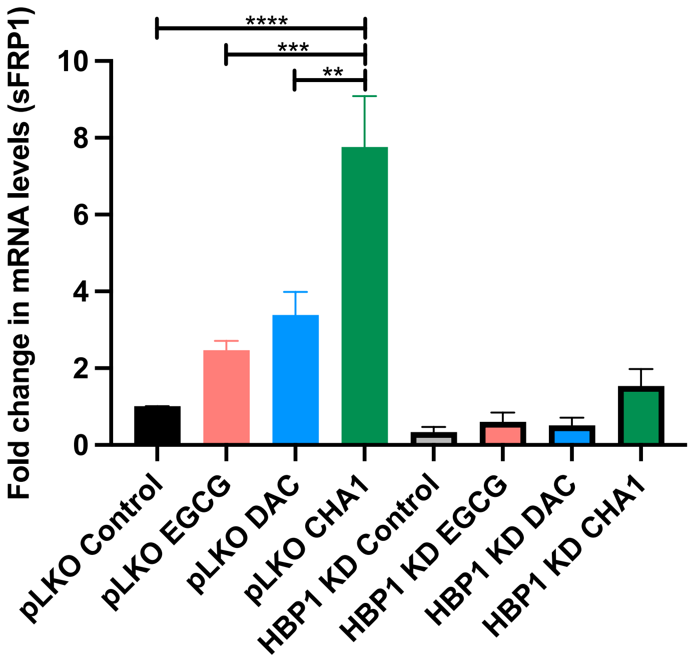

Fold change in mRNA levels (sFRP1)
10
8
6
4
2
0
****
***
**
pLKO Control
pLKO EGCG
pLKO DAC
pLKO CHA1
HBP1 KD Control
HBP1 KD EGCG
HBP1 KD DAC
HBP1 KD CHA1


G

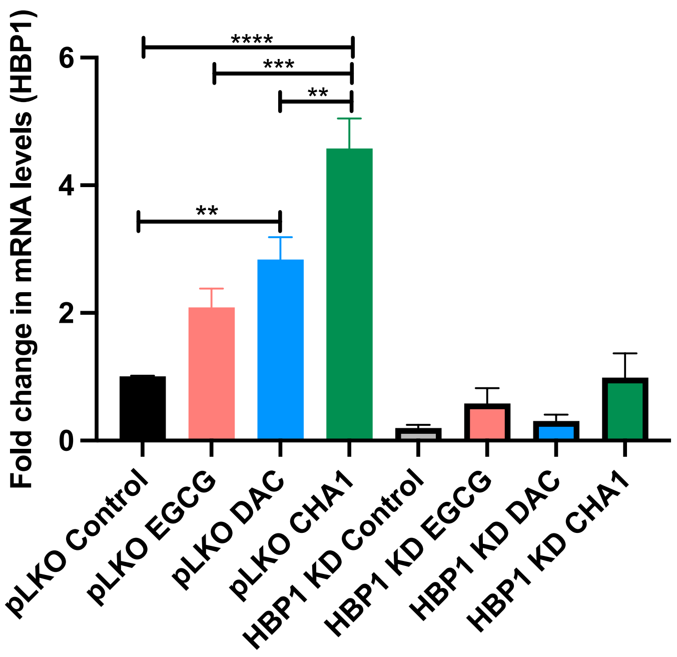

Fold change in mRNA levels (HBP1)
6
4
2
0
****
***
**
**
pLKO Control
pLKO EGCG
pLKO DAC
pLKO CHA1
HBP1 KD Control
HBP1 KD EGCG
HBP1 KD DAC
HBP1 KD CHA1

TNBC human xenograft model
QOD treatment

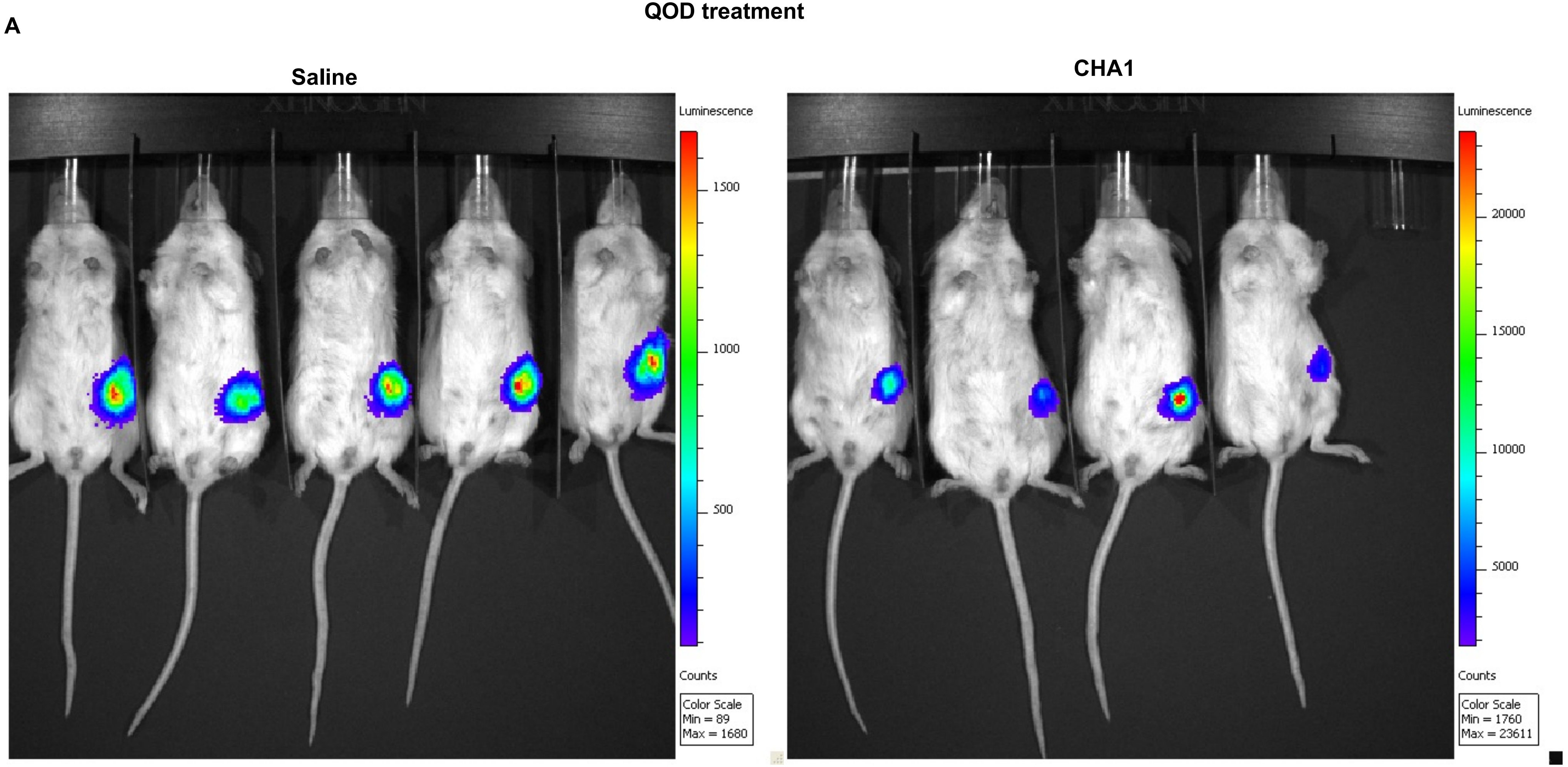

B

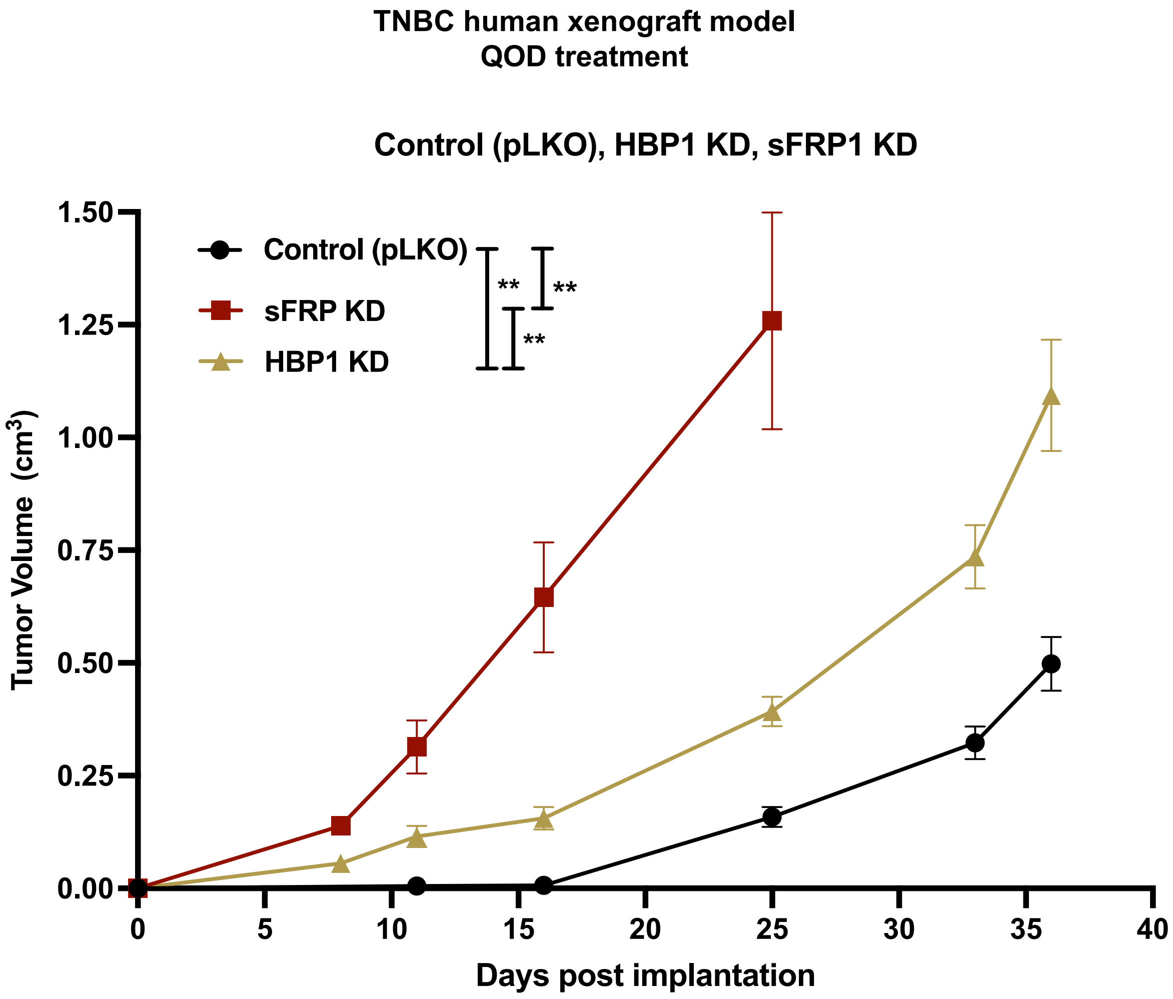
TNBC human xenograft model
QOD treatment
Control (pLKO), HBP1 KD, sFRP1 KD
Control (pLKO)
sFRP KD
HBP1 KD
**
**
**
Tumor Volume (cm3)
1.50
1.25
1.00
0.75
0.50
0.25
0.00
0
5
10
15
20
25
30
35
40
Days post implantation

C

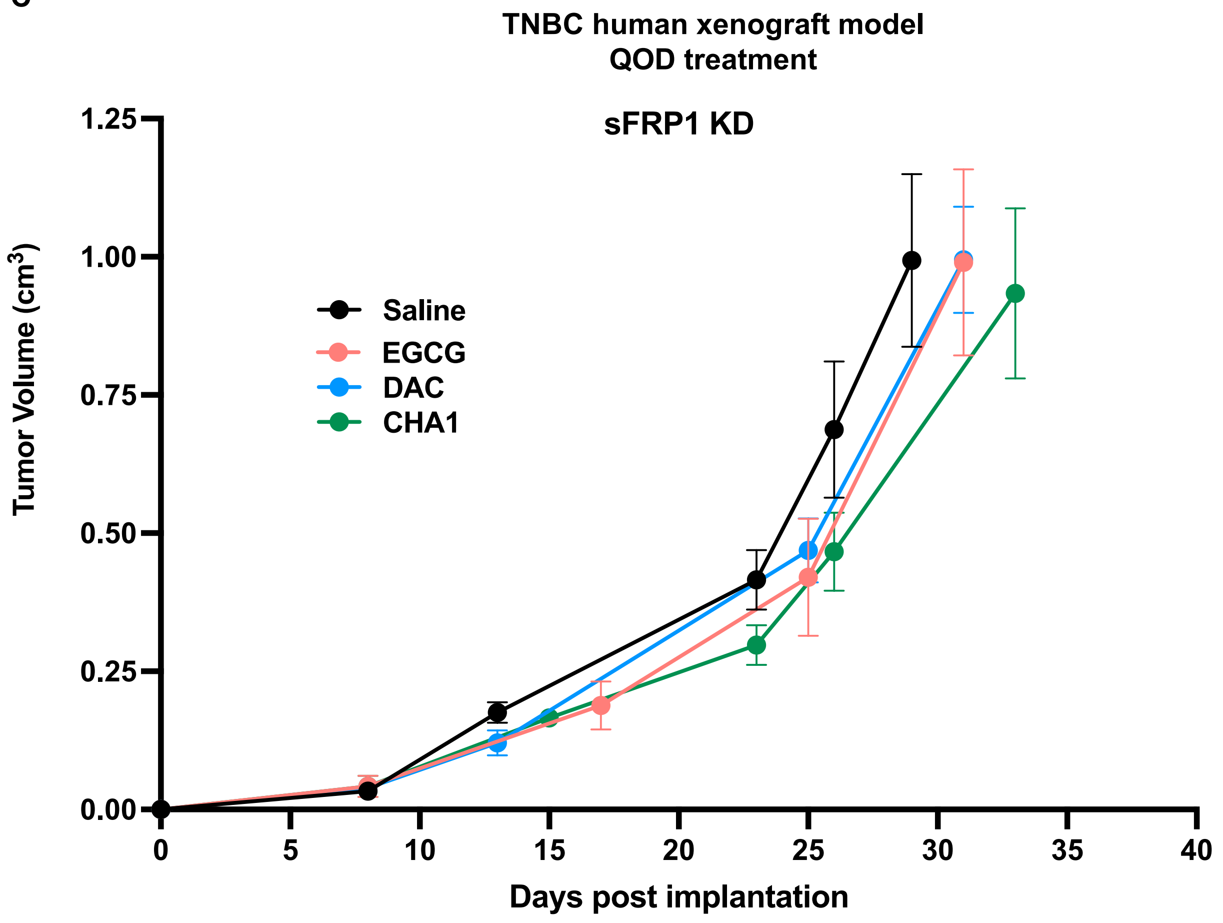
TNBC human xenograft model
QOD treatment
sFRP1 KD
Tumor Volume (cm3)
1.25
1.00
0.75
0.50
0.25
0.00
Saline
EGCG
DAC
CHA1
0
5
10
15
20
25
30
35
40
Days post implantation

D

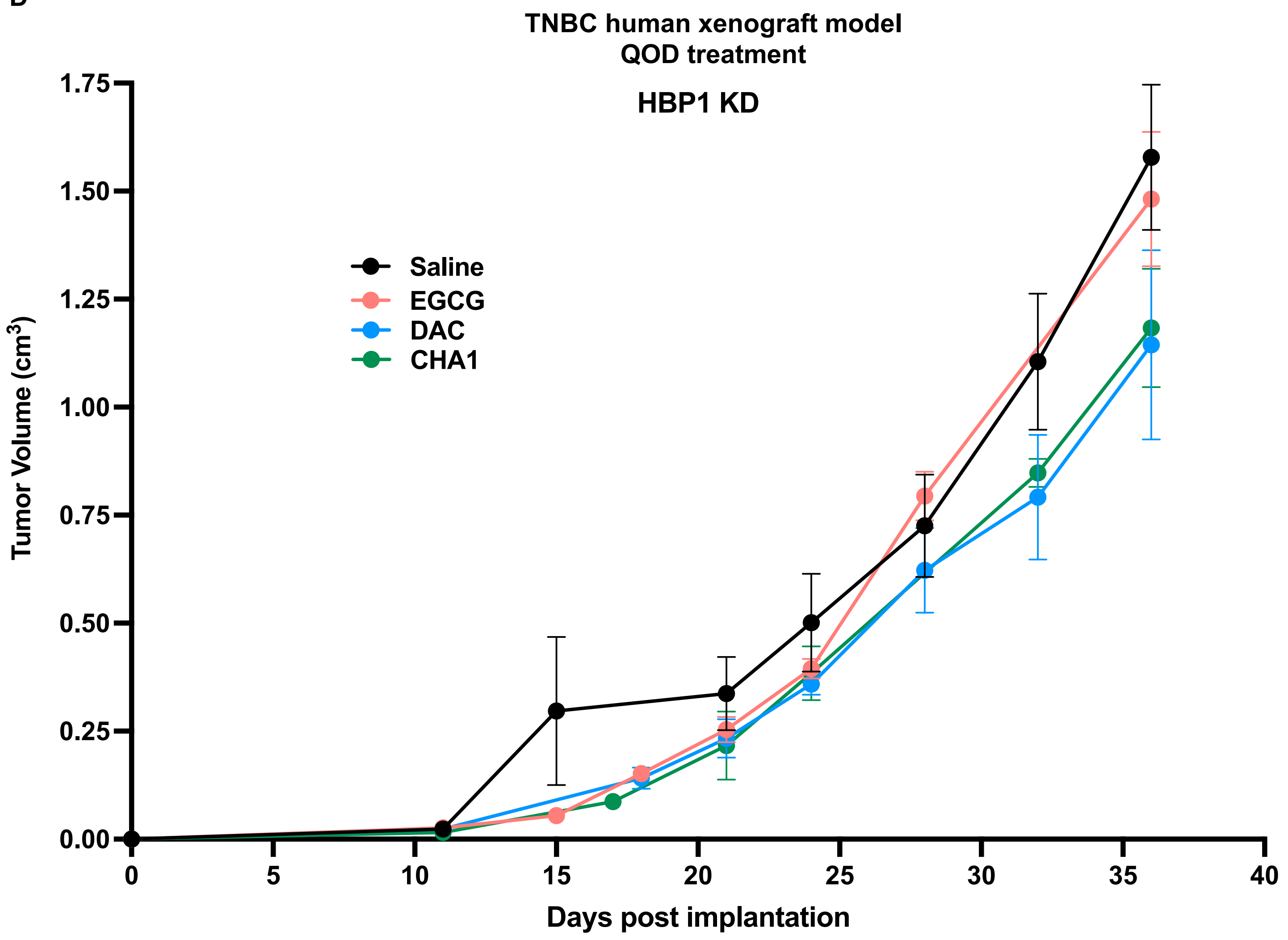

TNBC human xenograft model
QOD treatment
HBP1 KD
Saline
EGCG
DAC
CHA1
Tumor Volume (cm3)
1.75
1.50
1.25
1.00
0.75
0.50
0.25
0.00
0
5
10
15
20
25
30
35
40
Days post implantation

E

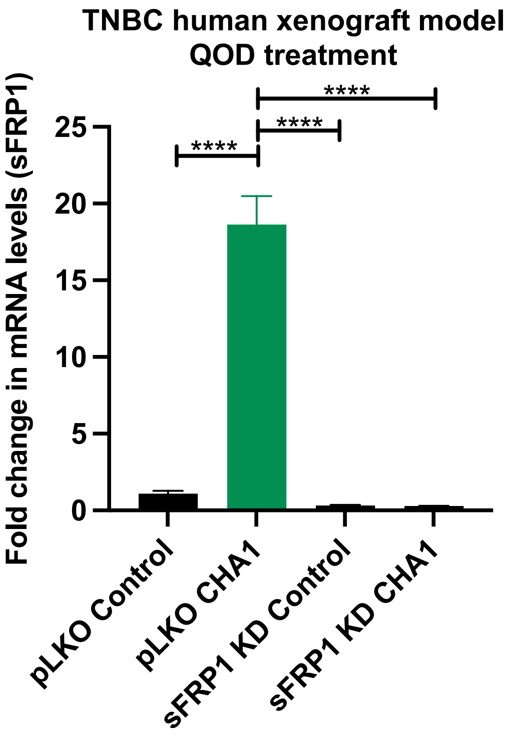
TNBC human xenograft model
QOD treatment
Fold change in mRNA levels (sFRP1)
25
20
15
10
5
0
****
****
****
pLKO Control
pLKO CHA1
sFRP1 KD Control
sFRP1 KD CHA1

F

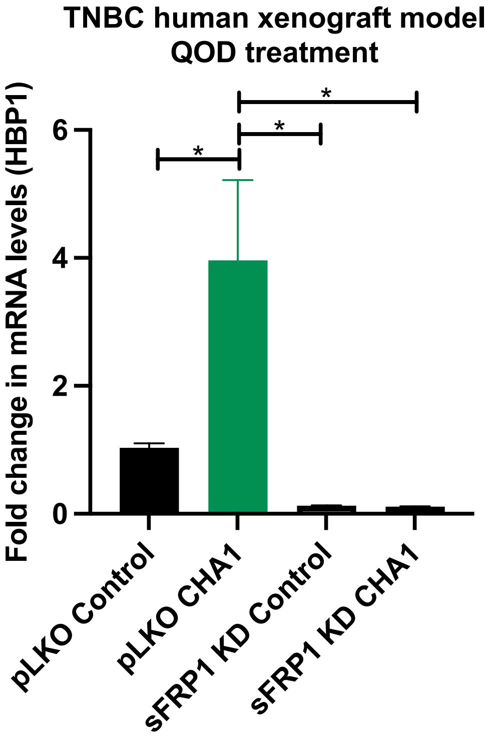
TNBC human xenograft model
QOD treatment
Fold change in mRNA levels (HBP1)
6
4
2
0
*
*
*
pLKO Control
pLKO CHA1
sFRP1 KD Control
sFRP1 KD CHA1

G

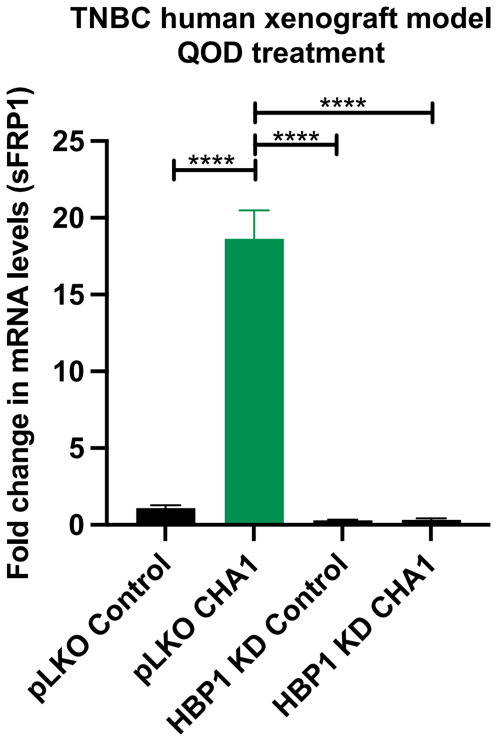
TNBC human xenograft model
QOD treatment
Fold change in mRNA levels (sFRP1)
25
20
15
10
5
0
****
****
****
pLKO Control
pLKO CHA1
HBP1 KD Control
HBP1 KD CHA1

H

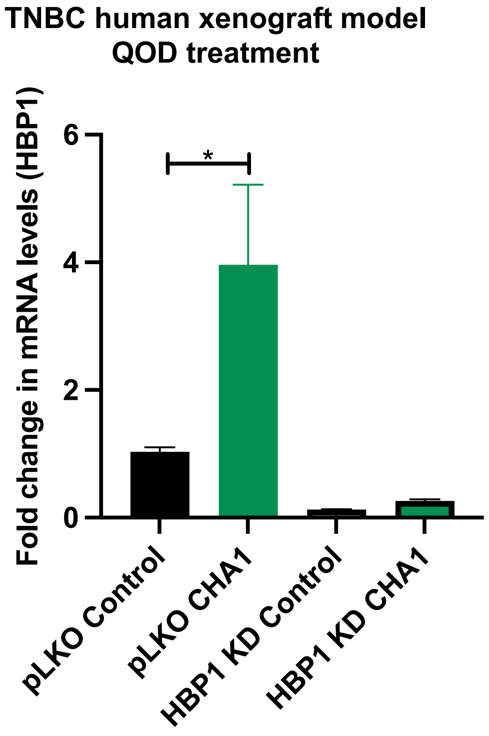
TNBC human xenograft model
QOD treatment
Fold change in mRNA levels (HBP1)
6
4
2
0
*
pLKO Control
pLKO CHA1
HBP1 KD Control
HBP1 KD CHA1

I

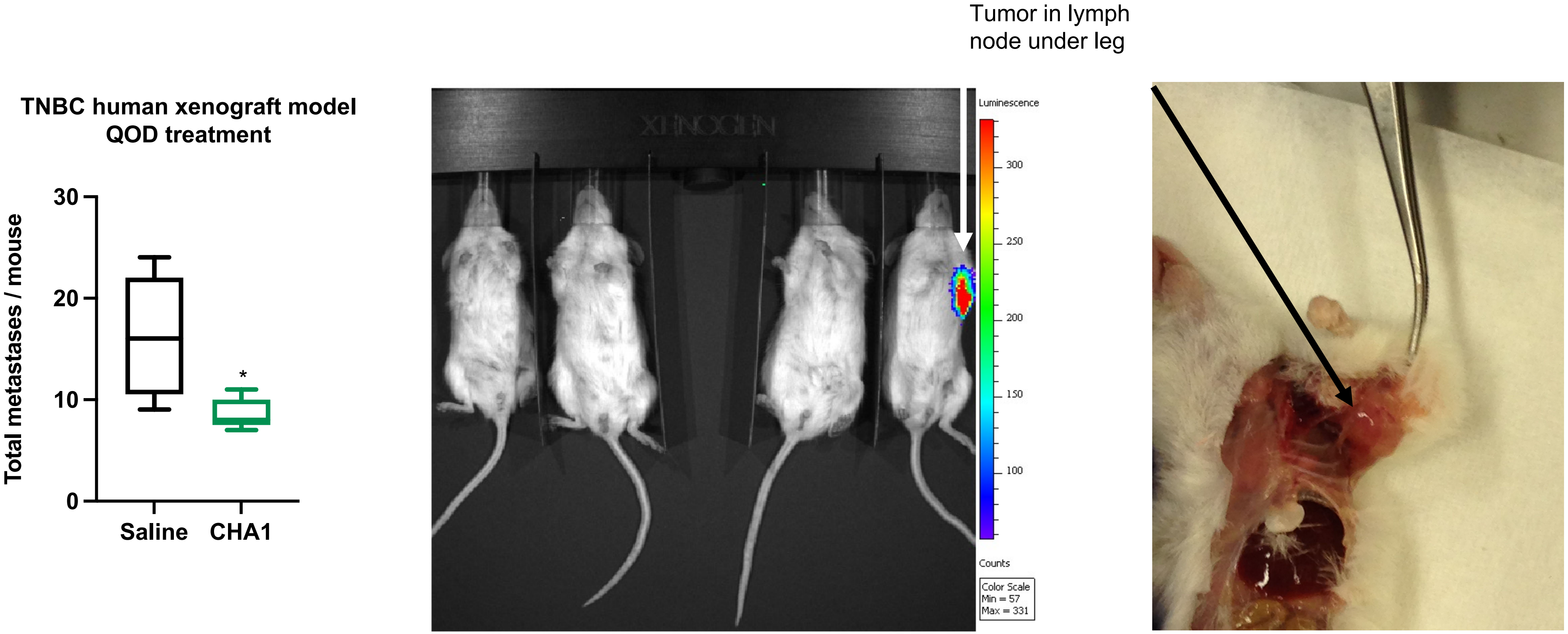
TNBC human xenograft model
QOD treatment
Total metastases / mouse
30
20
10
0
*
Saline
CHA1
Tumor in lymph
node under leg
Luminescence
300
250
200
150
100
Counts
Color Scale
Min = 57
Max = 331

J

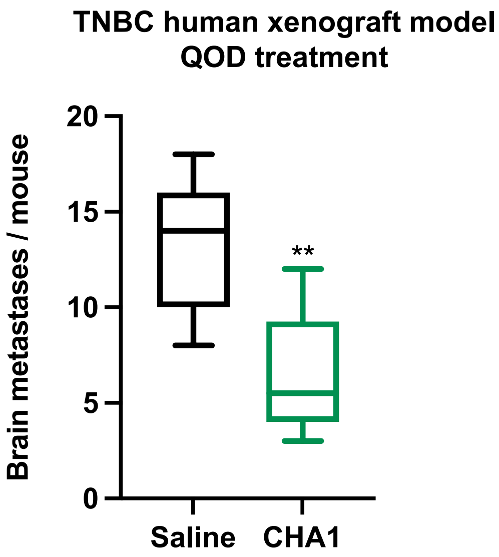


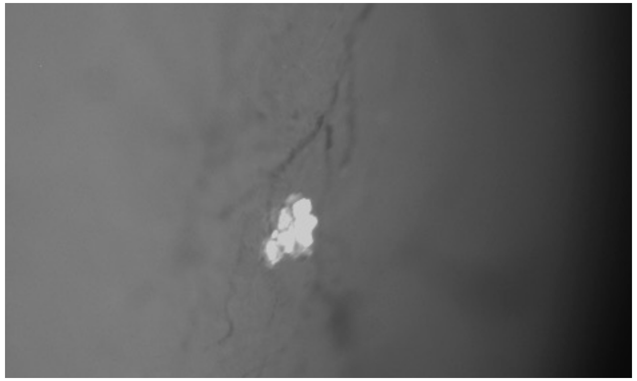

GFP imaging of a small brain metastasis comprising of a cluster of ~ 20 tumor-derived cells within the cortex of a control saline-treated mouse.

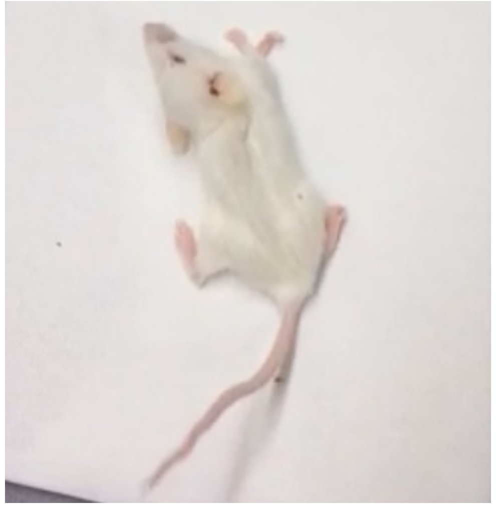

Control mouse observed seizing ~12 weeks post tumor resection.

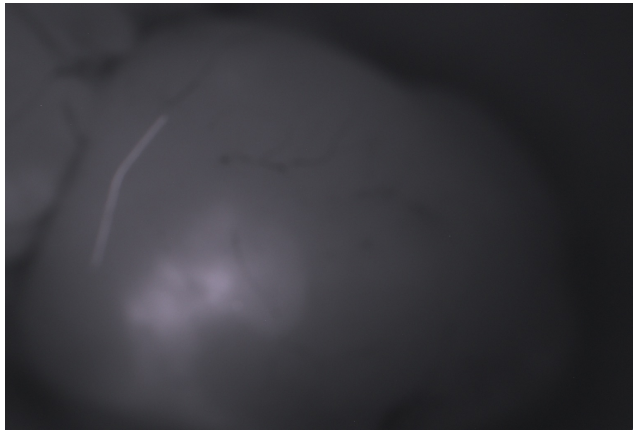

GFP imaging of a large metastasis in the left cortex from the seizing control mouse to the left.

K

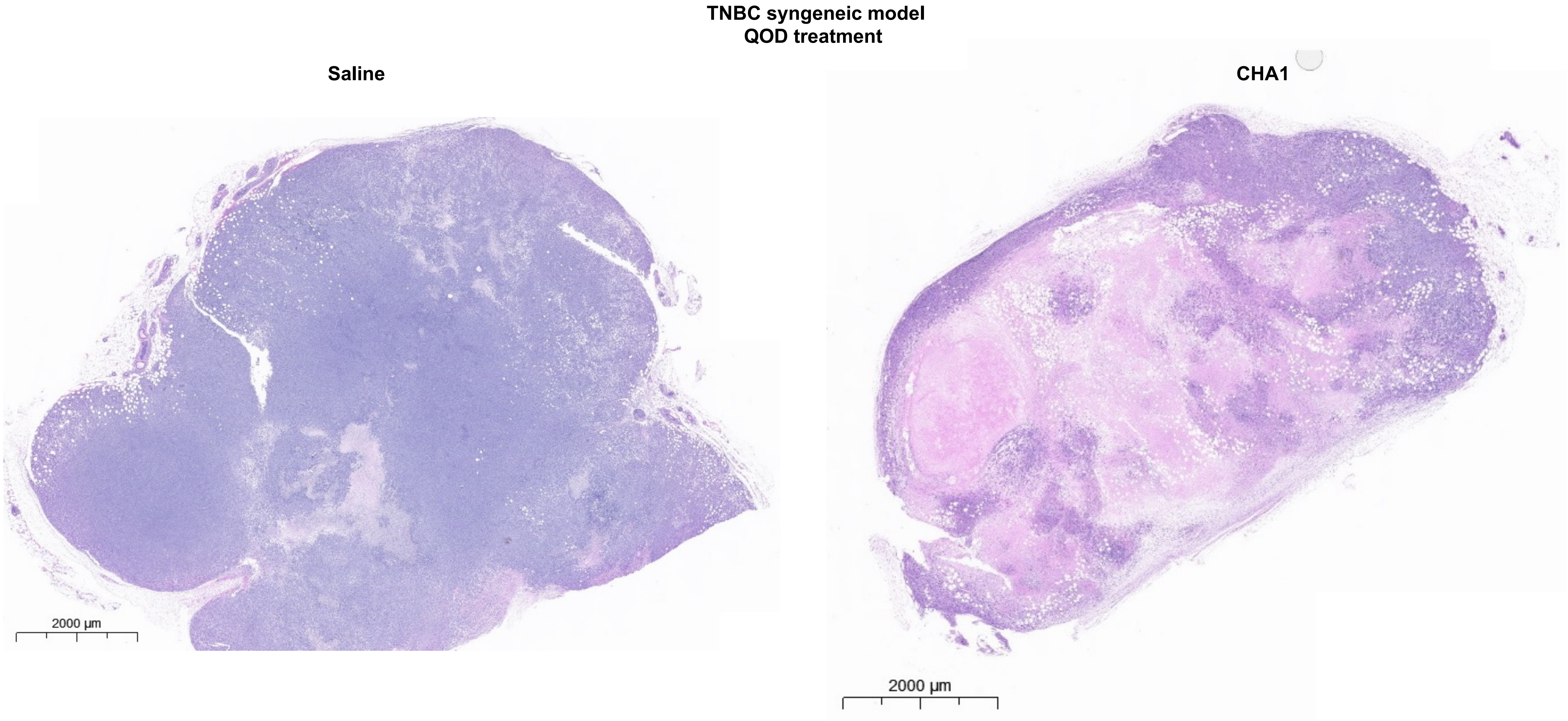
TNBC syngeneic model
QOD treatment
Saline
CHA1
2000 µm
2000 µm

L

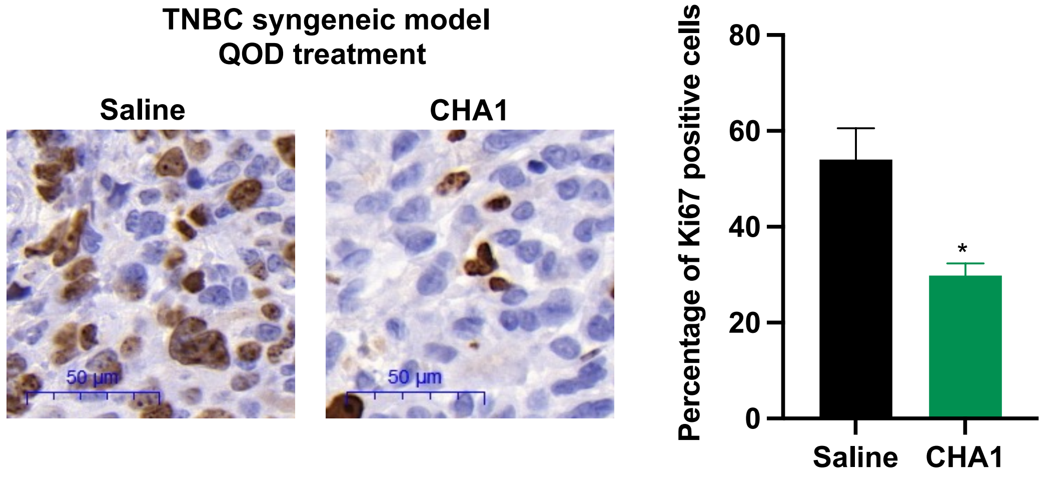
TNBC syngeneic model
QOD treatment
Saline
CHA1
50 µm
50 µm
Percentage of Ki67 positive cells
80
60
40
20
0
*
Saline
CHA1

M

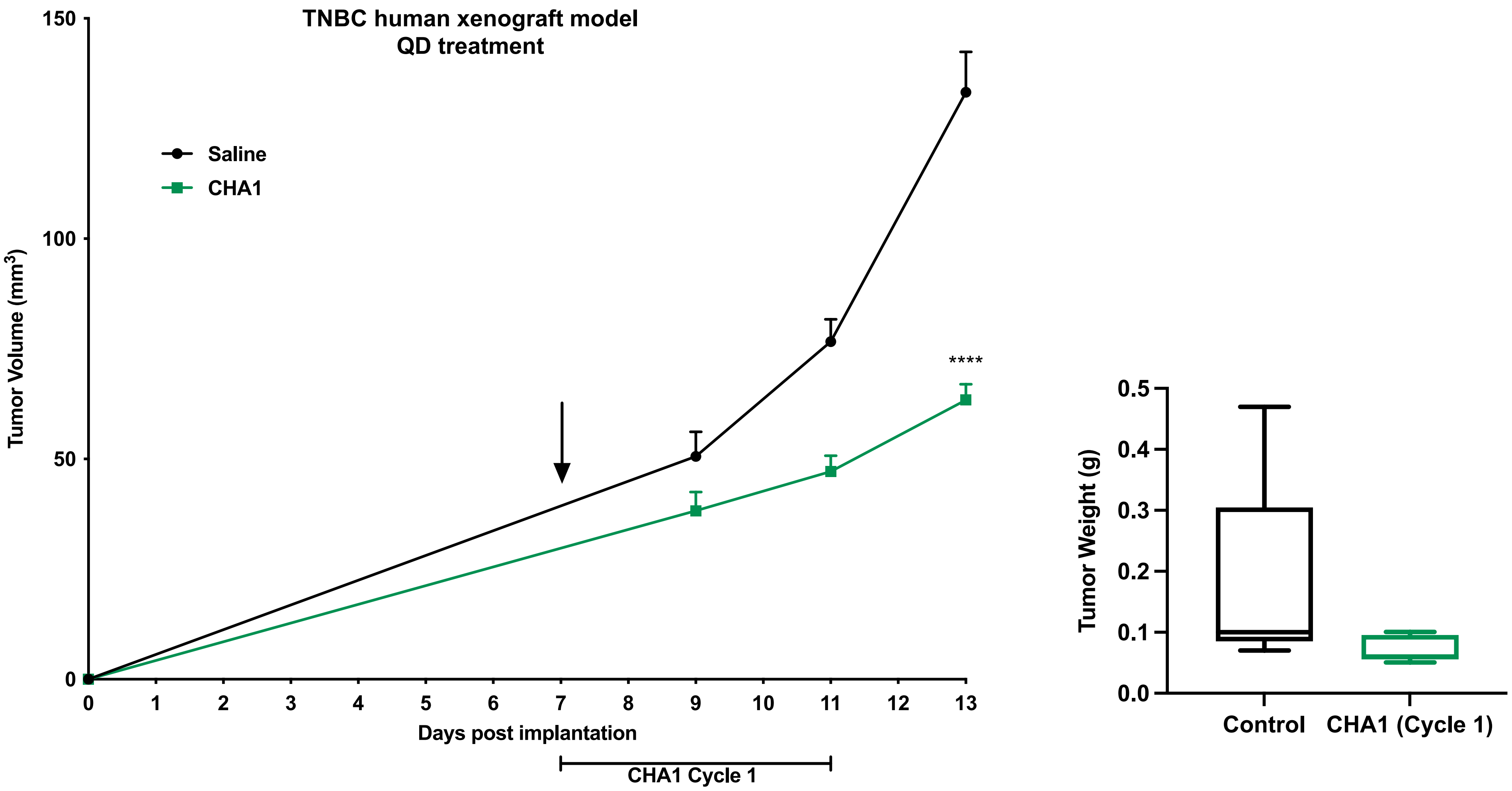

TNBC human xenograft model
QD treatment
Saline
CHA1
Tumor Volume (mm3)
150
100
50
0
****
0
1
2
3
4
5
6
7
8
9
10
11
12
13
Days post implantation
CHA1 Cycle 1
Tumor Weight (g)
0.5
0.4
0.3
0.2
0.1
0.0
Control
CHA1 (Cycle 1)

A

TNBC human xenograft
QOD treatment

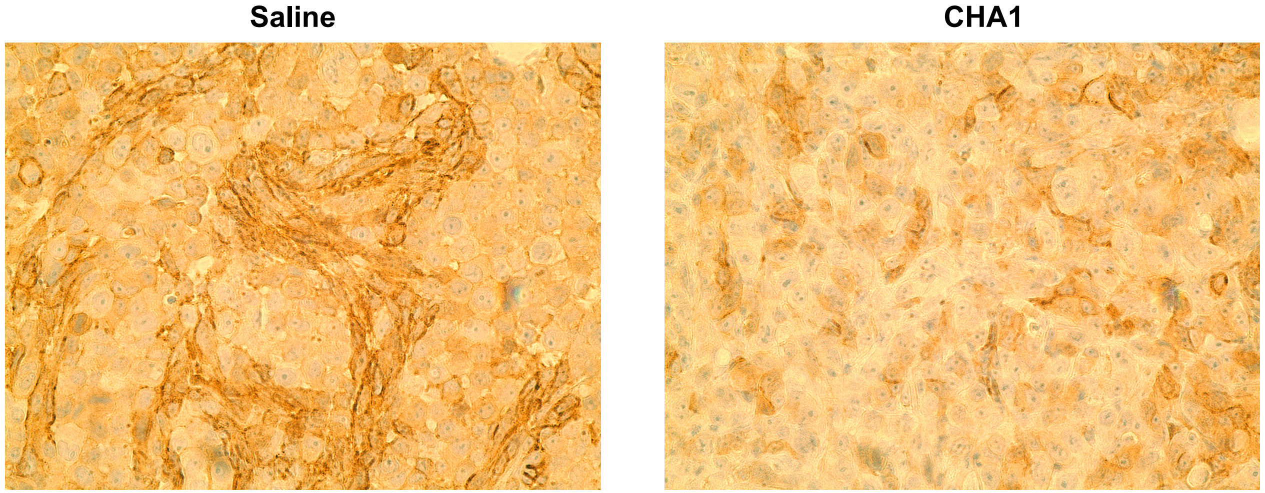

B

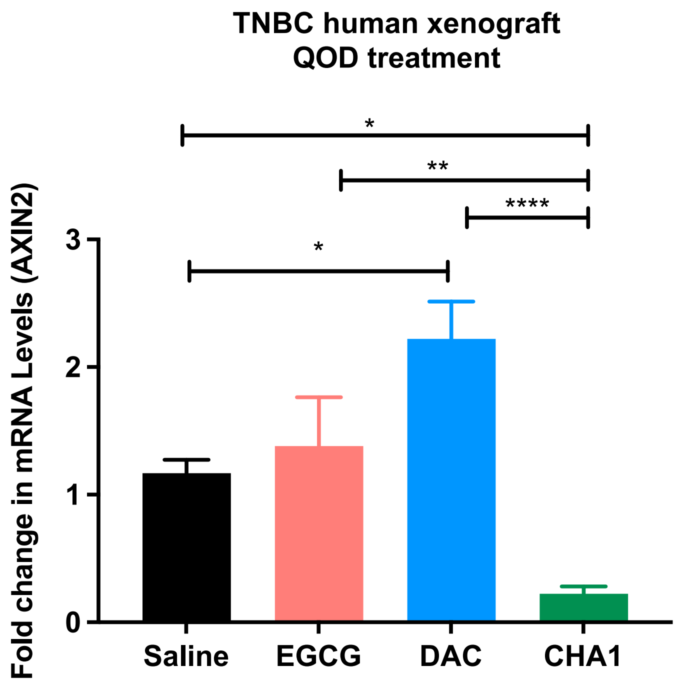
TNBC human xenograft
QOD treatment
*
**
****
*
Fold change in mRNA Levels (AXIN2)
3
2
1
0
Saline
EGCG
DAC
CHA1

C

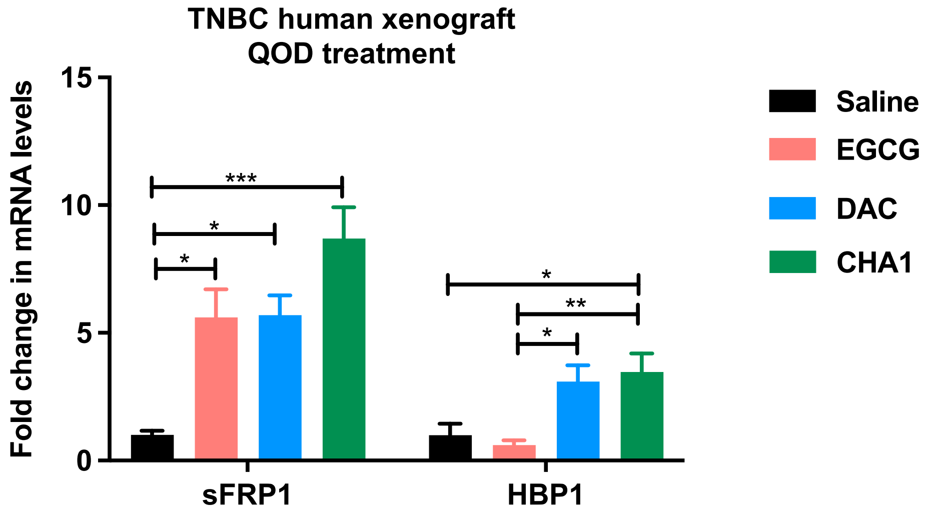
TNBC human xenograft
QOD treatment
Fold change in mRNA levels
15
10
5
0
***
*
*
*
**
*
sFRP1
HBP1
Saline
EGCG
DAC
CHA1

A

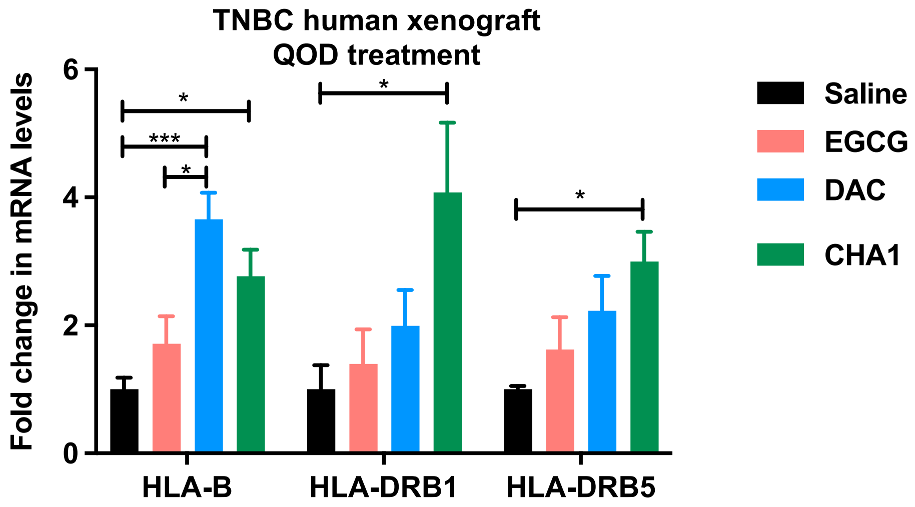
TNBC human xenograft
QOD treatment
Fold change in mRNA levels
0
2
4
6
*
***
*
*
*
HLA-B
HLA-DRB1
HLA-DRB5
Saline
EGCG
DAC
CHA1

**B**

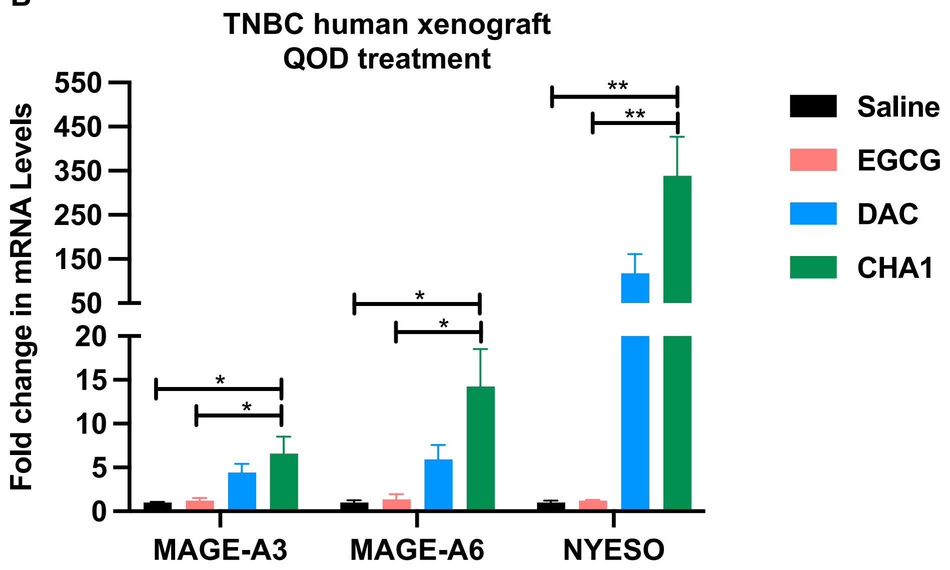
TNBC human xenograft
QOD treatment
Fold change in mRNA Levels
550
450
350
250
150
50
20
15
10
5
0
*
*
*
*
**
**
MAGE-A3
MAGE-A6
NYESO
Saline
EGCG
DAC
CHA1

C

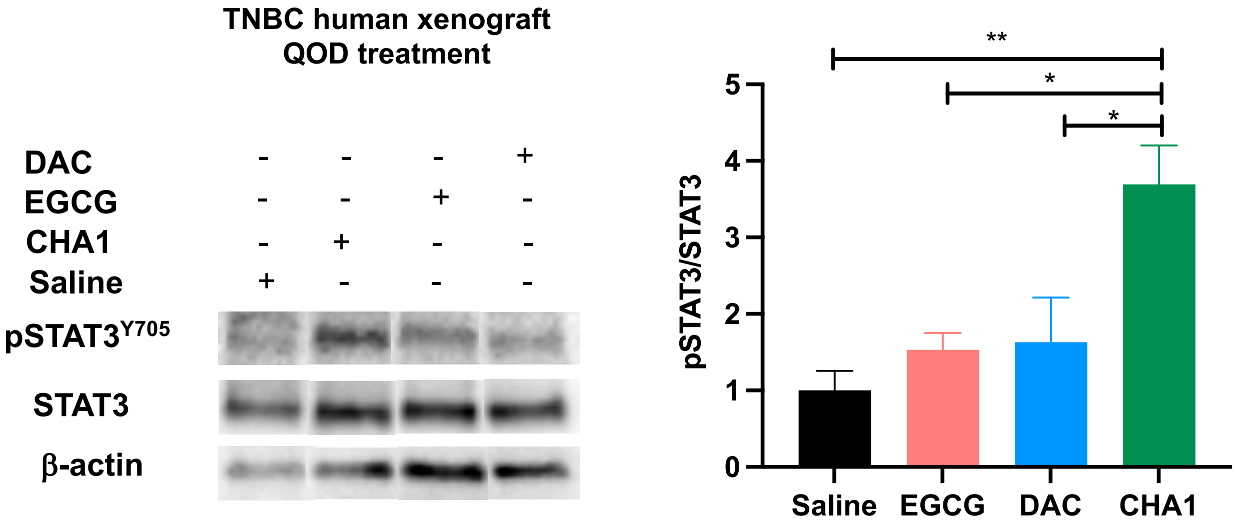
TNBC human xenograft
QOD treatment
DAC - - - +
EGCG - - + -
CHA1 - + - -
Saline + - - -
pSTAT3Y705
STAT3
β-actin
pSTAT3/STAT3
0
1
2
3
4
5
**
*
*
Saline
EGCG
DAC
CHA1

D

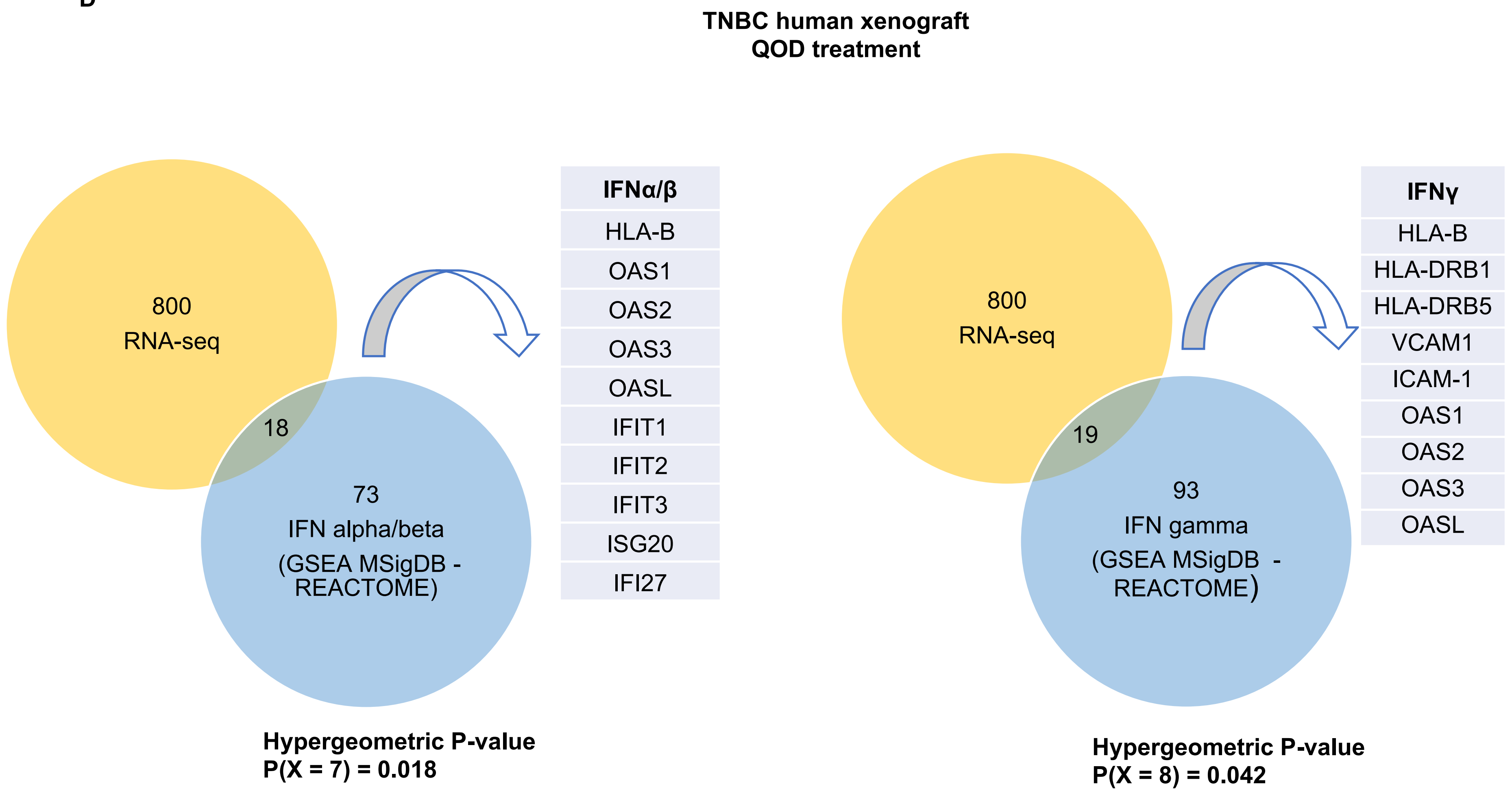
TNBC human xenograft
QOD treatment
800
RNA-seq
18
73
IFN alpha/beta
(GSEA MSigDB -
REACTOME)
IFNα/β
HLA-B
OAS1
OAS2
OAS3
OASL
IFIT1
IFIT2
IFIT3
ISG20
IFI27
Hypergeometric P-value
P(X = 7) = 0.018
800
RNA-seq
19
93
IFN gamma
(GSEA MSigDB -
REACTOME)
IFNγ
HLA-B
HLA-DRB1
HLA-DRB5
VCAM1
ICAM-1
OAS1
OAS2
OAS3
OASL
Hypergeometric P-value
P(X = 8) = 0.042

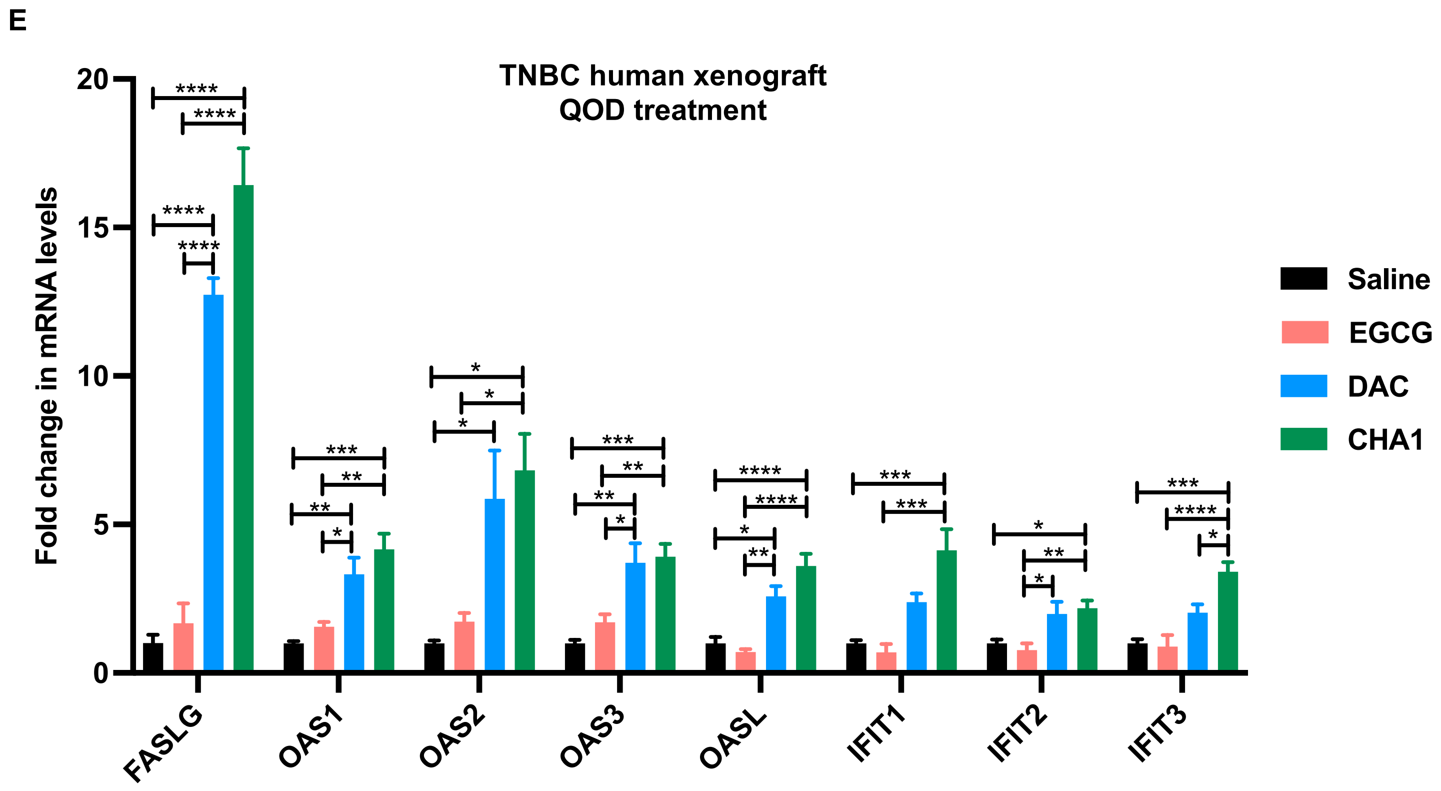
E
TNBC human xenograft
QOD treatment
Fold change in mRNA levels
20
15
10
5
0
FASLG
OAS1
OAS2
OAS3
OASL
IFIT1
IFIT2
IFIT3
Saline
EGCG
DAC
CHA1

F

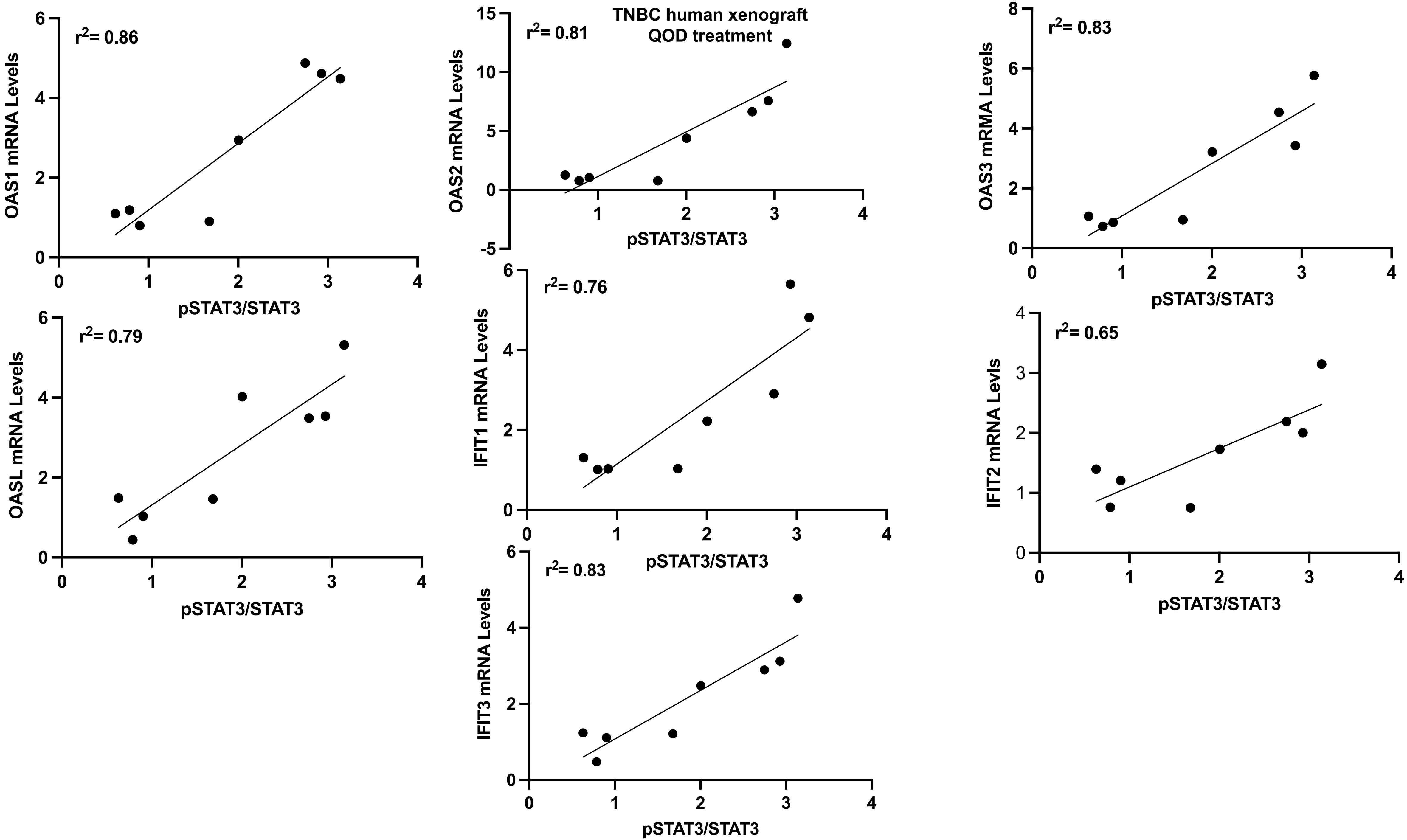

TNBC human xenograft
QOD treatment
$r^2$= 0.86
OAS1 mRNA Levels
pSTAT3/STAT3
$r^2$= 0.81
OAS2 mRNA Levels
pSTAT3/STAT3
$r^2$= 0.83
OAS3 mRMA Levels
pSTAT3/STAT3
$r^2$= 0.79
OASL mRNA Levels
pSTAT3/STAT3
$r^2$= 0.76
IFIT1 mRNA Levels
pSTAT3/STAT3
$r^2$= 0.65
IFIT2 mRNA Levls
pSTAT3/STAT3
$r^2$= 0.83
IFIT3 mRNA Levels
pSTAT3/STAT3

A

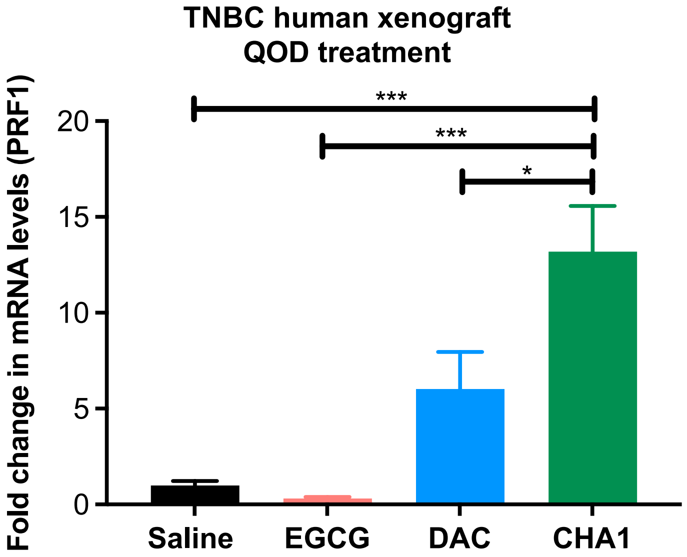
TNBC human xenograft
QOD treatment
Fold change in mRNA levels (PRF1)
20
15
10
5
0
***
***
*
Saline
EGCG
DAC
CHA1

B

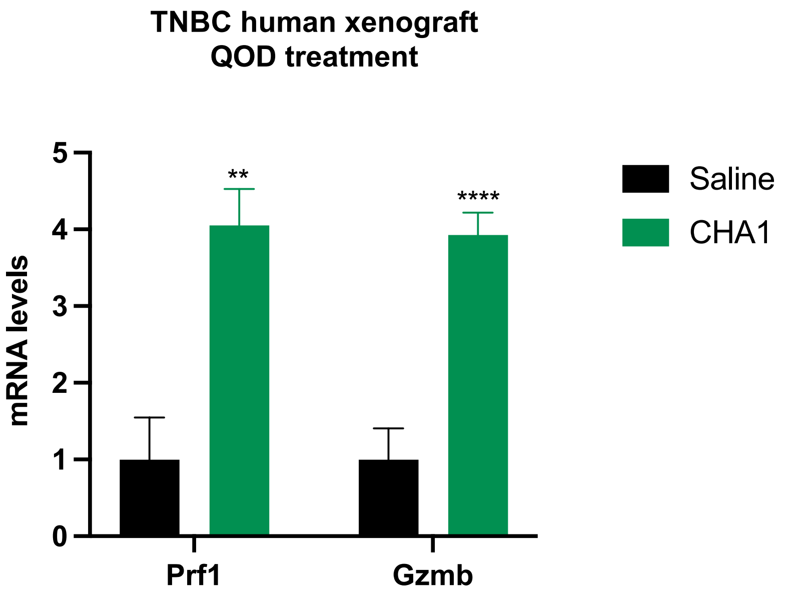
TNBC human xenograft
QOD treatment
mRNA levels
5
4
3
2
1
0
**
****
Prf1
Gzmb
Saline
CHA1

C

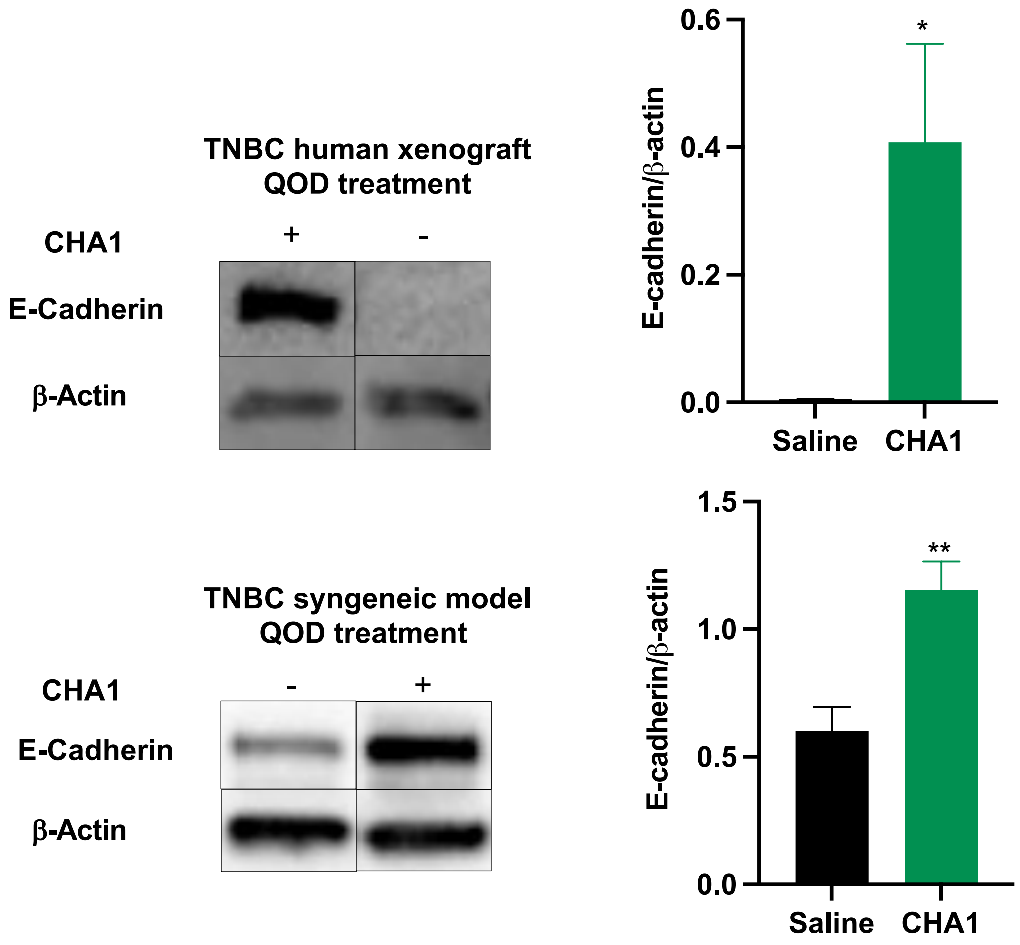
TNBC human xenograft
QOD treatment
CHA1
+
-
E-Cadherin
β-Actin
E-cadherin/β-actin
0.6
0.4
0.2
0.0
*
Saline
CHA1
TNBC syngeneic model
QOD treatment
CHA1
-
+
E-Cadherin
β-Actin
E-cadherin/β-actin
1.5
1.0
0.5
0.0
**
Saline
CHA1

D

TNBC human xenograft
QOD treatment

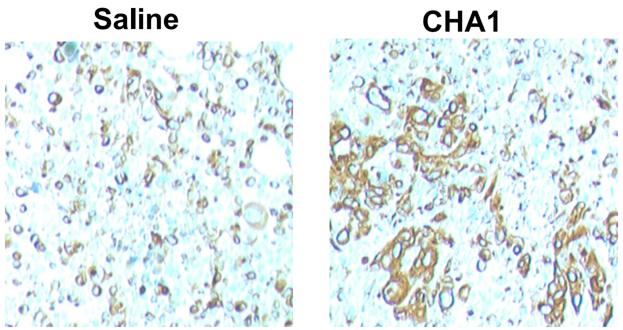

E

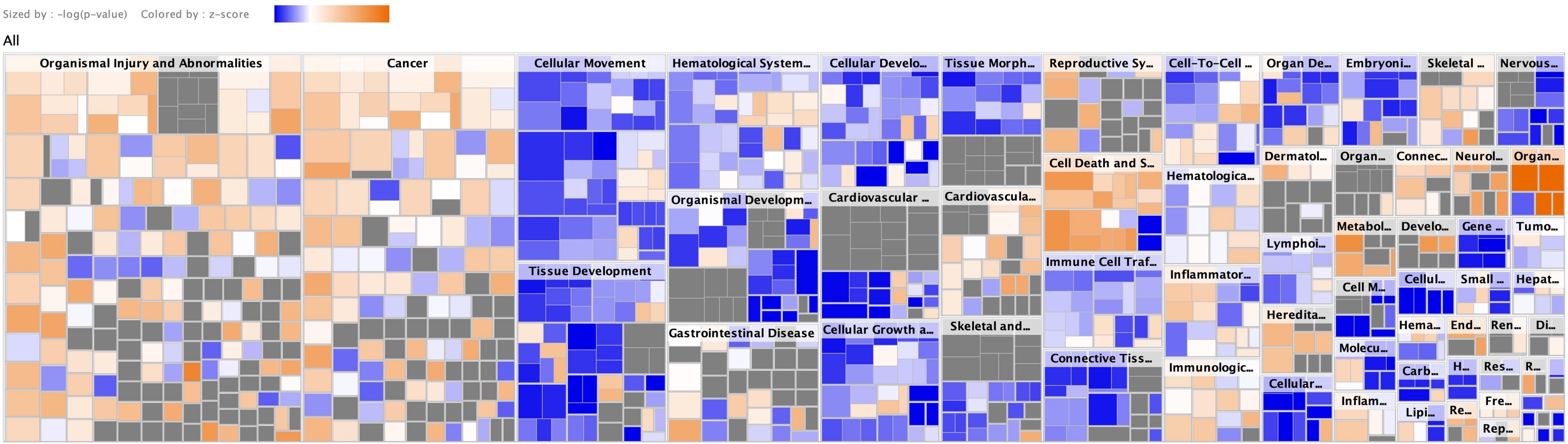


© 2000-2021 QIAGEN. All rights reserved.

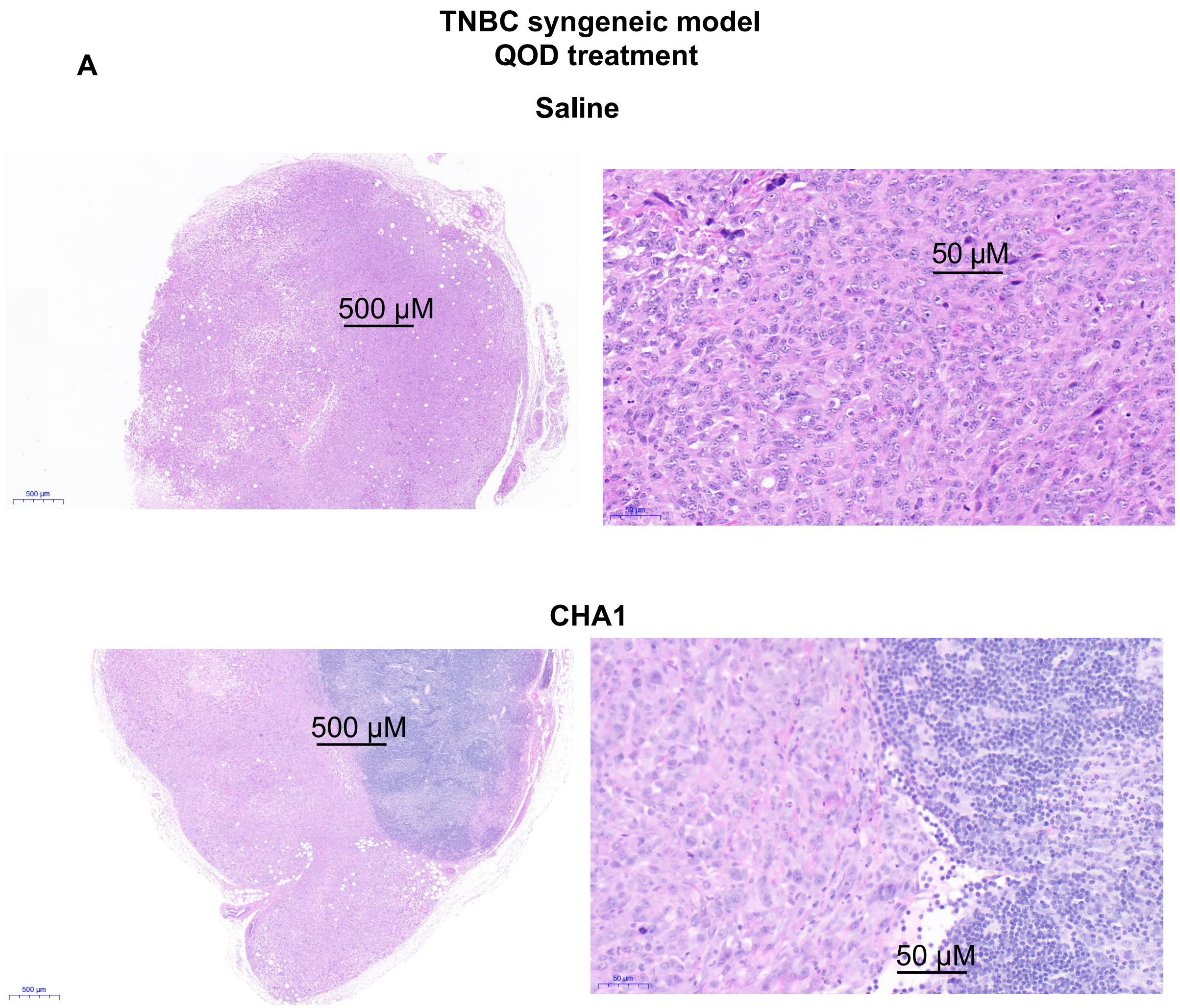
A
TNBC syngeneic model
QOD treatment
Saline
500 μM
50 μM
CHA1
500 μM
50 μM

**B**

**TNBC syngeneic model**
**CHA1 QOD treatment**
**$CD8^+$ T-cell staining**

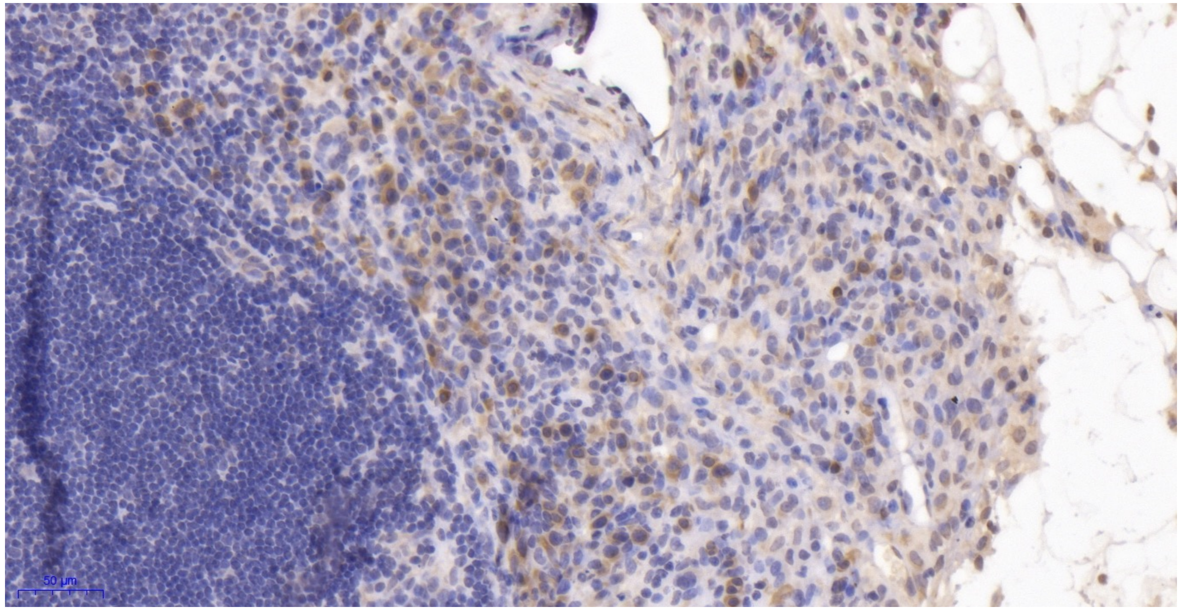


**C**

**TNBC syngeneic model**
**CHA1 QOD treatment**
**$F480^+$ macrophage staining**

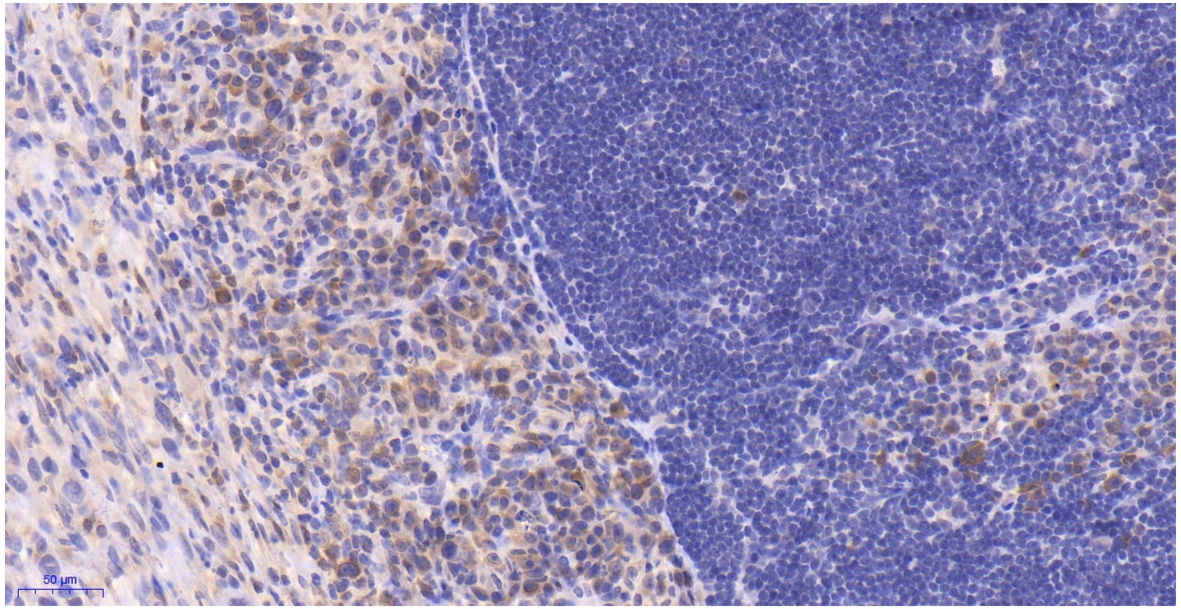

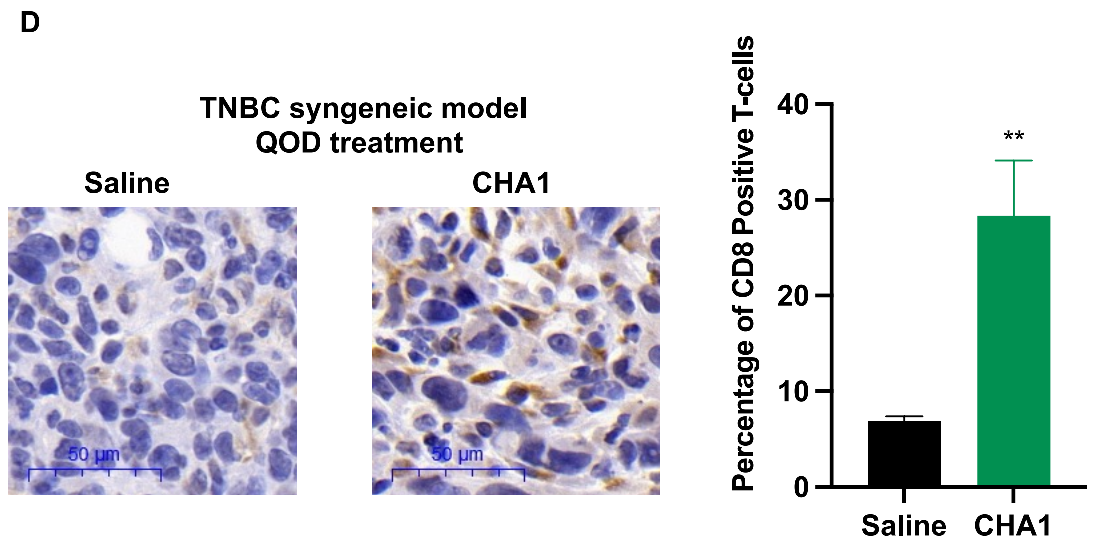
D
TNBC syngeneic model
QOD treatment
Saline
CHA1
50 µm
50 µm
Percentage of CD8 Positive T-cells
40
30
20
10
0
**
Saline
CHA1

A

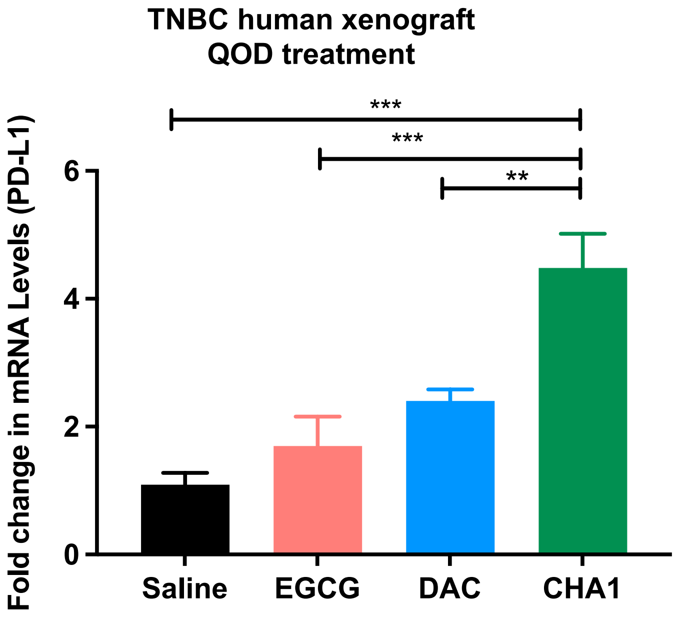
TNBC human xenograft
QOD treatment
***
***
**
Fold change in mRNA Levels (PD-L1)
6
4
2
0
Saline
EGCG
DAC
CHA1

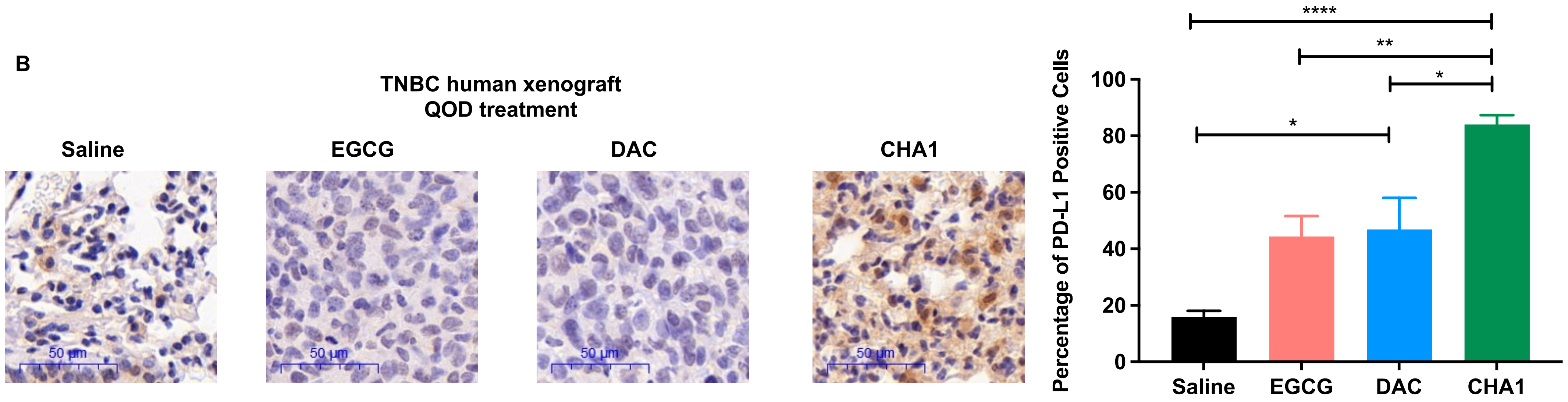
B
TNBC human xenograft
QOD treatment
Saline
EGCG
DAC
CHA1
50 µm
50 µm
50 µm
50 µm
Percentage of PD-L1 Positive Cells
0
20
40
60
80
100
****
**
*
*
Saline
EGCG
DAC
CHA1

C

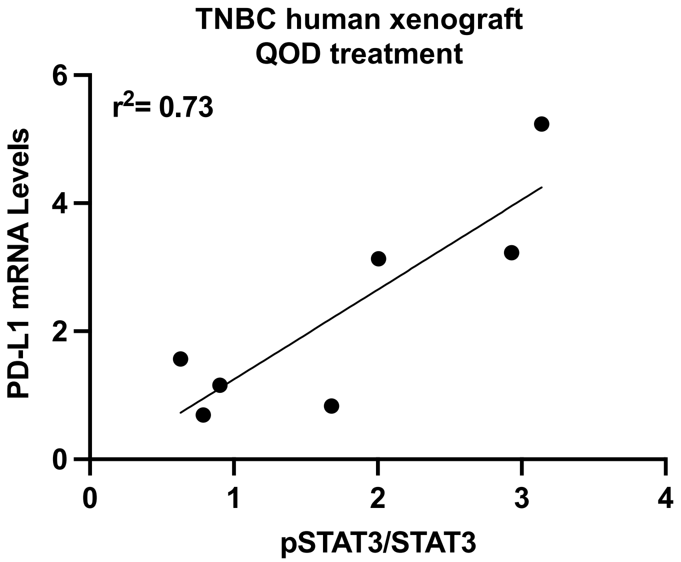
TNBC human xenograft
QOD treatment
r2= 0.73
PD-L1 mRNA Levels
pSTAT3/STAT3
0
2
4
6
0
1
2
3
4

**D**

| Published T-cell inflamed signature |
|---|
| CCL2 |
| CCL3 |
| CCL4 |
| CCL5 |
| CD27 |
| CD276 |
| CD4 |
| CD8A |
| CMKLR1 |
| CTLA4 |
| CXCL10 |
| CXCL9 |
| CXCR6 |
| EOMES |
| FOXP3 |
| GZMB |
| GZMK |
| HLA-DMA |
| HLA-DMB |
| HLA-DOA |
| HLA-DOB |
| HLA-DQA1 |
| HLA-DRB1 |
| HLA-E |
| ICOS |
| IDO1 |
| IFNG |
| IRF1 |
| LAG3 |
| NKG7 |
| PD-L1 |
| PD-L2 |
| PRF1 |
| PSMB10 |
| STAT1 |
| TIGIT |
| TNF |

| Overlap genes |
|---|
| Ccl4 |
| Ccl5 |
| Cd4 |
| Cd8a |
| Cxcl10 |
| Cxcl9 |
| Gzmb |
| HLA-DRB1 |
| Icos |
| Ido1 |
| Ifng |
| Lag3 |
| PDL-1, Pdl-1 |
| Prf1 |
| Psmb10 |
| Stat1 |
| Tigit |

**hypergeometric p-value = 7.7e-46**

| CHA1 gene signature |
|---|
| Ccl4 |
| Ccl5 |
| Cd4 |
| CD74 |
| Cd8a |
| Ctla-4 |
| Cxcl10 |
| Cxcl9 |
| Gzmb |
| HAL-DRB1 |
| HLA-B |
| HLA-DRB5 |
| ICAM-1 |
| Icos |
| Ido1 |
| IFI27, Ifi27 |
| IFIT1, Ifit1 |
| IFIT2 |
| IFIT3, Ifit3 |
| Ifng |
| ISG20 |
| Lag3 |
| OAS1, Oas1a, Oas1b, Oas1g |
| OAS2 |
| OAS3, Oas3 |
| OASL, Oasl1, Oasl2 |
| Pd-1 |
| PD-L1, Pd-l1 |
| Prf1 |
| Psmb10 |
| PSMB9 |
| Stat1 |
| Tigit |
| VCAM |

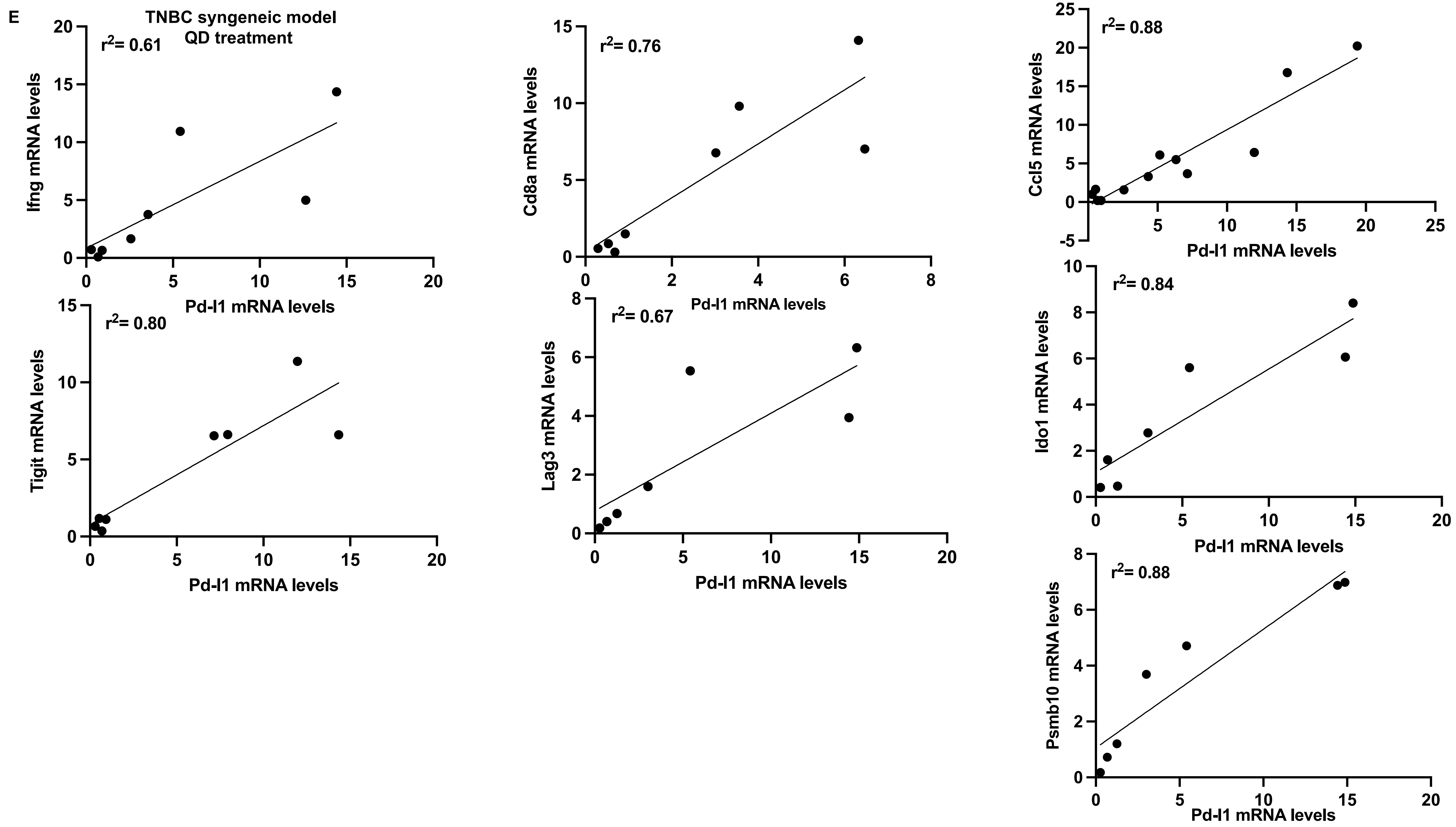
E
TNBC syngeneic model
QD treatment
r2= 0.61
Ifng mRNA levels
Pd-l1 mRNA levels
r2= 0.76
Cd8a mRNA levels
Pd-l1 mRNA levels
r2= 0.88
Ccl5 mRNA levels
Pd-l1 mRNA levels
r2= 0.80
Tigit mRNA levels
Pd-l1 mRNA levels
r2= 0.67
Lag3 mRNA levels
Pd-l1 mRNA levels
r2= 0.84
Ido1 mRNA levels
Pd-l1 mRNA levels
r2= 0.88
Psmb10 mRNA levels
Pd-l1 mRNA levels

F

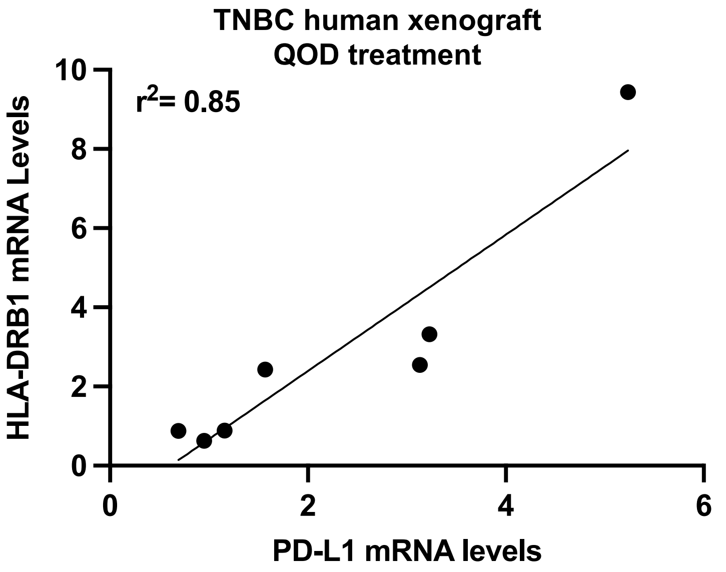

TNBC human xenograft
QOD treatment
r2= 0.85
HLA-DRB1 mRNA Levels
PD-L1 mRNA levels
0
2
4
6
8
10
0
2
4
6

G

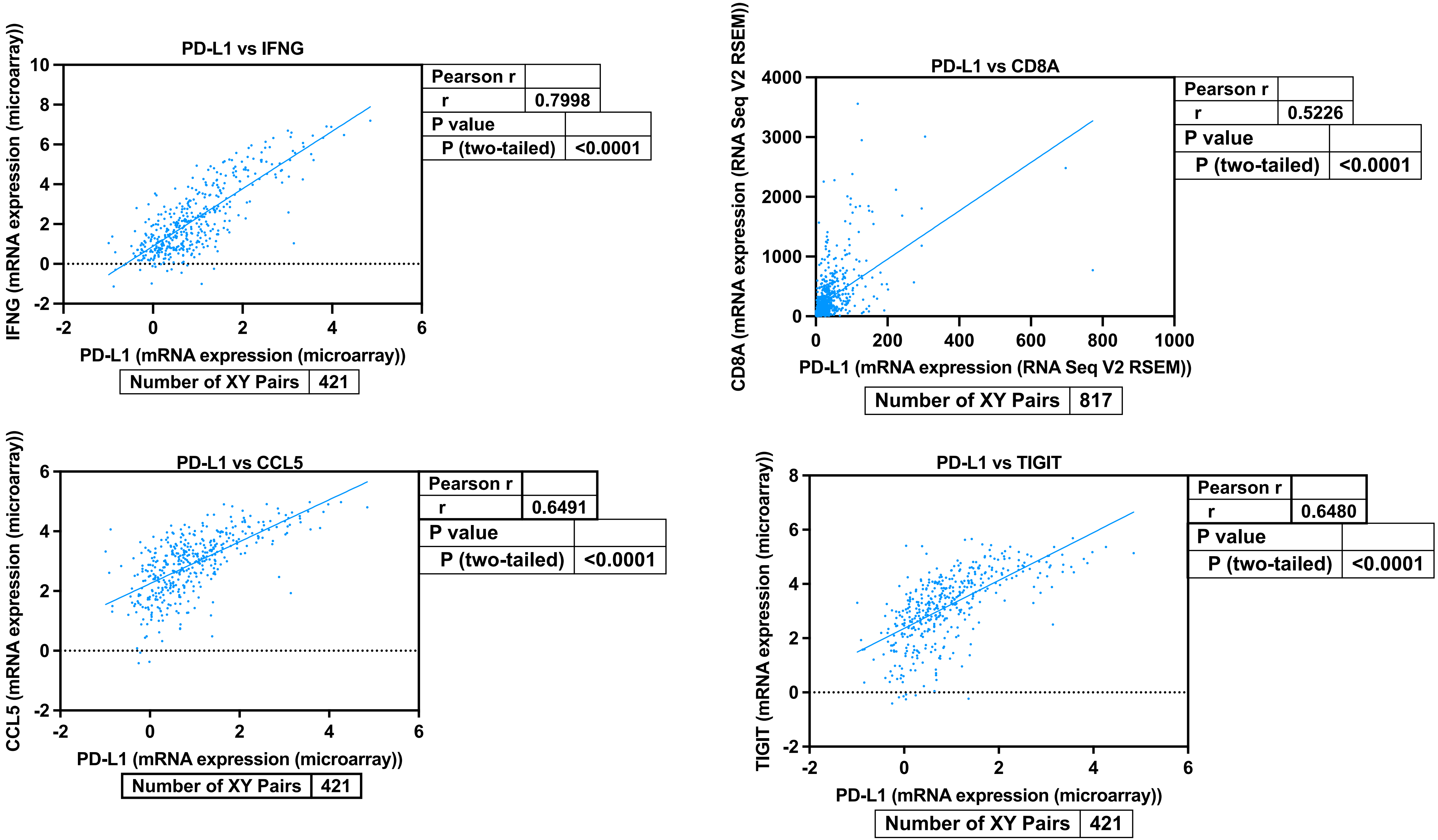

PD-L1 vs IFNG
IFNG (mRNA expression (microarray))
PD-L1 (mRNA expression (microarray))
Pearson r
r 0.7998
P value
P (two-tailed) <0.0001
Number of XY Pairs 421
PD-L1 vs CD8A
CD8A (mRNA expression (RNA Seq V2 RSEM))
PD-L1 (mRNA expression (RNA Seq V2 RSEM))
Pearson r
r 0.5226
P value
P (two-tailed) <0.0001
Number of XY Pairs 817
PD-L1 vs CCL5
CCL5 (mRNA expression (microarray))
PD-L1 (mRNA expression (microarray))
Pearson r
r 0.6491
P value
P (two-tailed) <0.0001
Number of XY Pairs 421
PD-L1 vs TIGIT
TIGIT (mRNA expression (microarray))
PD-L1 (mRNA expression (microarray))
Pearson r
r 0.6480
P value
P (two-tailed) <0.0001
Number of XY Pairs 421

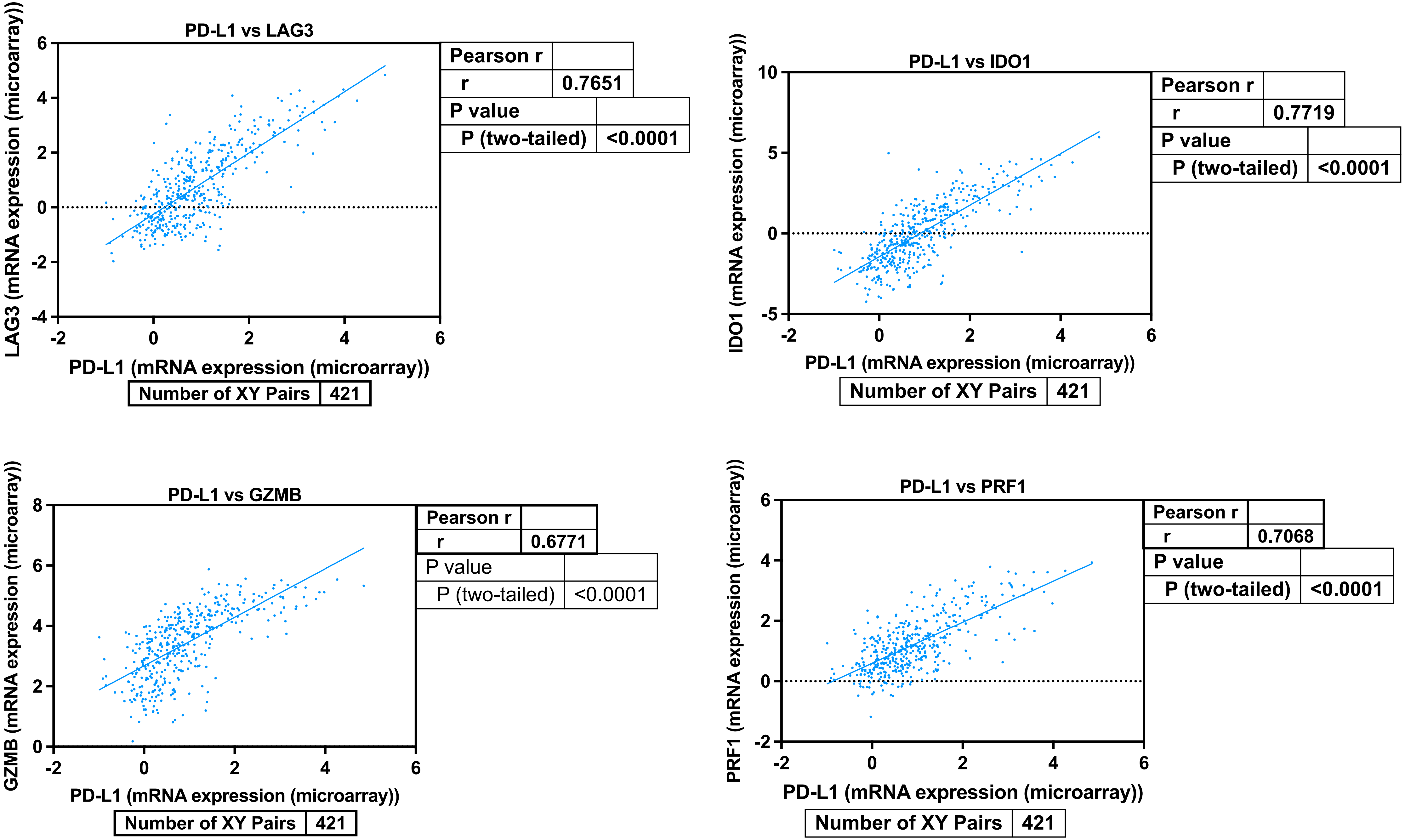
PD-L1 vs LAG3
LAG3 (mRNA expression (microarray))
PD-L1 (mRNA expression (microarray))
Pearson r
r 0.7651
P value
P (two-tailed) <0.0001
Number of XY Pairs 421
PD-L1 vs IDO1
IDO1 (mRNA expression (microarray))
PD-L1 (mRNA expression (microarray))
Pearson r
r 0.7719
P value
P (two-tailed) <0.0001
Number of XY Pairs 421
PD-L1 vs GZMB
GZMB (mRNA expression (microarray))
PD-L1 (mRNA expression (microarray))
Pearson r
r 0.6771
P value
P (two-tailed) <0.0001
Number of XY Pairs 421
PD-L1 vs PRF1
PRF1 (mRNA expression (microarray))
PD-L1 (mRNA expression (microarray))
Pearson r
r 0.7068
P value
P (two-tailed) <0.0001
Number of XY Pairs 421

A

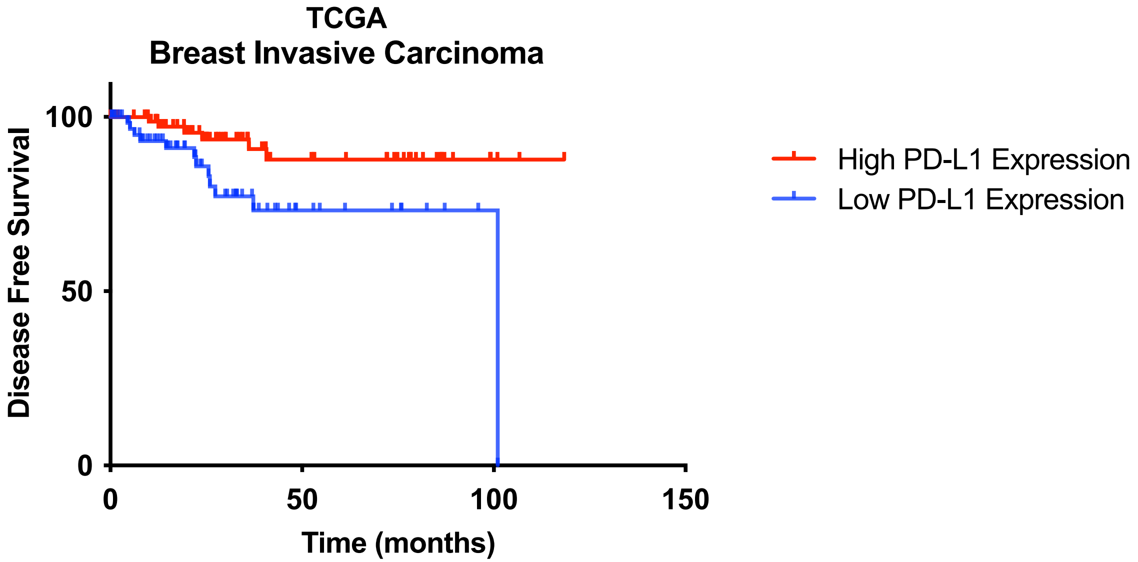


| | |
|---|---|
| P value | 0.0179 |

| | | |
|---|---|---|
| Hazard Ratio (logrank) | A/B | B/A |
| Ratio (and its reciprocal) | 0.3256 | 3.071 |
| 95% CI of ratio | 0.1265 to 0.8380 | 1.193 to 7.903 |

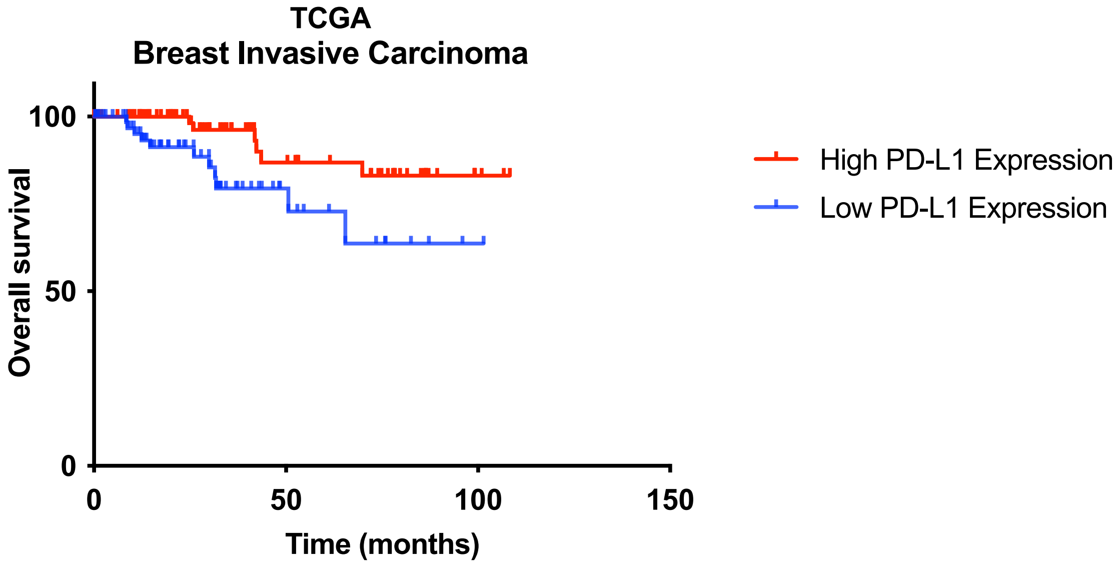


| | |
|---|---|
| P value | 0.0200 |

| | | |
|---|---|---|
| Hazard Ratio (logrank) | A/B | B/A |
| Ratio (and its reciprocal) | 0.3282 | 3.046 |
| 95% CI of ratio | 0.1230 to 0.8760 | 1.142 to 8.130 |

B

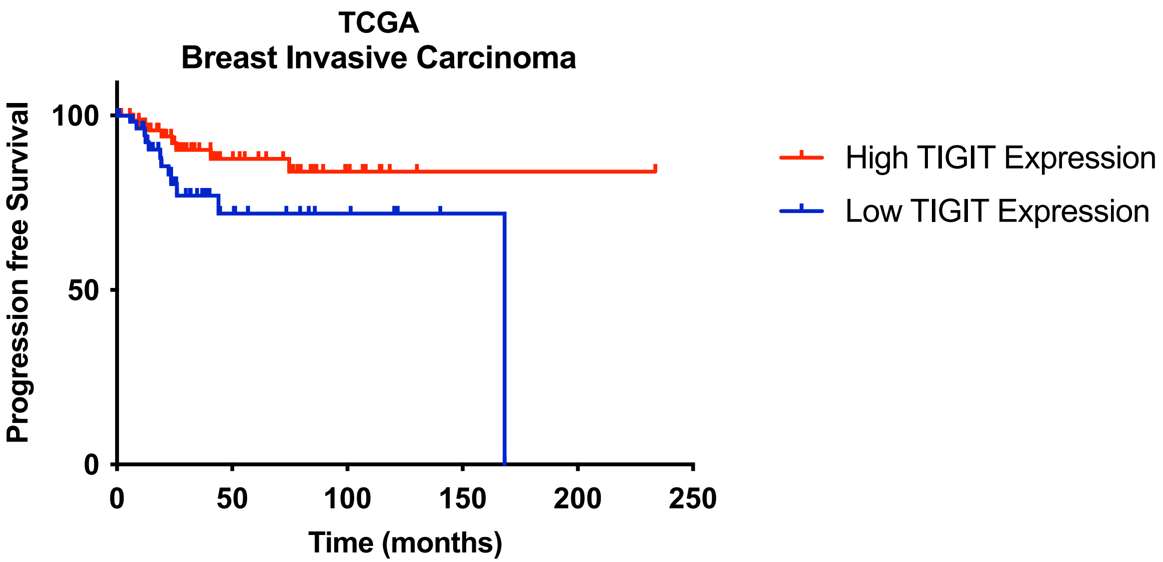


| | |
|---|---|
| P value | 0.0534 |

| | | |
|---|---|---|
| Hazard Ratio (logrank) | A/B | B/A |
| Ratio (and its reciprocal) | 0.4273 | 2.340 |
| 95% CI of ratio | 0.1740 to 1.049 | 0.9531 to 5.746 |

C

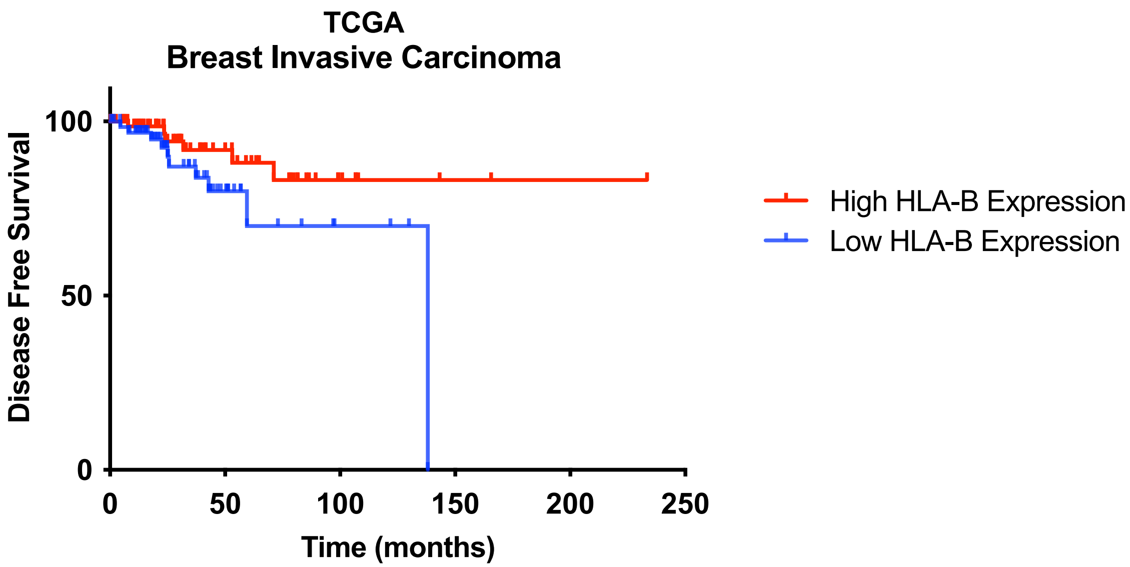


| | |
|---|---|
| P value | 0.0770 |

| | | |
|---|---|---|
| Hazard Ratio (logrank) | A/B | B/A |
| Ratio (and its reciprocal) | 0.4175 | 2.395 |
| 95% CI of ratio | 0.1542 to 1.130 | 0.8846 to 6.487 |

D

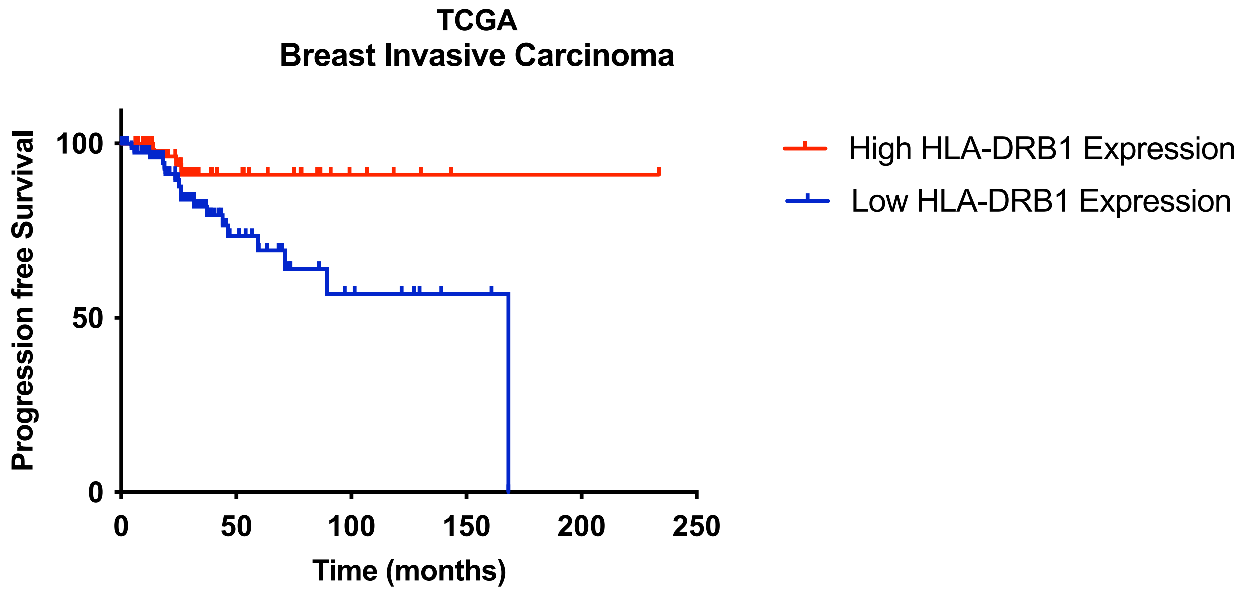


| | | |
|---|---|---|
| | P value | 0.0136 |
| Hazard Ratio (logrank) | A/B | B/A |
| Ratio (and its reciprocal) | 0.2799 | 3.572 |
| 95% CI of ratio | 0.1207 to 0.6493 | 1.540 to 8.286 |

E

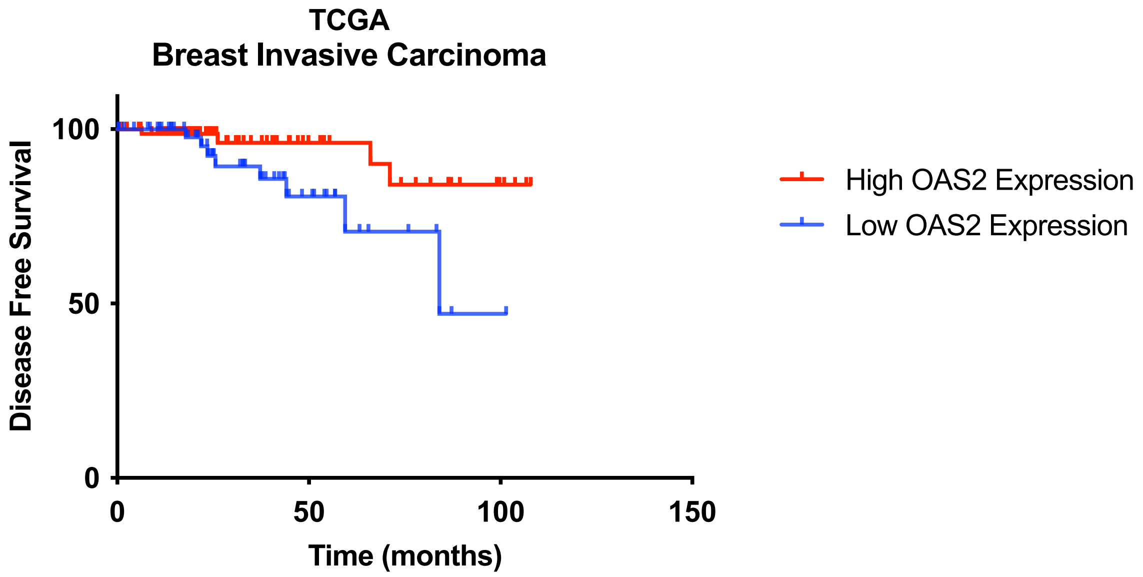


| | | |
|---|---|---|
| | P value | 0.0356 |
| Hazard Ratio (logrank) | A/B | B/A |
| Ratio (and its reciprocal) | 0.3033 | 3.297 |
| 95% CI of ratio | 0.09441 to 0.9745 | 1.026 to 10.59 |

F

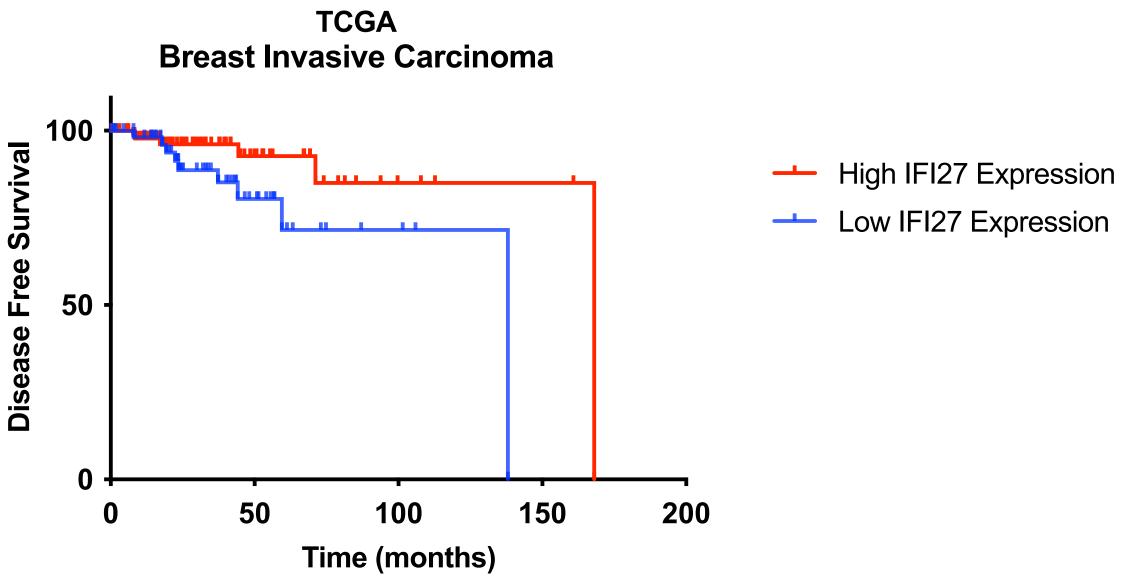


| | |
|---|---|
| P value | 0.0581 |

| | | |
|---|---|---|
| Hazard Ratio (logrank) | A/B | B/A |
| Ratio (and its reciprocal) | 0.3907 | 2.559 |
| 95% CI of ratio | 0.1369 to 1.115 | 0.8970 to 7.303 |

G

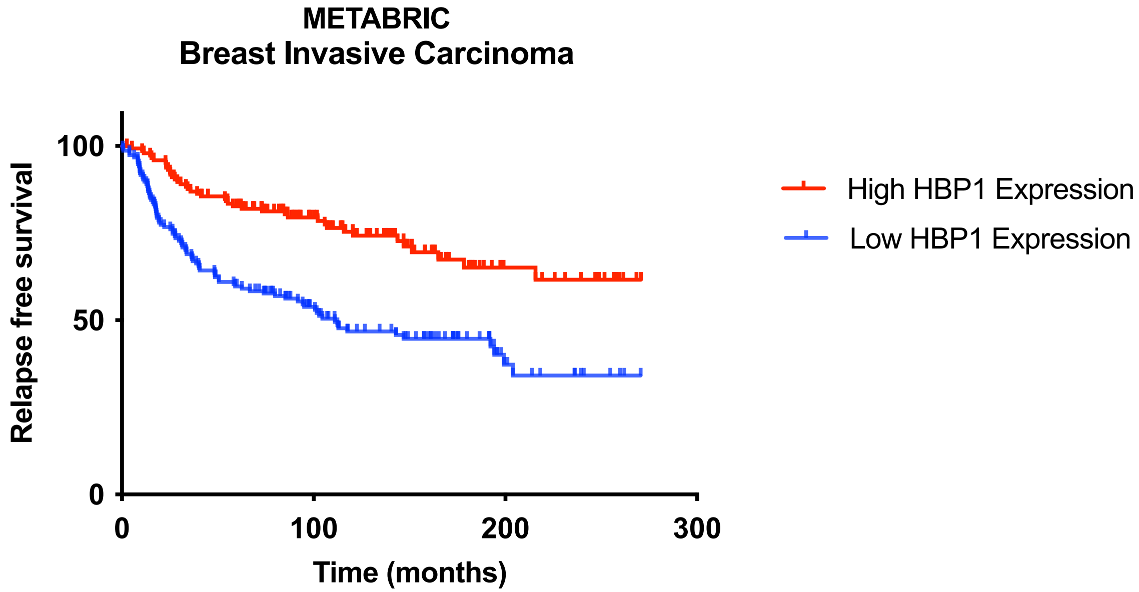


| | |
|---|---|
| P value | <0.0001 |

| | | |
|---|---|---|
| Hazard Ratio (logrank) | A/B | B/A |
| Ratio (and its reciprocal) | 0.4095 | 2.442 |
| 95% CI of ratio | 0.2877 to 0.5829 | 1.716 to 3.476 |

A

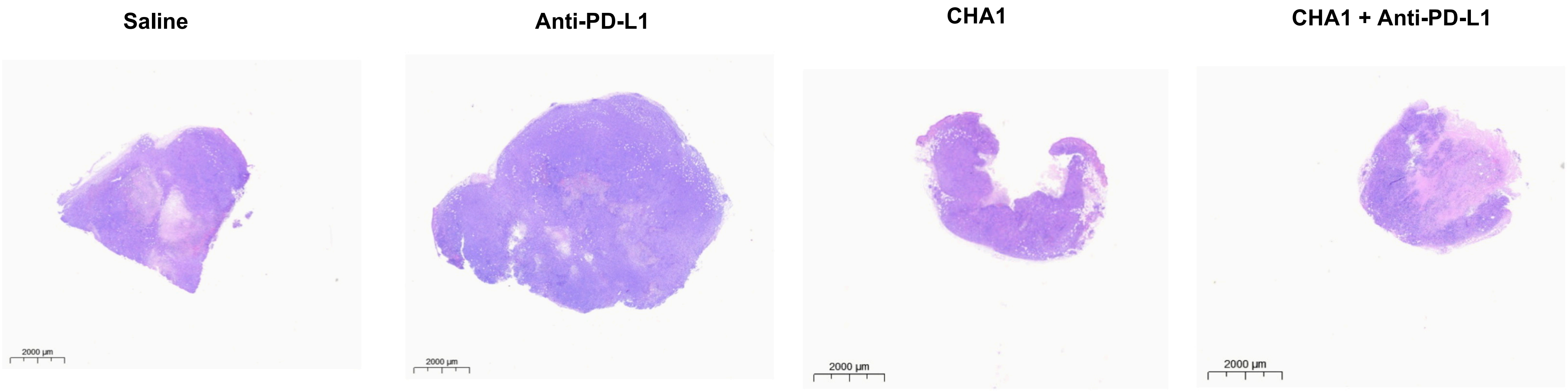
Saline
Anti-PD-L1
CHA1
CHA1 + Anti-PD-L1
2000 µm
2000 µm
2000 µm
2000 µm

B

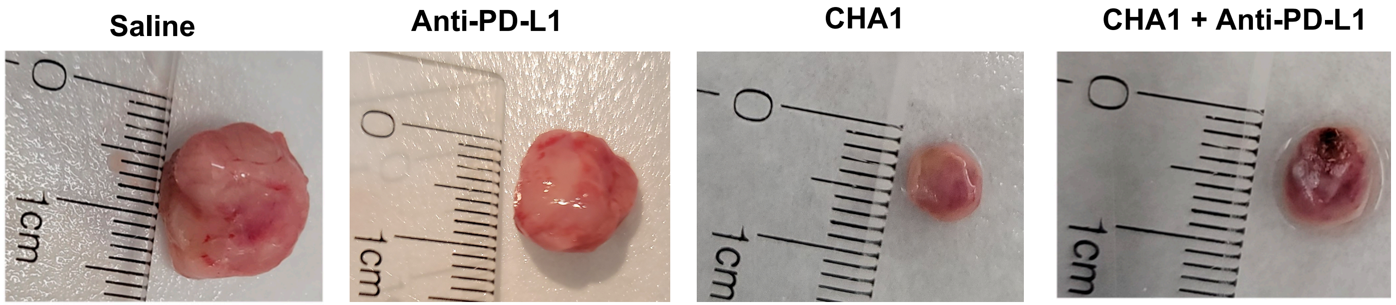

**Safety and efficacy considerations for combined use of EGCG and Decitabine**

A toxicological analysis in mice demonstrated tolerable dosing for 14 straight days i.p. at 21.1 mg/kg, while another report showed a dose of 50 and 75 mg/kg/i.p/day in mice , for 3 consecutive days or 3 times a week for eight weeks, were reported to have no elevated morbidity. The EGCG dose we used (16.5 mg/kg i.p.) was therefore well within safety limits. Further, the mouse dose was estimated to be somewhat less than an i.v.dose of 10 mg/kg, which reaches a peak plasma blood level (Cmax) of 0.28 ±0.08 umol/L. This plasma blood level can safely be achieved in humans orally with consumption of between 200 to 400 mg EGCG (1-4).

| Dose type | Dose mg/kg | Dose umol/kg | plasma uncong. (umol/L) | plasma total (umol/L) | Equivalent human dose |
|---|---|---|---|---|---|
| IV (lambert paper)* | 10 | 21.8 | 13.6±2 | 2.7 ±0.7 | 0.8 mg/kg (48 mg total) |
| IG (lambert paper)* | 75 | 163.8 | 0.04 ±0.01 | 0.28 ±0.08 | 6.0 mg/kg (360 mg total) |
| IV (toxicological report)** | | 235.6 | | 18.84 | |
| IP (toxicological report)** | 108 | 235.6 | | 4.56 | 8.64 mg/kg 518.4 mg total |
| PO (toxicological report)** | | 235.6 | | 0.3 | |
| IP (our dose) | 16.5 | 36 | | | 1.32 mg/kg (79.2 mg total) |

DAC has been approved by FDA for use in humans at 15 $mg/m^2$ and 20 $mg/m^2$ (8). It has been reported that 5 mg/kg in mice was equivalent the low dose,15 $mg/m^2$, i.e. the FDA approved dose used in human hematopoietic malignancies. Further, the 15 $mg/m^2$ dose was reported to achieve nanomolar plasma concentration in humans and mice. The dose conversion and calculation were based on the FDA recommendation (https://www.fda.gov/downloads/drugs/guidances/ucm078932.pdf) and the literature. Finally, it has been reported that plasma collected after 4 hr. i.v. infusion of DAC at total doses of 2 mg/kg in mice achieved equivalent nanomolar plasma concentration that reported in human. Similar to the reasoning we used for EGCG, because we were combining the compounds and administering over a longer period of time, we weighted our initial studies towards safety and chose a lower dose of DAC at 0.5 mg/kg i.p. (5-8).

#### DAC PK in mice

| DOSE | route of administration | Infusion rate | Infusion time | Cmax umol/L | Cmax ng/ml | HED (mg/kg) | HED ($mg/m^2$) |
|---|---|---|---|---|---|---|---|
| 5 mg /kg | IV infusion (samples collected at 4 h (steady state) * | 0.278mg/kg/h | 18 h | 0.350877 | 80 | 0.4 | 14.8 |
| 10 mg/kg | | 0.556 mg/kg/h | | 0.83333 | 190 | 0.8 | 29.6 |
| 20 mg/kg | | 1.11 mg/kg/h | | 2.675439 | 610 | 1.6 | 59.2 |
| 2 mg/kg | IV infusion (samples collected at 4 h (steady state) ** | 0.133 mg/kg/h | 15 h | 0.313158 | 71.4 | 0.16 | 5.92 |
| 1 mg/kg (male) | Oral (samples collected within in 1 hour) *** | | | 1.75 | | 0.08 | 2.96 |
| 1 mg/kg (female) | | | | 2.05 | | | |
| OUR DOSE 0.5 mg/kg | IP | | | | | 0.04 | 1.48 |

### DAC PK human

**Three Day Regimen:**

- Administer DAC at a dose of 15 mg/m2 by continuous intravenous infusion over 3 hours repeated every 8 hours for 3 days.
- Repeat cycles every 6 weeks upon hematologic recovery (ANC at least 1,000/μL and platelets at least 50,000/μL) for a minimum of 4 cycles.
- A complete or partial response may take longer than 4 cycles. Delay and reduce dose for hematologic toxicity.

**Five Day Regimen:**

- Administer DAC at a dose of 20 mg/m2 by continuous intravenous infusion over 1 hour daily for 5 days.
- Delay and reduce dose for hematologic toxicity.
- Repeat cycles every 4 weeks upon hematologic recovery (ANC at least 1,000/μL and platelets at least 50,000/μL) for a minimum of 4 cycles. A complete or partial response may take longer than 4 cycles.

https://www.accessdata.fda.gov/drugsatfda_docs/label/2018/021790s021lbl.pdf

| concentration in plasma (umol/L) | Human dose |
|---|---|
| 0.32 | 15 mg/m$^2$ 3-hr infusion every 8 hours for 3 days |
| 0.645 | 20 mg/m$^2$ 1-hr infusion daily for 5 days |

In human, The terminal elimination **half-life** ($t_{1/2}$) of **decitabine** is 37–47 minutes. Consequently, steady-state in plasma is reached at the end of the 3-hour infusion
