## Supplemental Tables for "CHA1: A New Combinatorial Therapy That Reciprocally Regulates Wnt and JAK/STAT/Interferon Signaling to Re-program Breast Tumors and the Tumor-Resident Landscape"

1 Supplemental Table I

**Wnt Target Genes**

| Gene ID | log2 Fold Change | P-value |
|---|---|---|
| MMP2 | -2.95 | 0.00003 |
| NTRK2 | -4.64 | 0.00005 |
| SMO | -3.61 | 0.00005 |
| TWIST1 | -3.24 | 0.0001 |
| TCF4 | -2.29 | 0.00005 |
| IGF1 | -3.31 | 0.00008 |
| ID2 | -1.78 | 0.00009 |
| CEBPD | -1.89 | 0.00015 |
| CTGF | -0.56 | 0.00115 |
| SFRP2 | -2.41 | 0.0013 |
| TGFB3 | -1.14 | 0.0014 |
| VEGFA | -0.64 | 0.00195 |
| DKK1 | -0.47 | 0.00655 |
| TCF7L1 | -1.00 | 0.01165 |
| TLE1 | -0.45 | 0.0119 |
| EGR1 | -0.49 | 0.0134 |
| EFNB1 | -0.51 | 0.02185 |

2 Supplemental Table II

**Antigen Presentation Pathway**

| Gene ID | Log2 Fold Change | P-value |
|---|---|---|
| CD74 | 1.646 | 5.00E-05 |
| HLA-B | 0.72 | 5.00E-05 |
| HLA-DMA | 0.894 | 0.0161 |
| HLA-DMB | 1.616 | 0.01965 |
| HLA-DPA1 | 1.527 | 0.00875 |
| HLA-DRA | 1.204 | 0.00865 |
| HLA-DRB1 | 1.31 | 0.01205 |
| HLA-DRB5 | 1.699 | 0.0725 |
| HLA-F | -2.036 | 0.0036 |
| HLA-G | -3.024 | 5.00E-05 |
| PSMB9 | 1.457 | 5.00E-05 |

3

**Graft-versus-Host Disease Signaling**

| Gene ID | Log2 Fold Change | P-value |
|---|---|---|
| FASLG | 3.6177 | 0.01905 |
| HLA-B | 0.72 | 5.00E-05 |
| HLA-DMA | 0.894 | 0.0161 |
| HLA-DMB | 1.616 | 0.01965 |
| HLA-DRA | 1.204 | 0.00865 |
| HLA-DRB1 | 1.31 | 0.01205 |
| HLA-DRB5 | 1.699 | 0.0725 |
| HLA-F | -2.036 | 0.0036 |
| HLA-G | -3.024 | 5.00E-05 |
| PRF1 | 3.68202 | 0.0038 |

Supplemental Table III

#### Supplemental Table IV

**4**

**Allograft Rejection Signaling**

| Gene ID | Log2 Fold Change | P-value |
|---|---|---|
| FASLG | 3.6177 | 0.01905 |
| HLA-B | 0.72 | 5.00E-05 |
| HLA-DMA | 0.894 | 0.0161 |
| HLA-DMB | 1.616 | 0.01965 |
| HLA-DPA1 | 1.527 | 0.00875 |
| HLA-DRA | 1.204 | 0.00865 |
| HLA-DRB1 | 1.31 | 0.01205 |
| HLA-DRB5 | 1.699 | 0.0725 |
| HLA-F | -2.036 | 0.0036 |
| HLA-G | -3.024 | 5.00E-05 |
| PRF1 | 3.68202 | 0.0038 |

**5**

**Autoimmune Thyroid Disease signaling**

| Gene ID | Log2 Fold Change | P-value |
|---|---|---|
| HLA-B | 0.72 | 5.00E-05 |
| HLA-DMA | 0.894 | 0.0161 |
| HLA-DMB | 1.616 | 0.0197 |
| HLA-DPA1 | 1.527 | 0.00875 |
| HLA-DRA | 1.204 | 0.00865 |
| HLA-DRB1 | 1.31 | 0.012 |
| HLA-DRB5 | 1.699 | 0.0725 |
| HLA-F | -2.036 | 0.0036 |
| HLA-G | -3.024 | 5.00E-05 |
| LAT | 1.362 | 0.0011 |
| TGFB3 | -1.143 | 0.0014 |
| TNFRSF11B | 0.892 | 0.0179 |
| TNFRSF1B | 0.964 | 0.0261 |

#### Supplemental Table V

**6**

**Th1 pathway**

| Gene ID | Log2 Fold Change | P-value |
|---|---|---|
| APH1B | 0.728 | 0.0049 |
| DLL1 | -5.148 | 0.00355 |
| GATA3 | -1.767 | 5.00E-05 |
| HLA-B | 0.72 | 5.00E-05 |
| HLA-DMA | 0.894 | 0.0161 |
| HLA-DMB | 1.616 | 0.0197 |
| HLA-DPA1 | 1.527 | 0.00875 |
| HLA-DRA | 1.204 | 0.00865 |
| HLA-DRB1 | 1.31 | 0.012 |
| HLA-DRB5 | 1.699 | 0.0725 |
| ICAM1 | 1.331 | 0.0013 |
| ICOSLG | 1.551 | 0.00745 |
| NFATC2 | -1.1 | 0.0394 |
| NOTCH3 | 1.329 | 0.00105 |

Supplemental Table VI

**7**

**Th2 pathway**

| Gene ID | Log2 Fold Change | P-value |
|---|---|---|
| APH1B | 0.728 | 0.0049 |
| BHLHE41 | -0.905 | 0.0007 |
| DLL1 | -5.148 | 0.00355 |
| GATA3 | -1.767 | 5.00E-05 |
| HLA-B | 0.72 | 5.00E-05 |
| HLA-DMA | 0.894 | 0.0161 |
| HLA-DMB | 1.616 | 0.0197 |
| HLA-DPA1 | 1.527 | 0.00875 |
| HLA-DRA | 1.204 | 0.00865 |
| HLA-DRB1 | 1.31 | 0.012 |
| HLA-DRB5 | 1.699 | 0.0725 |
| ICAM1 | 1.331 | 0.0013 |
| ICOSLG | 1.551 | 0.00745 |
| JAG2 | -1.323 | 0.00545 |
| MAF | -0.923 | 0.021 |
| NFATC2 | -1.1 | 0.0394 |
| NOTCH3 | 1.329 | 0.00105 |
| S1PR1 | -2.063 | 0.00045 |
| SPI1 | -0.89 | 0.0336 |

Supplemental Table VII

8

**Th1 and Th2 Activation Pathway**

| Gene ID | Log2 Fold Change | P-value |
|---|---|---|
| APH1B | 0.728 | 0.0049 |
| BHLHE41 | -0.905 | 0.0007 |
| DLL1 | -5.148 | 0.00355 |
| FGFR2 | -4.486 | 5.00E-05 |
| GATA3 | -1.767 | 5.00E-05 |
| HLA-B | 0.72 | 5.00E-05 |
| HLA-DMA | 0.894 | 0.0161 |
| HLA-DMB | 1.616 | 0.0197 |
| HLA-DPA1 | 1.527 | 0.00875 |
| HLA-DRA | 1.204 | 0.00865 |
| HLA-DRB1 | 1.31 | 0.012 |
| HLA-DRB5 | 1.699 | 0.0725 |
| ICAM1 | 1.331 | 0.0013 |
| ICOSLG | 1.551 | 0.00745 |
| JAG2 | -1.323 | 0.00545 |
| MAF | -0.923 | 0.021 |
| NFATC2 | -1.1 | 0.0394 |
| NOTCH3 | 1.329 | 0.00105 |
| S1PR1 | -2.063 | 0.00045 |
| SPI1 | -0.89 | 0.0336 |

Supplemental Table VIII

### Supplemental Table IX

| Diseases or Functions Annotation | p-value | Predicted Activation State | Activation z-score |
|---|---|---|---|
| Size of body | 3.2E-14 | Decreased | -5.463 |
| Development of neurons | 1.87E-16 | Decreased | -4.984 |
| Viral Infection | 1.17E-15 | Decreased | -4.768 |
| Organization of cytoskeleton | 7.31E-16 | Decreased | -4.346 |
| Organization of cytoplasm | 7.18E-14 | Decreased | -4.346 |
| Microtubule dynamics | 1.05E-15 | Decreased | -4.051 |
| Quantity of cells | 2.33E-24 | Decreased | -3.845 |
| Cell survival | 3.54E-17 | Decreased | -3.797 |
| Morphogenesis of neurons | 1.44E-11 | Decreased | -3.782 |
| Cell viability | 1.9E-15 | Decreased | -3.653 |
| Transcription of DNA | 5.71E-12 | Decreased | -3.606 |
| Formation of cellular protrusions | 8.23E-12 | Decreased | -3.572 |
| Vasculogenesis | 1E-21 | Decreased | -3.388 |
| Infection by RNA virus | 7.72E-12 | Decreased | -3.376 |
| Migration of endothelial cells | 6.67E-15 | Decreased | -3.2 |
| Development of vasculature | 4.72E-30 | Decreased | -3.108 |
| Angiogenesis | 1.18E-26 | Decreased | -3.1 |
| Migration of vascular cells | 3.72E-16 | Decreased | -3.092 |
| Differentiation of nervous system | 1.49E-14 | Decreased | -2.973 |
| Development of head | 6.04E-20 | Decreased | -2.882 |
| Growth of epithelial tissue | 4.72E-25 | Decreased | -2.866 |
| Development of body axis | 2.12E-22 | Decreased | -2.71 |
| Development of Visual system | 6.93E-11 | Decreased | -2.708 |
| Formation of eye | 1.91E-10 | Decreased | -2.708 |
| Synthesis of lipid | 5.04E-12 | Decreased | -2.688 |
| Quantity of neurons | 1.78E-12 | Decreased | -2.635 |
| Movement of vascular endothelial cells | 1.34E-11 | Decreased | -2.633 |
| Cell movement of endothelial cells | 1.32E-17 | Decreased | -2.622 |
| Proliferation of stem cells | 1.07E-12 | Decreased | -2.597 |
| Transcription of RNA | 1.4E-16 | Decreased | -2.589 |
| Growth of connective tissue | 9.67E-22 | Decreased | -2.436 |
| Proliferation of connective tissue cells | 2.21E-20 | Decreased | -2.357 |
| Development of sensory organ | 6.62E-14 | Decreased | -2.304 |
| Uptake of monosaccharide | 3.19E-11 | Decreased | -2.3 |
| Differentiation of neurons | 4.21E-11 | Decreased | -2.296 |
| Proliferation of neuronal cells | 9.27E-13 | Decreased | -2.285 |
| Quantity of connective tissue | 4.14E-21 | Decreased | -2.284 |
| Formation of lung | 1.72E-11 | Decreased | -2.266 |
| Quantity of metal ion | 9.31E-12 | Decreased | -2.234 |
| Proliferation of epithelial cells | 4.17E-21 | Decreased | -2.198 |
| Growth of embryo | 1.66E-11 | Decreased | -2.179 |
| Formation of cytoskeleton | 2.39E-14 | Decreased | -2.176 |
| Quantity of metal | 4.72E-11 | Decreased | -2.161 |
| Development of epithelial tissue | 1.34E-15 | Decreased | -2.155 |

Supplemental Table IX (Continued)

| | | | |
|---|---|---|---|
| Transcription | 4.53E-18 | Decreased | -2.148 |
| Quantity of lymphatic system cells | 2.65E-13 | Decreased | -2.146 |
| Cell movement of connective tissue cells | 9.63E-11 | Decreased | -2.058 |
| Migration of muscle cells | 6.35E-13 | Decreased | -2.054 |
| Growth of skin | 1.73E-15 | Decreased | -2.014 |
| Proliferation of dermal cells | 7.53E-14 | Decreased | -2.004 |
| Development of cardiovascular tissue | 2.6E-11 | Decreased | -2.001 |
| Development of endothelial tissue | 5.4E-11 | Decreased | -2.001 |
| Advanced malignant solid tumor | 2.17E-20 | Increased | 2 |
| Metastatic solid tumor | 5.95E-20 | Increased | 2 |
| Left ventricular dysfunction | 1.31E-11 | Increased | 2.045 |
| Extraintestinal functional disorder | 2.03E-10 | Increased | 2.123 |
| Blood pressure | 2.03E-13 | Increased | 2.125 |
| Visceral metastasis | 1.18E-17 | Increased | 2.195 |
| Dystrophy of muscle | 2.52E-12 | Increased | 2.219 |
| Structural renal abnormality | 4.39E-12 | Increased | 2.378 |
| Cell death of renal tubule | 3.57E-11 | Increased | 2.409 |
| Apoptosis of tumor cell lines | 1.98E-18 | Increased | 2.482 |
| Necrosis of epithelial tissue | 1.16E-12 | Increased | 2.575 |
| Diabetes mellitus | 3.12E-21 | Increased | 2.72 |
| Apoptosis of renal tubule | 1.59E-10 | Increased | 2.749 |
| Necrosis | 5.68E-34 | Increased | 2.785 |
| Progressive neurological disorder | 1.32E-11 | Increased | 2.795 |
| Neuromuscular disease | 8.17E-11 | Increased | 2.937 |
| Glucose metabolism disorder | 1.05E-27 | Increased | 3.626 |
| Apoptosis | 7.25E-27 | Increased | 3.666 |
| Aplasia or hypoplasia | 2.01E-10 | Increased | 4.696 |
| Neonatal death | 8.6E-11 | Increased | 6.172 |
| Perinatal death | 4.67E-13 | Increased | 6.664 |
| Morbidity or mortality | 1.07E-32 | Increased | 8.287 |
| Organismal death | 3.69E-33 | Increased | 8.356 |

### Supplemental Table X

**S10 Table: Human primer sequences**

| | Forward | Reverse | Temp |
|---|---|---|---|
| 18S | GCCCGAAGCGTTTACTTTG | CTTAATCATGGCCTCAGTTCC | any |
| AXIN2 | GTGTGAGGTCCACGGAAACT | GAATCATCCGTCAGCGCATC | 61 |
| CD74 | CAGCGCGACCTTATCTCCAA | GGTACAGGAAGTAGGCGGTG | 62.5 |
| CTLA-4 | TAATTGATCCAGAACCGTGCC | CTCTGTTGGGGGCATTTTCAC | 56.7 |
| DDX58 | CTGGACCCTACCTACATCCTG | GGCATCCAAAAAGCCACGG | 61.2 |
| DHX58 | GGGCCTCCAAACTCGATGG | TTCTGGGGTGACATGATGCAC | 57 |
| ERV3-1 | AGCCGGAGCTTCTGGTGTAG | AGTGGGTCCTGGCGTCTTA | 55.3 |
| FASL | TGGAAATAAACTGCACCCGGAC | CTCTTTGCACTTGGTGTTGCTGG | 63 |
| HBP1 | GGCGACGGGTTTGTC AGAG | TGCCAGATTGGGTAGGATCAC | 61.3 |
| HLA-B | CTAGCAGTTGTGGTCATCGGAG | GGAGGCGTGAAGAAATCCTG | 61.5 |
| HLA-DRB5 | AGCATGGTGTGTCTGAAGC | CCCGTTGAAGAAATGACACTCA | 59.7 |
| ICAM1 | CTATAAAGGATCACGCGCCC | GGCAGGATGACTTTTGAGGG | 62 |
| IFI27 | TTCTTTGGGTCTGGCTCAAGT | GCCACAACTCCTCCAATCACA | 59.5 |
| IFIH1 (MDA-5) | GGAGTCAAAGCCCACCATCT | GGTGACGAGACCATAACGGA | 54.7 |
| IFIT1 | AAGCAGGACCCACAAGAATGT | GGCATTTCATCGTCATCAATGG | 58 |
| IFIT3 | GAACATGCTCAAGCAGA | CAGTTGTGTCCACCCTTCCT | 54.7 |
| IFTI2 | AACAAAAAGGAACCAGAGGCCA | TAGTTGCCGTAGGCTGCTCTC | 61.2 |
| ISG20 | GAAGCACGACTTCCAGGCAC | TCTTCCACCGAGCTGTGTCC | 61.2 |
| MAGE-A3 | CCCTGAGCAACGAGCGAC | GACTCTGGTCAGGGCAACAG | 61.6 |
| MAGE-A6 | CCCTGAGCAACGAGCGA | ACTCTGGTCAGGGCAACAG | 59.5 |
| NY-ESO-1 | TGTCCGGCAACATACTGACT | AAAAACACGGGCAGAAAGCAC | 60 |
| OAS1 | TCCGTGAAGTTTGAGGTCCAG | AGGTTTATAGCCGCCAGTCA | 59.5/61.3 |
| OAS2 | GCTCAGAAGCTGGGTTGGTTT | GGATGTCACGTTGGCTTCTCT | 58 |
| OAS3 | CGTCAAACCCAAGCCACAAG | TTCCACAACCCCTGTAGGCA | 60.6 |
| OASL | GAAGGAAGAGGTGCTAGACGC | GAAACAGCTCAGAAACGCCAC | 60.6 |
| PD-L1 | TATGCCTTGGTGTAGCACTGA | CCGATGAACCCCTAAACCACA | 58.6 |
| PD-L2 | GAACCCAGGACCCATCCAAC | CCCAAGACCACAGGTTCAGA | 57.4 |
| PRF1 | CACCAGGACCAGTACAGCTT | ATGAAGTGGGTGCCGTAGTT | 61.6 |
| PSMB9 | GGGGCGTTGTGATGGGTTC | GCAGCCAAAACAAGTGGAGGT | 60.6 |
| sFRP1 | CTGATAACTGGTTGCTGTGTC | CATCCATGTCCTGTGTATCTGC | 61.5 |

### Supplemental Table XI

**S11 Table: Mouse primer sequences**

| | Forward | Reverse | Temp |
|---|---|---|---|
| Axin2 | GACAGTAGCGTAGATGGAGTCC | CGGCTTTCCAGCTCCAGTTT | 62.4 |
| b2m | TCGCTTCAGTCGTCAGCAT | GCAGTTCAGTATGTTCGGCT | 55.8/59.7 |
| Ccl4 | TCTCTCCTCTTGCTCGTGGC | ATCTGTCTGCCTCTTTTGGTCA | 58.8 |
| Ccl5 | TGCTGCTTTGCCTACCTCTC | TTGAACCCACTTCTTCTCTGGG | 55.9 |
| Cd4 | TGGGGAAGGAAGGGGAATCA | AACCATCAAACTGCGAAGGC | 54.9/55.4 |
| Cd8a | CGGATTGGACTTCGCCTGT | TGCAAACACGCTTTCGGCT | 58.8 |
| Ctla-4 | TCCAAGGACTGAGAGCTGTT | GGGCATTTTCACATAGACCCC | 54.5 |
| Cxcl10 | TGAGAGACATCCCGAGCCAA | CCCTATGGCCCTCATTCTCAC | 59.9 |
| Cxcl9 | CTCGGACTTCACTCCAACACA | GTAGTGGATCGTGCCTCGG | 60.6 |
| Fasl | TGAGTTCACCAACCAAAGCCT | TCACTCCAGAGATCAGAGCGG | 55.9 |
| Gzmb | TGTGTGCTATGTGGCTGGTT | CCCGAAAGGAAGCACGTTTG | 55.4 |
| H2Aa | TGGAGGTGAAGACGACATTGA | AGCTATGTTTTGCAGTCCACC | 57.4 |
| H2Ab1 | CCTGTGCCTTAGAGATGGCT | GTTGGTGAAGTAGCACTCGC | 55.1 |
| H2d1 | AAGTGGGAGCAGAGTGGTG | ATGTCAGCAGGGTAGAAGCC | 55.8 |
| H2k1 | ATCGCCCTGAACGAAGACC | CACATGGGCCTTTGGGGAA | 54 |
| Hbp1 | GGAAGACTTTGCTAGAGCCG | CAGTGAGCAAGCCATCTTCT | 57 |
| Ido1 | GCTTTGCTCTACCACATCCAC | TGTCCTCTCAGTCCGTCCG | 57.4 |
| Ifi27 | TGGAGAGAGCTGGGGAAATCG | AAAACGCCATGAGGACCAGT | 55.8 |
| Ifit1 | GCTACCACCTTTACAGCAACC | CCTTTCAGGTGCCTCACGTA | 61.8 |
| Ifit3 | CTCAGCCCACACCCAGCTTTT | GTTGCACACCCTGTCTTCCATA | 55.7 |
| Ifng | ATGAACGCTACACACTGCATC | CCATCCTTTTGCCAGTTCCTC | 57.1 |
| Lag3 | CTGTCACGTTGGCGGTCATC | GGACAGATTGTTCAGGGGACG | 60.2 |
| Mage-a3 | AAGAGAGTGCTCAGGCCCAA | TGCCACACAGGGAGAGTAGG | 58.7/59.7 |
| MuERV-L | TGGTGGTCGAGATGGAGGTTA | CCGTGAATGGTGGTTTTAGCA | 55.3/59.5 |
| Oas1a | TGTACGAGGTTCAGCATGAGAG | GGCTTGCTGGAAGTATTAACATGA | 58.6 |
| Oas1b | TCTGCTTTATGGGGCTTCGG | TCGACTCCCATACTCCCAGG | 59.6 |
| Oas1g | TTGAGGTCCAGAGTTCATGGTG | CAGATGAGGATGGTGTAGATTAAGG | 57.9 |
| Oas3 | TGGGTCACAAGTCTGCCTTC | ACTTGAGTCAGGAAGTGGCG | 57.3 /59.6 |
| Oasl1 | AGACCTGGGAGACCATCACT | TTTGACCAACCGAAGGAGGC | 63.6 |
| Oasl2 | GAAGAACCTCCTCCGGTTGG | CACGGCTACGAACCCTTCAT | 58.6/59.6 |
| Pd-1 | GGAGACTGCTACTGAAGGCG | GTGAAGGTGGCATTTGCTCC | 55.4 |
| Pd-l1 | GGACTACAAGCGAATCACGC | TTCTCTTCCCACTCACGGGT | 55.4 |
| Pd-l2 | TCGGTGTGATTGGCTTCCAG | TCTTTAGGGGCTGTCACGGT | 55.4 |
| Prf1 | CCCACTCCAAGGTAGCCAAT | GAGCTGTTAAAGTTGCGGGG | 56.4 |
| Psmb10 | GAATGCGTCCTTGGAACACG | GGGGCGATGAAGTGGATCTT | 59.6/60.2 |
| Sfrp1 | ACTGGCCCGAGATGCTCAAA | CATCCTCAGTGCAAACTCGCT | 62 |
| Stat1 | CTGTCATCCCGCAGAGAGAA | CTGCTGAAGCTCGAACCACT | 60.2 |

Supplemental Table XI (Continued B)

| | | | |
|---|---|---|---|
| Tigit | TGAGCCAGTTTCAGTTGGAGG | TATCTATCGTGCCTGCTGTGG | 61 |

**S12 Table: Primary antibodies used for western blotting**

| Reagent | Source |
|---|---|
| β-actin | Sigma-Aldrich |
| β-catenin | Millipore |
| E-cadherin | Cell signaling |
| pSTAT-3$^{Y705}$ | Cell signaling |
| STAT-3 | Cell signaling |

Supplemental Table XII
